# Evidence for a travel direction signal in humans that is independent of head direction

**DOI:** 10.1101/2022.08.22.504860

**Authors:** You Cheng, Sam Ling, Chantal E. Stern, Andrew Huang, Elizabeth R. Chrastil

**Author notes:** **For correspondence:** (ERC). Department of Neurology, Harvard Medical School, Massachussets General Hospital, Charlestown, MA, USA.

## Abstract

We often assume that travel direction is redundant with head direction, but from first principles these two factors provide differing spatial information. Although head direction has been found to be a fundamental component of human navigation, it is unclear how self-motion signals for travel direction contribute to forming a travel trajectory. Employing a novel motion adaptation paradigm from visual neuroscience designed to preclude a contribution of head direction, we found high-level aftereffects of perceived travel direction, indicating that travel direction is a fundamental component of human navigation. Interestingly, we discovered a higher frequency of reporting perceived travel toward the adapted direction compared to a no-adapt control – an aftereffect that runs contrary to low-level motion aftereffects. This travel aftereffect was maintained after controlling for possible response biases and approaching effects, and it scaled with adaptation duration. These findings demonstrate the first evidence of how a pure travel direction signal might be represented in humans, independent of head direction.

## Introduction

In daily navigation, for activities as simple as going to a grocery store, we form a trajectory of our travel paths. How do we use self-motion information to form this travel trajectory? Travel trajectory is derived from time and velocity. Velocity in turn is composed of speed and travel direction, which is the direction of one’s body movement. Although *head direction* – the direction that one’s head is facing toward (also called facing direction) – is typically assumed to be the direction of travel, in reality these two factors offer different spatial information. For instance, when we walk on the street, we constantly look around at our surroundings, changing our head direction while maintaining a constant travel direction. Thus, from first principles *travel direction*, rather than head direction or facing direction, is the most important component in forming a travel trajectory, as well as in maintaining spatial-vector memory over time (***Stone et al., 2017; Hulse et al., 2021***).

Only a handful of human and rodent studies have examined whether travel direction is coded separately from heading direction. Although they do not offer the same amount of information, spatially-tuned head direction cells have previously been used as a theoretical basis for the formation of a travel trajectory. Indeed, head direction cells have been discovered in the rat brain, which selectively fire in the direction a rat is facing towards, independent of its location (***Chen et al., 1994; Ranck Jr, 1984; Taube et al., 1990; Taube, 1995***), demonstrating that head direction is a fundamental component of the internal orientation system of navigation. These head direction cells are found in thalamus (***Taube, 1995***), retrosplenial cortex (***Chen et al., 1994; Cho and Sharp, 2001***), presubiculum (***Ranck Jr, 1984***), extrastriate cortex (***Chen et al., 1994***), and entorhinal cortex (***Frank et al., 2000; Quirk et al., 1992***). Head direction cells could fire independently of whether an animal is moving or motionless (***Taube, 1998***), whereas travel direction should involve motion, therefore it might be more difficult to record activity related to travel direction. One study that contrasted head and travel direction found that head direction is coded more strongly than travel direction in a population of rodent entorhinal neurons (***Raudies et al., 2015***). Further experimental findings in rodents and *Drosophila* also suggest the existence of neural signals for travel direction (***Lyu et al., 2021; Olson et al., 2017***).Taken together, while there is evidence to suggest a dissociation between coding of head direction and travel direction, the role that travel direction plays in the internal representation system of human navigation remains unclear.

The goal of this study was to determine whether there are behavioral signatures of travel direction as a fundamental component of human spatial navigation, independent of facing direction. To test the role of travel direction in human navigation, we utilized a motion adaptation paradigm adopted from vision science. In motion adaptation, neurons selective for visual motion features (e.g., moving downward) will adapt to visual stimuli that corresponds to its selective firing properties after prolonged exposure to the same stimuli (***Barlow, 1990***). This adaptation often results in a decrease in the neural response to the same stimuli, compared to an unadapted stimulus (***Brown and Masland, 2001; Lisberger and Movshon, 1999; Maffei et al., 1973; Miller et al., 1991***). This neuronal change is represented at the behavioral level as a motion aftereffect (MAE) – a visual phenomenon produced after motion adaptation such that a stationary stimulus will appear to move in the opposite direction of the previously viewed motion. These motion aftereffects are suggested to be associated with an amalgam of adaptation of motion-selective opponency cells at several visuocortical sites (***Antal et al., 2004; Ashida and Osaka, 1994; Bach and Ullrich, 1994; Barlow and Hill, 1963; Bex et al., 1999; Culham et al., 1998; Kohn and Movshon, 2003; Mather et al., 2008; Sutherland, 1961; Verstraten et al., 1998***). Thus, our study operates under the assumption that if travel direction exhibits adaptation-like effects, then it is a fundamental component of the representation system for human navigation.

In the present study, participants were adapted to travel direction by viewing movement in a hallway in a constant direction. To dissociate travel direction from head direction, head direction randomly reversed while travel direction was kept constant during adaptation (Fig. 1 and Video 1). We expected to observe motion aftereffects compared to a control condition with no adaptation. Typically, motion aftereffects are found in the opposite direction to the adapted motion (***Anstis et al., 1998; Mather et al., 1998***), and so we predicted motion aftereffects in the opposite direction of travel. However, high-level motion aftereffects are frequently seen in the same direction as the locally adapted motion because they use non-retinotopic visual features, although they might go in the opposite direction from the globally perceived movement (***Culham et al., 2000; Dubé and Von Grünau, 1992; Hiris and Blake, 1992; Nishida and Sato, 1995; Von Grünau, 1986***). Motion after-effects in other sensory modalities have been reported to go in the same direction as travel. For example, in a podokinetic (walking-based) adaptation study, blindfolded people were adapted to a circular trajectory but perceived themselves to be going straight; when released to move freely, they formed the same circular trajectory (***Earhart et al., 2001***). This occurred both when walking forward or backward, indicating that the motion adaptation could be transferred to both directions of locomotion. Navigation is considered high-level cognition (***Wolbers and Hegarty, 2010***) whose information processing generally centers around higher level brain regions rather than visual areas (***Chrastil, 2013; O’Keefe and Nadel, 1978***), and navigation process are typically non-retinotopic (***Chrastil et al., 2019; Giudice, 2018; Loomis et al., 1993***). Thus, we theorized that we could instead observe high-level motion aftereffects in the same direction as the motion adaptation.

**Figure 1.**
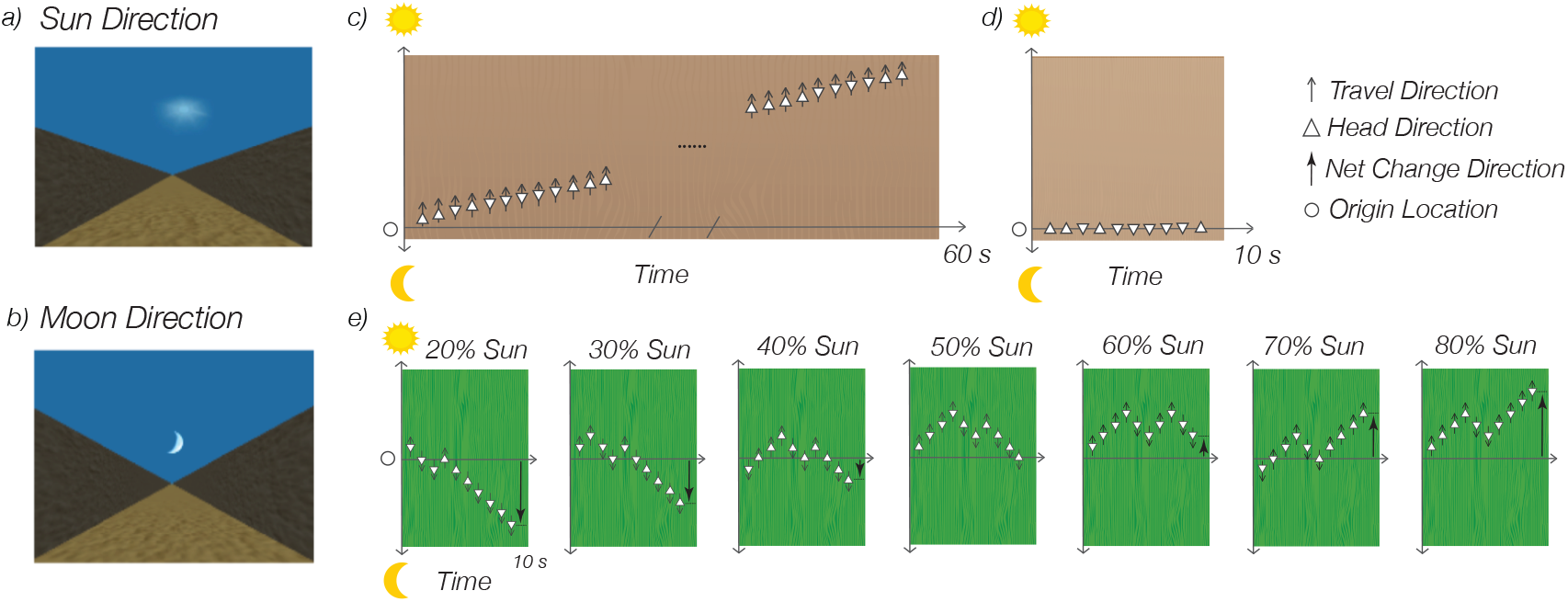
Experiment 1. Hallway during the adaptation phase, facing the a) sun or b) moon direction. Note that in the virtual environment, both the sun and the moon were rendered to move with the viewer at a constant distance; the size of the moon and the sun did not change with self-motion and participants could not evaluate distance change based on perceptual changes in either the sun or the moon. The extreme length of the hallway and random textures also precluded using changes in the hallway itself for location cues. The ground for the hallways turned green during the test phase to provide a visual cue for when to start tracking movement direction. c) Overhead illustration of the 60-second adaptation phase for the sun group. During the adaptation phase, visual movement traveled toward the sun while the facing direction occasionally changed. Half the participants were adapted to a similar moon condition, with travel direction toward the moon. d) Overhead illustration of the 10-second initial phase for the control session, which was the same for both the sun and moon groups. There was no visual travel, but the facing direction randomly changed. e) The test phase, which was the same for all sessions in all conditions. Visual movement traveled back-and-forth between the sun and the moon during a 10-second interval. Participants were asked to decide whether the total movement was more toward the sun or more toward the moon in that interval. The facing direction randomly changed during the test phase. Here, we show one example from each of the seven test phase conditions of the percent of net movement toward the adaptation direction (20%, 30%, 40%, 50%, 60%, 70%, 80%). See also the experiment videos 1.

In addition, as far as we know, motion adaptation paradigms have only been used previously to study motion from the third person view. The present study is the first to utilize such a paradigm to study a first-person view of motion - self-motion. This novel implementation of the motion adaptation paradigm may also lead to different adaptation effects than observed in previous visual perception studies. Regardless of the direction of the effect, a motion aftereffect would be the behavioral signature of travel direction selectivity that is represented in a particular way in the human brain, potentially operating as a fundamental basis function in human navigation.

## Results

### High-level motion aftereffects of travel direction

To test whether there is a travel direction signal in humans, we used a visual motion adaptation paradigm in desktop virtual reality (VR), designed to isolate travel direction from the contribution of head direction. In the initial adaptation phase, participants experienced 60 seconds of visual self-motion toward a cardinal direction (towards a sun or towards a moon, Fig. 1a,b) along a virtual hallway (Fig. 1c). In the test phase, participants then experienced a series of visual back and forth movements, toward and away from the initial cardinal direction (Fig. 1e). We then asked participants to report their net travel direction during the test phase. Critically, in both adaptation and test phases the heading direction alternated occasionally, cancelling out any effect of overall heading direction, to dissociate heading from travel direction. To maintain an adaptation state, participants experienced shorter 10-second “top-up” adaptation between each trial.

We compared the adaptation condition to a no-adapt control condition, in which participants viewed a static hallway with occasional heading changes (Fig. 1d); the control condition had the exact same test phase as the adaptation condition. By parametrically manipulating the coherence of global travel direction (Fig. 1e), we acquired psychometric functions for perceived travel direction, which allowed us to assess whether travel direction adaptation shifted the psychometric function, relative to the control.

In Experiment 1, we recruited 60 participants (31 females). To guard against response bias for one of the cardinal directions (i.e., a preference for selecting the sun or the moon), we divided the participants into two groups through random assignment, with each group adapting to either the sun or the moon direction. (Fig. 1 a - e; Methods Experiment 1). The groups were combined for analysis (see Supplemental Information Fig. 7 for analysis of each group separately).

We observed significant motion aftereffects of travel direction in the adaptation condition compared to the control (*F* (1, 59) = 11.38, *p* = 0.001, 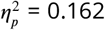, 95% confidence interval (*CI*) = [0.028, 0.332]) (Fig. 2). Although we found a significant shift in the function with adaptation, the pattern was not in the direction we initially predicted: motion adaptation increased travel estimation in the same direction as the adaptation, instead of producing a traditional opponent-process aftereffect in the opposite direction. As expected, there was a main effect of the actual percentage of motion on perceived direction (*F* (2.14, 126.49) = 522.34, *p <* 0.001, 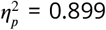, 95%*C.I*. = [0.882, 0.912]), with the perceived percentage of movement in the adaptation direction increased with the actual percentage. There was also an interaction between the experimental condition and the actual percentage of movement (*F* (4.22, 248.94) = 4.91, *p <* 0.001, 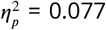, 95%*C.I*. = [0.021, 0.122]), which indicates that the difference between the adaptation and control conditions depended on the actual percentage of movement in the adaptation direction. Post-hoc analyses revealed that the adaptation significantly increased the perceived percentage of movement in the adaptation direction in several conditions where the actual percentage was below 60%. These planned comparisons further confirmed that the adaptation condition was increased toward the adaptation direction compared to the control condition (Fig. 2).

**Figure 2.**
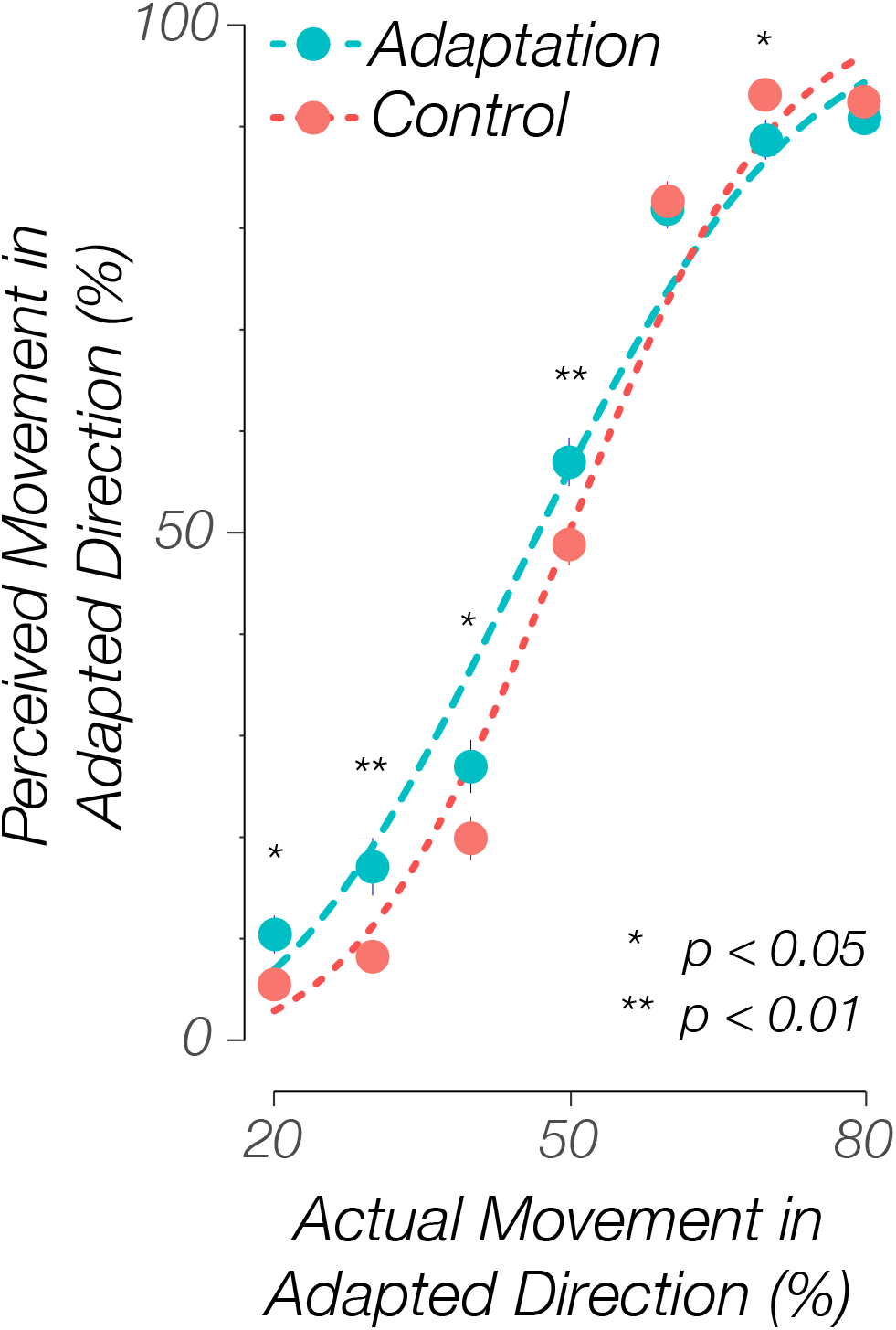
Experiment 1. The perceived percentage of movement in the adaptation direction compared with the actual percentage for all subjects (n = 60). Solid dots indicate the grand average value, and error bars indicate standard errors. Dashed lines indicate the average psychometric Weibull functions. The adaptation condition had an overall significantly higher reported percentage than the control condition (*p* = 0.001). The adaptation condition showed significantly higher reported percentages than corresponding control conditions at 20%, 30%, 40%, and 50%, supporting the aftereffect. This result suggests an aftereffect in the same direction of travel. There was also a significant interaction between condition and the actual percentage. The bias psychometric function (i.e., *α*) for the adaptation condition did not significantly shift compared to the control (*p* = 0.252). The uncertainty psychometric function (i.e., *β*) became more flattened when adapted, indicating that observers’ detectability of the difference between the two directions was decreased by adaptation (*p* = 0.007). * p *<* 0.05; ** p *<* 0.01, Tukey correction.

We conducted several additional analyses to preclude possible explanations besides a motion aftereffect. We separately analyzed results from the sun and moon adaptation groups and found the same result as the combined data (with somewhat weaker effects for the moon group), which ensured that neither the particular adaptation direction nor response bias disproportionately affected the results (Supplemental Information Fig. 7). We also found no serial position effects (i.e., no primacy or recency effects), suggesting that no particular portion of the test phase disproportionately contributed to the reported rate (e.g. participants only paid attention to the last second of the movement) (Supplemental Information Fig. 8 - 11). We found no reaction time differences between conditions. We only found the expected higher reaction time at the 50%, indicating that participants found the condition with equal amounts of travel in each direction difficult to judge, which is an indication that our task was effective (Supplemental Information Fig. 6). We also specifically analyzed just the first test trials in each block after the initial 60-second adaptation and found similar patterns of results as the entire data set; this finding suggests that the opposite motion aftereffect was not due to insufficient adaptation during the top-ups (Supplemental Information Fig. 12). Further, the test trials were the same for the control and test conditions, so it is unlikely that remembering the movement direction they experienced for more or less time (availability heuristic) would underlie the aftereffect. In addition, the pattern of results was stable after we controlled for strategies in the analyses (see Methods Experiment 1 Data Analysis; Supplemental Information Experiment1). Finally, we filtered out 9 subjects (around 15% of the subjects; 3 from the sun adaptation group, 6 from the moon adaptation group), based on a Weibull function (***Mood et al., 1974***) and the results from the remaining subjects still revealed the same pattern as the raw data. This finding suggests that the observed opposite motion aftereffect was not due to outlier subjects (see Methods Experiment 1 Data Analysis for details of the filtering procedure; Supplemental Information Fig. 13).

### Higher uncertainty in estimating travel directions after adaptation

To quantify the magnitude of the aftereffect, we fitted each participant’s data with a Weibull function (***Mood et al., 1974***). *α* and *β* values were derived for each fitted function: *α* means a bias to respond to the “sun” or the “moon” direction in the task, while *β* means the detectability of the difference in the task (Methods). We found no difference in the bias (*α*) between the adaptation and the control conditions ((59) = 1.16, *p* = 0.252, *C* = 0.196, 95%*C.I*. = [−0.143, 0.536],). The detectability (*β*) in the adaptation condition was significantly lower than in the control condition ((59) = −2.79, *p* = 0.007, *Cohen’ s d* = −0.439, 95%*C.I*. = [−0.765, −0.113]), indicating that people had more uncertainty in making judgments in the adaptation condition (Fig. 2).

In summary, our first experiment found a significantly increased perception of movement toward the direction of adaptation, consistent with a motion aftereffect. Participants also exhibited more uncertainty after the adaptation, compared to control, suggesting that the adaptation was affecting their judgments of movement direction during the test periods. Together, these findings are consistent with a role for travel direction that is independent of head direction.

### Motion aftereffects remain when adaptation heading is orthogonal to travel direction

In Experiment 2 (*n* = 30 participants; 16 females), we attempted to address additional questions about response biases and approaching effects from Experiment 1. Specifically, we wondered whether in Experiment 1 people felt like they were approaching the adapted direction due to the alignment of head and travel directions along the same axis. Although we separated travel direction from head direction by randomly flipping head direction throughout the task, travel direction was still on the same axis as head direction – the direction that aligns with the front-back body axis. One possible outcome of this alignment of travel direction and head direction is that people might more easily generate a feeling of approaching the adapted travel direction. Thus, to test this alternative possibility and to preclude approaching effects during the adaptation, in Experiment 2 we changed the facing direction to be perpendicular to travel direction (Fig. 3a-e and Video 2; Methods Experiment 2). Because travel direction and facing direction were never aligned, this experiment provides an even more stringent test of our motion aftereffect hypothesis.

**Figure 3.**
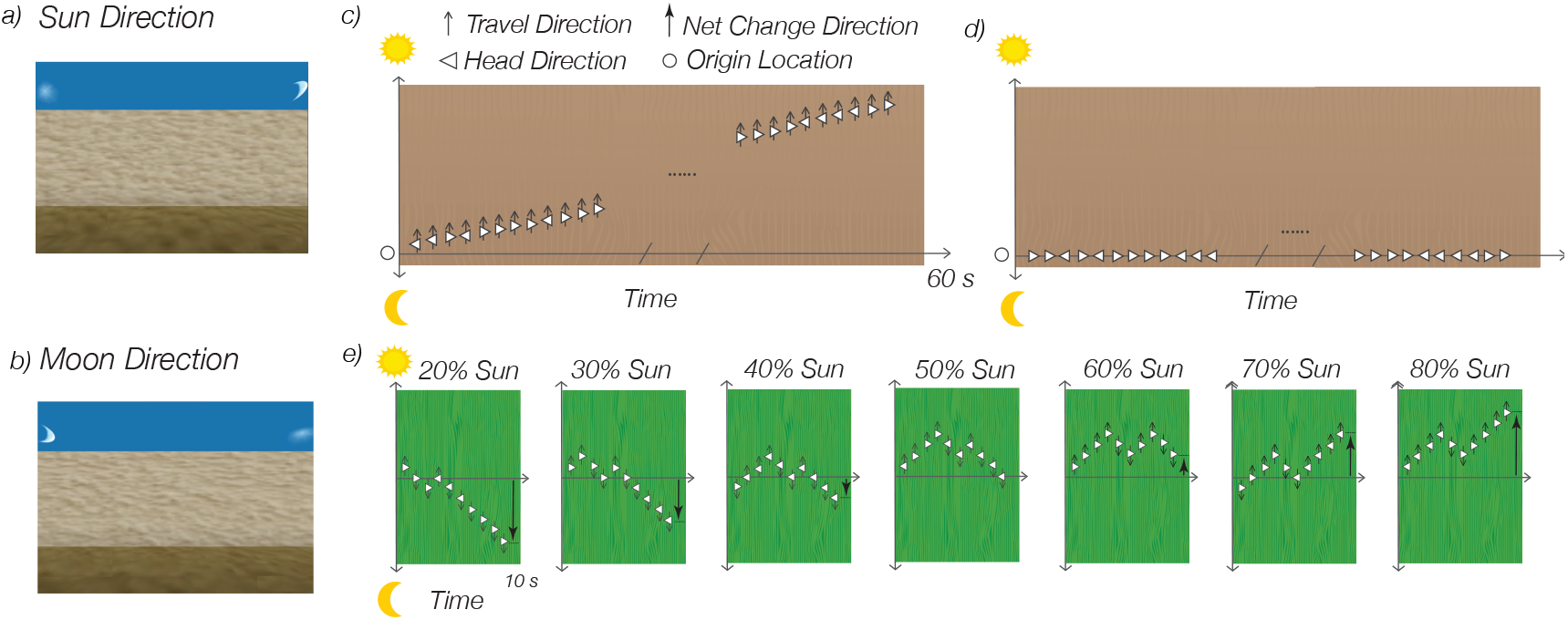
Experiment 2. a) Hallway during the adaptation phase, adapting to the sun direction. b) Hallway during the adaptation phase, adapting to the sun direction, but with the opposite facing direction compared to a). The ground for the hallways turned green during the test phase to provide a visual cue for when to start tracking movement direction. c) Overhead illustration of the 60-second adaptation phase for the experimental condition. During the adaptation phase, visual movement traveled toward the sun while the facing direction occasionally changed. d) Overhead illustration of the 60-second initial phase for the control session. There was no visual travel, but the facing direction randomly changed. e) The test phase, which was the same for all sessions in all conditions. Visual movement traveled back-and-forth between the sun and the moon during a 10-second interval. Participants were asked to decide whether the total movement was more toward the sun or more toward the moon in that interval. The facing direction randomly changed during the test phase. Here we show one example from each of the seven test phase conditions of the percent of net movement toward the adaptation direction (20%, 30%, 40%, 50%, 60%, 70%, 80%). See also the experiment video 2.

We observed a trend for a difference between the perceived percentage in the adaptation and the control conditions, but it was not statistically significant (*F* (1, 29) = 2.92, *p* = 0.098, 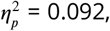, 95%*C.I*. = [0.000, 0.321],). As expected, there was a main effect of the actual percentage report (*F* (2.10, 60.89) = 232.24, *p <* 0.001, 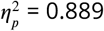, 95%*C.I*. = [0.861, 0.909]) that showed the perceived percentage increased with the actual percentage. There was no interaction between the experimental condition and the actual percentage (*F* (3.31, 95.97) = 2.07, *p* = 0.103, 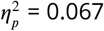, 95%*C.I*. = [0.000, 0.122],). However, for the planned Tukey HSD paired t-tests between adaptation trials and control trials within each actual percentage, we found that adaptation increased the perceived percentage where the actual percentages were 20% and 40%. These planned comparisons found that the adaptation condition was increased toward the adaptation direction, compared to the control condition (Fig. 4), consistent with Experiment 1.

**Figure 4.**
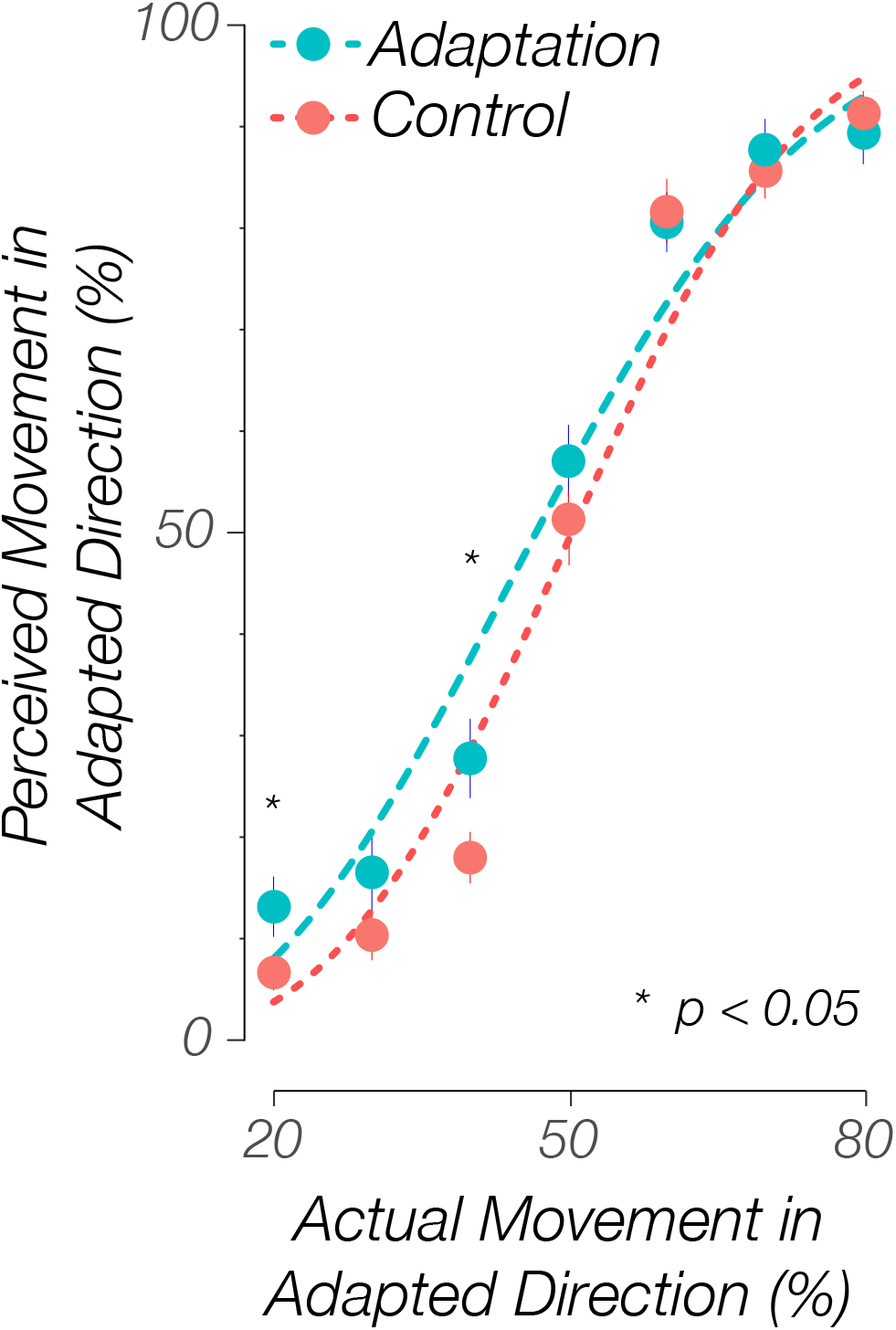
Experiment 2. The perceived percentage of movement in the adaptation direction compared with the actual percentage for all subjects (n = 30). Solid dots indicate the grand average value, and error bars indicate standard errors. Dashed lines indicate the average psychometric Weibull functions. The adaptation condition showed significantly higher reported percentages than corresponding control conditions at 20% and 40%, supporting the aftereffect. This result suggests an aftereffect in the same direction of travel. The bias psychometric function (i.e., *a*) did not significantly shift when adapted (*p* = 0.670). The uncertainty psychometric function (i.e., *β*) indicates that observers’ detectability of the difference between the two directions was not significantly decreased by adaptation (*p* = 0.210). * p *<* 0.05, Tukey correction.

Furthermore, after controlling for people’s self-reported strategies by including types of strategies (e.g., counting, focusing on part of the environment, and others) as a factor in the analyses (see Methods Experiment 2 Data Analysis), the difference between the adaptation and the control condition became more distinguishable, such that adaptation increased people’s reported rate (*F* (1, 27) = 5.58, *p* = 0.026, 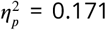, 95%*C.I*. = [0.000, 0.418]). There was also a significant interaction between condition and the actual percentage (*F* (3.53, 95.40) = 3.23, *p* = 0.020, 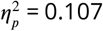, 95%*C.I*. = [0.012, 0.178]). Tukey HSD analyses revealed that people reported significantly more travel toward the adaptation direction in the adaptation session than in the control session at 20% (*p* = 0.003), 40% (*p* = 0.005), 50% (*p* = 0.031), and marginally more at 30% (*p* = 0.070) (see Supplemental Information Experiment 2 for more details). The motion aftereffect also got stronger after excluding 7 subjects (about 25% of subjects) based on implausible parameter estimates from the Weibull function fitting procedure (Supplemental Information Fig.15; see Methods Experiment 2 Data Analysis for details of the filtering procedure).

We fit each subject’s data into the Weibull function as we did in Experiment 1. We found no difference between the adaptation and the control conditions in either response bias (*α*) (*t* (29) = 0.43, *p* = 0.670, *Cohen’ s d* = 0.112, 95%*C.I*. = [−0.409, 0.633], *ns*) or detectability (β) (*t* (29) = −1.28, *p* = 0.210, *Cohen’ s d* = −0.251, 95%*C.I*. = [−0.648, −0.147], *ns*) (Fig. 4).

Thus, the observed aftereffects in Experiment 2 were weaker compared with those observed in Experiment 1, but still largely followed the pattern of a motion aftereffect in the direction of travel. Experiment 2 had fewer participants than Experiment 1; the sample size was based on our power analysis from the Experiment 1 results. This weaker aftereffect could be also due to the unusual travel direction in the task: in daily life, people experience walking forward or backward more often than walking laterally. Moreover, especially for the adaptation condition, tracking four directions (front-back for head direction and left-right or sun-moon directions for travel direction) in Experiment 2 was likely more challenging than in Experiment 1, where people only tracked two directions (front-back or sun-moon directions for both travel direction and head direction). After including the strategy as an additional factor or excluding subjects based on implausible parameter estimates from the Weibull function fitting procedure, we observed much stronger motion aftereffects, indicating that we were able to successfully replicate the results from Experiment 1 in this more challenging scenario.

### Motion aftereffects scale with duration of adaptation

In Experiment 3 (*n* = 28 participants; 16 females), we attempted to address remaining questions from Experiment 1 and 2 about the adaptation duration. An alternative explanation for the “opposite” adaptation effect is that the adaptation time used in the previous two experiments might be too long or too short to induce a sufficient adaptation effect for travel direction. We initially set the adaptation time (i.e., 60s initial adaptation, 10s top-ups) based on previous visual adaptation studies, but since adaptation effects of travel direction have never been studied before, we had few *a priori* expectations regarding whether adaptation duration would produce larger or smaller effects. Based on previous visual adaptation literature, the magnitude of a classic motion after-effect should scale depending on the amount of adaptation, as increased adaptation time yields a greater decrease in the neural responsiveness to the same stimuli (***Fang et al., 2005; Fang and He, 2005; Leopold et al., 2005; Vautin and Berkley, 1977***). To take a closer look at whether and how adaptation time affects motion aftereffects of travel direction, we conducted an experiment where we tested four adaptation time periods: 18s, 36s, 54s, and 72s. Motion was in the direction of the hallway like in Experiment 1, and for simplicity we only used the sun direction for this study. Because we added more adaptation conditions, we also only tested three levels of percentage of actual movement in the adaptation direction: 30, 50, and 70% (see Methods Experiment 3).

We were again able to successfully replicate the primary results from Experiment 1 and Experiment 2. We observed a tendency for a difference between the experimental and control conditions at the 18s adaptation time period (*F* (1, 27) = 3.88, *p* = 0.059, 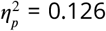, 95%*C.I*. = [0.000, 0.370]). The effect grew to become significant at 36s (*F* (1, 27) = 5.20, *p* = 0.031, 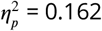, 95%*C.I*. = [0.000, 0.408]), as well as at 54s (*F* (1, 27) = 13.25, *p* = 0.001, 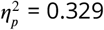, 95%*C.I*. = [0.070, 0.555]), and 72s adaptation time periods (*F* (1, 27) = 7.22, *p* = 0.012, 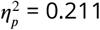, 95%*C.I*. = [0.011, 0.455]). This result suggests that the magnitude of the travel motion aftereffect scales with adaptation time (Fig. 5, Supplemental Information Fig. 18).

**Figure 5.**
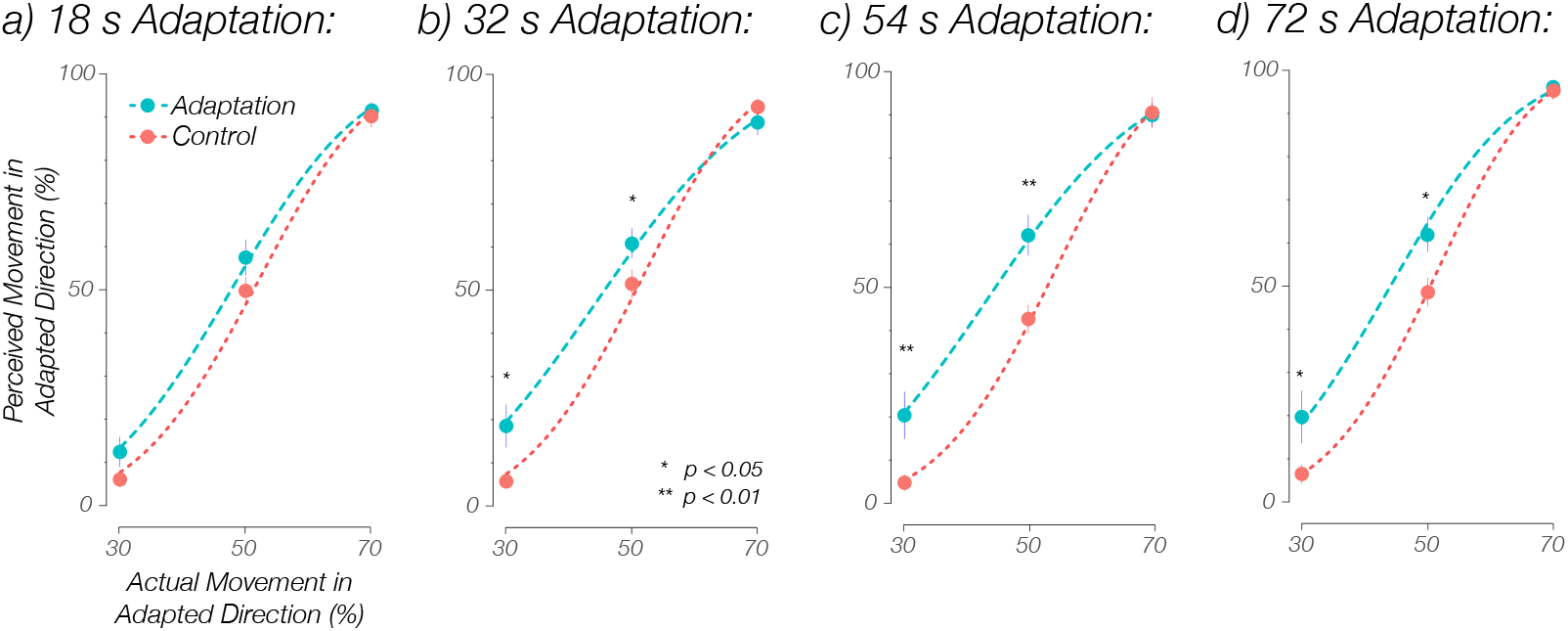
Experiment 3. The perceived percentage of movement in the adaptation direction compared with the actual percentage for all subjects, separated by adaptation time periods (n = 28). Solid dots indicate the grand average value, and error bars indicate standard errors. Dashed lines indicate the average psychometric Weibull functions. a) The reported rate for 18s adaptation trials. The adaptation condition showed a trend for higher reported percentages than the control conditions (*p* = 0.059). b) The reported rate for 36s adaptation trials. The adaptation condition showed significantly higher overall reported percentages than the control conditions (*p* = 0.031), particularly at 30% (*p* = 0.015) and 50% (*p* = 0.032), supporting the aftereffect in the same direction of travel. c) The reported rate for 54s adaptation trials. The adaptation condition showed significantly higher overall reported percentages than the control conditions (*p* = 0.001), particularly at 30% (*p* = 0.007) and 50% (*p* = 0.001), supporting the aftereffect. d) The reported rate for 72s adaptation trials. The adaptation condition showed significantly higher reported overall percentages than the control conditions (*p* = 0.012), particularly at 30% (*p* = 0.047) and 50% (*p* = 0.032), supporting the aftereffect in the same direction of travel. The bias psychometric function (i.e., *α*) marginally shifted toward a lower percentage of reported movement toward the adaptation direction when adapted at 72s adaptation (*p* = 0.056) time period, but the shift was not significant at 18s (*p* = 0.315), 36s (*p* = 0.407), or 54s (*p* = 0.669) adaptation time periods. The uncertainty psychometric function (i.e., *β*) indicates that observers’ detectability of the difference between the two directions was decreased by adaptation but were not significant at 18s (*p* = 0.543) or 36s (*p* = 0.132) adaptation time periods, but was significant at 54s (*p* = 0.017) adaptation time period and marginally significant at 72s (*p* = 0.074) adaptation time period. * p *<* 0.05, ** p *<* 0.01, Tukey correction.

Next, we made pair-wise comparisons between the different adaptation time conditions at each percentage of actual movement in the adaptation direction (Supplemental Information Fig. 16a-f, Supplemental Information Fig. 17, Supplemental Information Fig. 19). This analysis revealed that when the actual percentage was at 70%, the 72s adaptation trials had significantly higher perceived percentage than the 36s adaptation trials (*p* = 0.014) (Fig. 16e) and marginally higher than the 54s adaptation trials (*p* = 0.055) (Fig. 16f).

All patterns of results were maintained after excluding 4 subjects (around 14% of subjects) based on implausible parameter estimates from the Weibull function fitting procedure (Supplemental Information Fig. 20 - Fig. 23). These findings are consistent with motion aftereffects scaling with adaptation time. These findings were replicated in an independent sample of participants (*n* = 31 participants; 16 females), with slightly different instructions and some blocks having adaptation to the moon direction (see Supplemental Information Fig. 24 - 29). In addition, the main pattern of results did not shift after controlling for strategies in the analyses for both Experiment 3 and the replication (see Methods Experiment 3 Data Analysis, Supplemental Information Data Analysis, Experiment 3, Experiment 4). The replication (i.e., Experiment 4, see Supplement) precludes the possible explanation of demand characteristics because the different instructions occasionally indicated that adaptation would be to the opposite direction than what actually occurred, yet the effect remained in those situations.

Further, Weibull analyses showed that when the adaptation time was 54s (*t* (27) = −2.54, *p* = 0.017, *Cohen’ s d* = −0.616, 95%*C.I*. = [−1.146, −0.086]) and 72s (*t* (27) = −1.86, *p* = 0.074, *Cohen’ s d* = −0.146, 95%*C.I*. = [−0.623, 0.331]), people had significantly more uncertainty in making judgments in the adaptation condition compared to the control condition. When adaptation time was 72s, there was a tendency to bias responses (*α*) toward the adaptation direction (*t* (27) = −2.00, *p* = 0.056, *Cohen’ s d* = −0.575, 95%*C.I*. = [−1.199, 0.048]) (Fig. 5).

Overall, the results in Experiment 3 successfully replicated effects from Experiment 1 and Experiment 2. Importantly, we observed the same “opposite” motion aftereffect of travel direction for differing adaptation time lengths, precluding the alternative explanation that the “opposite” motion aftereffect was due to the adaptation time being too long or too short. Furthermore, the travel aftereffect was scaled with adaptation duration, such that longer adaptation duration tended to have larger aftereffects. Together, these findings provide additional support for a motion aftereffect for travel direction that is independent of head direction.

## Discussion

In a series of experiments, we employed a method from visual perception in a novel way to study travel direction during self-motion. We observed systematic travel motion aftereffects across all experiments in both raw data (Fig. 2, Fig. 4, Fig. 5) and filtered data (Supplemental Information Fig. 13, Fig. 15, Fig. 20). The aftereffect was not due to response bias for a particular cardinal direction, approaching effects, or serial position effects. Moreover, the motion aftereffect scaled to a longer adaptation time span.

The aftereffects were observed in the same direction as the adapted travel direction, which fits the characteristics of previous high-level motion aftereffects paradigms (***Culham et al., 1998, 2000***). First, the adaptation of travel direction we observed was likely high-level. It is unclear whether a high-level aftereffect is more likely to be implemented on the level of a single neuron or through neural systems. We designed the study so that the possible influence of low-level optic flow information would be cancelled out due to the constant changes of head direction in the experiment. Second, subjects were instructed to focus on the global net change in position, meaning that they had to integrate their movement over time. Third, the test phase was also dynamic, requiring integration of travel direction over time. The current task differs from previous motion adaptation experiments in that we let participants take a first-person (egocentric) perspective to attentively track self-motion, rather than simply viewing stimuli move across the screen (e.g., the waterfall effect). Taken together, this novel experimental design makes the effects we observed in the current study unique among high-level motion aftereffects.

This “opposite” motion aftereffect (which is actually in the same direction of travel) indicates that a non-opponent process underlies the travel direction system. Several previous studies have also found motion aftereffects in the direction of travel, but via different sensory modalities, including the *podokinetic aftereffect* where spatial orientation is changed via remodeling somatosensory signals between the trunk and feet (***Gordon et al., 1995; Weber et al., 1998; Earhart et al., 2001***), and the *jogging/running-in-place aftereffect* that involves recalibration of visuomotor control systems (***Anstis, 1995; Durgin and Pelah, 1999; Mulavara et al., 2010***).

What could be the possible relationship between the head direction system and the travel direction system in the brain? It is difficult to find direct answers to this question because, as mentioned in the introduction, the two direction systems have typically been conflated. However, we may get some clues from studies in which head direction and travel direction are perfectly aligned. Several studies have investigated heading direction using adaptation paradigms or “repetition suppression” in fMRI to look at the sensitivity to heading direction of various cortical visual motion areas (***Baumann et al., 2010; Cardin et al., 2012; Shine et al., 2016***). Researchers observed clear head direction adaptation in MT, MST (***Cardin et al., 2012***), medial parietal lobe (***Baumann et al., 2010***), as well as bilateral retrosplenial cortex, thalamus, and precuneus (***Shine et al., 2016***). We can take from these results that when head direction and travel direction are aligned, there is still adaptation taking place in the brain. The brain areas that demonstrate adaptation are higher-level motion systems, suggesting that these systems are involved in encoding heading direction in the human brain.

Possible travel direction pathways are more speculative. They could involve independent sensory inputs (e.g., vision, somatosensation) and feed-forward high-level motion processing pathways (e.g., MST, parietal lobe, hippocampus, etc.) (e.g., (***Chrastil et al., 2016; Sherrill et al., 2015***)). Recent findings of bi-directional cells in rodent dysgranular retrosplenial cortex (***Jacob et al., 2017***) may also be a good candidate for cells that are sensitive to travel direction. Further research using fMRI and computational modeling is needed to shed light on the relationship between the travel direction and head direction systems in the human brain, and the degree to which they have independent circuitry.

## Conclusion

In conclusion, we found high-level motion aftereffects of travel direction using a novel motion adaptation paradigm, which suggests that travel direction is a fundamental component of human navigation and indicates how it might be represented in the brain. Interestingly, the aftereffect is in the opposite direction to traditional motion aftereffects, suggesting that adapting to a travel direction will result in greater perception of moving towards the adapting direction. Critically, we dissociated head direction from travel direction across all experiments, indicating that travel direction has separate neural mechanisms from head direction in the human brain. Considering travel direction as a basic navigation component provides a new path to understanding the question of how people form their sense of direction. Furthermore, the results will encourage scientists who study navigation behavior of other species (rodents, birds, insects, etc.) to look for more direct neurological evidence of travel direction, rather than only test for head direction.

## Materials and Methods

### Experiment 1

#### Participants

Participants consisted of 77 University of California, Santa Barbara (UCSB) undergraduates who participated in return for course credit or monetary incentive ($12/hour). Our task is novel, thus there is limited previous data to use for a power analysis. Therefore, we based our sample size on previous studies of movement adaptation that used within-subjects designs (***Culham et al., 2000; Earhart et al., 2001***) and desktop navigation tasks (***Weisberg and Newcombe, 2016; Weisberg et al., 2014***). These studies yielded a target of 30 participants per condition. We used the outcomes of Experiment 1 for power analysis of subsequent experiments. The four criteria for prescreening participants were 1) no history or a current condition of psychiatric problems, 2) no learning disability or attention deficit disorder, 3) not currently taking psychoactive drugs, and 4) no history of head trauma.

Participants were discarded for either not completing both control and adaptation sessions (*n* = 15), responding with the same key all the time (*n* = 1), or responding too slowly (*RT >* 10 s; *n* = 1). The final pool consisted of 60 participants (29 males, 31 females; 38 not Hispanic or Latino, 15 Hispanic or Latino, 7 not reported; 20 Asian, 18 White, 2 African American, 3 American Indian/Alaskan Native, 1 Native Hawaiian/Pacific Islander, 3 more than one race, 13 not reported), with 30 participants randomly assigned in the moon condition virtual environment (16 males, 14 females) and 30 participants randomly assigned in the sun condition virtual environment (13 males, 17 females). Ages of the remaining 60 participants ranged from 18 to 30 (mean 20.32). All participants signed an informed consent form in agreement with the UCSB Institutional Review Board requirements in accordance with the Declaration of Helsinki.

#### Stimuli

The virtual environment was generated on a PC (Origin, NVIDIA GeForce GTX 980 graphics card, 15-inch display, 1920 × 1080 pixels display resolution) using Vizard software (WorldViz) to render the images. Participants experienced visual self-motion in a long (8566 virtual units) landmark-free virtual hallway (see Fig. 1). The hallway was long enough such that the visual angle to the end of the hallway did not change during movement and therefore could not act as a cue to distance traveled. The translational speed of self-motion was randomly sampled from 10 - 15 virtual units/second in each trial. The rotational speed of self-motion was randomly sampled from 130° – 150° /second. Speeds for adaptation and test phases were sampled separately in each trial. The hallway consisted of two walls and a ground surface with coarse grained texture; the textures were designed to prevent participants using them for location cues during movement. The walls were always colored dark brown, while the ground was colored light brown during the adaptation phase and was colored green during the test phase (see Fig. 1a - b). At one end of the hallway, in the sky, there was a sun and at the other end there was a moon. The sun and the moon were designed to be cardinal frames of reference and were rendered to move with the viewer at a constant distance. Therefore, the size of the moon and the sun did not change with self-motion, and participants could not evaluate distance change based on perceptual changes in either the sun or moon. In the moon condition, participants would initially move toward the moon frame of reference, and vice versa for the sun condition.

#### Task

The task consisted of an adaptation period followed by test trials (see Fig. 1c-e). In the adaptation phase for both the sun and the moon adaptation groups, each block started with 60 seconds of virtual travel toward the sun or the moon as the global travel direction, depending on their group (see Fig. 1c). During the adaptation phase, participants were instructed to pay attention to the movement in the hallway on the computer screen. To dissociate travel direction from heading direction, we included occasional 180° turns to change the local facing direction while maintaining the constant global travel direction. This heading change occurred through a rotation, such that the view turned around, rather than a sharp flip. The number of turns in all experiments were randomly sampled from a range varied by the time length of each trial (0-2 turns for 10s trials, 5-7 turns for 60s initial adaptation phase).

The control was the same for both groups, where the “adaptation” phase consisted of a still screen without movement. This phase only lasted 10 seconds, but with occasional 180° turns to change the facing direction (see Fig. 1d).

Immediately after the adaptation phase was a 10-second test phase. The test phase was the same for all conditions. The ground in the hallway would turn green, signifying the 10-second test period. In the test phase, the travel direction would change, such that participants experienced back-and-forth movement toward both the sun and the moon. The facing direction changed during the test phase, just like in the adaptation phase, with between 0 and 2 turns. The amount of back-and-forth movement on each trial was expressed in terms of a percentage of movement toward one of the two cardinal frames of reference (see Fig. 1e). The percentage of movement in one direction is complementary to the other direction such that they add up to 100%. For example, 20% of virtual movement toward the sun is equivalent to 80% of virtual movement toward the moon. In order to compare the sun and the moon conditions, all analyses are described as oriented toward the sun direction so that we could easily see the effect of adaptation in each condition.

In addition to the experimental condition (adaptation vs. control), this percentage of virtual movement was the primary independent variable in the study, ranging from 20% to 80% in 10% steps. Participants were instructed to pay attention to the overall direction they traveled in during the test period. When the movement in the test period stopped, on-screen text asked participants to judge whether their movement during the test period was overall more toward the sun or more toward the moon direction. They used their left hand to press the “D” key to indicate that they moved closer to the sun and used their right hand to press the “K” key to indicate that they moved closer to the moon. Although the task was untimed, participants were instructed to respond as quickly and accurately as possible.

As soon as participants pressed a response key, the hallway would turn brown again for 10 seconds as a “top-up” adaptation phase. The top-up adaptation of the next trial started with the same facing direction as the last screen of the previous testing trial so that participants could have a more coherent experience in the virtual environment. During the 10-second top-up phase, participants would experience the same movement as the initial 60-second adaptation (i.e., the same initial travel direction with occasional changes of facing directions) but for a shorter time length. Then, participants would be given another 10-second test of travel direction. In the control condition, the top-up was 10 seconds in the hallway without movement, but with occasional facing changes. The task continued alternating between the original travel direction top-up adaptations (brown) and test phase (green) until the block ended, and then participants could take a break (up to 5 minutes). Reported direction and reaction time for each trial were recorded.

#### Design

A 2 (experimental condition: adaptation, control; within subjects) × 7 (actual percentage of movement toward the sun: 20%, 30%, 40%, 50%, 60%, 70%, 80% rate; within subjects) within-subjects design was used. Each test condition was repeated for 12 trials, for a total of 84 trials (7 test conditions × 12 trials/test condition). These 84 trials were randomly separated into 3 blocks with a short break between blocks. A new 60-second adaptation occurred after each break to reinstate the adaptation. All stimuli were presented in random order for all participants.

In order to control for the influence of a particular adapting direction and of response bias, half of subjects were randomly assigned to the sun adaptation group, and the other half were assigned to the moon adaptation group. We combined the sun and moon adaptation groups for analysis, although we also analyzed them separately (see Supplemental Information). The experiment was conducted over two sessions for each participant, with one session the experimental task and the other session as the control. The order of completing these two sessions were counterbalanced among participants within each group.

#### Procedure

Participants first were greeted in the lab, given information about the study, and given consent forms to sign. They then completed a participant screening form and were given an instruction sheet to learn about the task.

Next, they performed the motion adaptation task. Participants sat approximately 50 cm in front of the computer screen. Before beginning the formal experiment, they were given additional instructions and the experimenter answered any questions. They completed 5 practice trials (the adaptation time and test conditions were different from the experimental trials), and then any additional questions were answered.

Each session lasted approximately 1 hour. Participants completed the two sessions on two separate days to prevent fatigue, with no more than one week between sessions. Finally, after each session, participants were asked to rate the difficulty of the task based on a 1 – 7 Likert scale (1 meant very easy and 7 meant very difficult) and to respond to an open-ended question about their strategy.

#### Data Analysis

R-studio and MATLAB were used for data analysis. We first removed outlier trials that were 3 standard deviations above or below of the mean of each subject’s reaction time. Approximately 0.33% of trials were removed: no trials were removed in the sun group, and in the moon group 0.04% trials were removed from the experimental session and 1.27% trials were removed from the control session. From the remaining trials for each participant, we calculated the reported percentage of movement toward the adaptation direction (i.e., sun or moon direction) as well as mean reaction time for each percentage level.

First, we conducted a 2 (experimental condition: adaptation, control; within subjects) × 7 (actual percentage toward the adaptation direction: 20% - 80% rate of actual movement toward the adaptation direction; within subjects) repeated-measures analysis of variance (ANOVA). The primary comparison was the difference between the adaptation and the control conditions within each actual percentage of movement since a motion aftereffect would lead to a shift in the curve for the adaptation conditions. Because this difference between adaptation and control was the primary outcome measure, we also conducted Tukey HSD paired t-tests between adaptation trials and control trials within each actual percentage of movement.

We then analyzed the data by fitting results with a Weibull function. In the current study, the Weibull function assumes that the perceived movement contrast between the moon and the sun scales proportionally to the signal-to-noise ratio of the actual movement contrast that supports the perception. Separate Weibull functions were fit to individual participants’s data for each experimental condition (adaptation and control) and each percentage of movement in the adaptation direction (20% - 80%) using Palamedes toolbox in MATLAB (***Prins and Kingdom, 2018***). Two parameters were derived for each fitted function: the *α* value (i.e., the point of subjective equality) measures a bias to respond the “sun” or the “moon” direction in the task, and the *β* value reflects the detectability of the difference in the task. We then filtered 9 subjects’ data (3 from the sun adaptation group, 6 from the moon adaptation group) whose *α* or *β* value were beyond 2 standard deviations above or below the mean, and conducted paired t-tests of the *α* and *β* values between the adaptation and control condition. In addition, we re-ran the same ANOVAs of reported rate and reaction time on the filtered data. Throughout the paper, “all subjects” refer to all the subjects whose data were used for the initial analyses, including subjects who were later excluded based on implausible parameter estimates from the Weibull function fitting procedure.

Based on post-study questionnaires, participants generally reported the same strategies for both adaptation and control sessions. More specifically, more than half of the subjects (*n* = 36) reported using counting strategies (e.g., mentally counting time, counting steps, physically counting by tapping fingers, etc.). The next set of subjects (*n* = 18) reported keeping focus on a certain part of the environment (e.g., wall, hallway, ground, sky, etc.) for distance estimation. Each of the remaining people (*n* = 6) used a unique strategy. For the filtered data, there were still subjects using counting (*n* = 31), focusing on a part of the environment (*n* = 14), and a unique strategy (*n* = 6). We controlled for the influence of strategies by adding strategy as a factor in the above ANOVA analyses for reported rate and reaction time.

### Experiment 2

#### Participants

The sample size in Experiment 2 was determined based on power analysis using the results of Experiment 1. We used G*Power software (http://www.gpower.hhu.de/) (***Erdfelder et al., 1996***) with an *α* = 0.05, *power* = 0.8, and *Cohen’ s f* measurement of effect size = 0.176 which is based on the weakest effect size (i.e., the interaction effect of the moon condition, see Extended Data Fig. 7c) from Experiment 1. The resulting sample size for within-group comparison was 24. Using this power analysis, we recruited 33 participants for Experiment 2, which is more than adequate for the main objectives of this study and which matched closely with the participant numbers from the Experiment 1 sun condition. Participants consisted of 33 University of California, Irvine (UCI) students who participated in return for monetary incentive ($12/hour). Three participants were discarded for misunderstanding the task instruction (*n* = 1), wrongly pressing the reverse response buttons (*n* = 1), or not completing both control and adaptation sessions (*n* = 1). The final pool consisted of 30 participants (14 males, 16 females; 20 not Hispanic or Latino, 9 Hispanic or Latino, 1 not reported; 14 Asian, 11 White, 2 African American, 1 American Indian, 1 other, 1 not reported). Ages of the remaining 30 participants ranged from 18 to 34 (mean 22.93). All participants signed an informed consent form in agreement with the UCI Institutional Review Board requirements in accordance with the Declaration of Helsinki.

The Stimuli, Task, Design, and Procedure were the same as Experiment 1, except that tasks were modified such that the head direction was always orthogonal to the travel direction (see Fig. 3). For simplicity, we only conducted adaptation to the sun direction. The initial control adaptation period lasted 60s in Experiment 2.

#### Data Analysis

The analysis was largely the same as Experiment 1. We first removed outlier trials that were 3 standard deviations above or below of the mean of each subject’s reaction time. Approximately 2.12% of trials were removed: 2.18% trials were removed from the experimental session and 2.06% trials were removed from the control session. From the remaining trials for each participant, we calculated the reported percentage of movement toward the sun as well as mean reaction time for each percentage level.

First, we conducted a 2 (experimental condition: adaptation, control; within subjects) × 7 (actual sun percentage: 20% - 80% rate of actual movement toward the sun; within subjects) repeated-measures analysis of variance (ANOVA) analysis. Because the primary comparison was the difference between the adaptation and the control conditions within each actual percentage of movement, we also conducted Tukey HSD paired t-tests between adaptation trials and control trials within each actual percentage of movement. We then filtered 7 subjects’ data and analyzed the data using parameters derived from Weibull fits, similar to Experiment 1.

Based on post-study questionnaires, participants generally reported the same strategies for both adaptation and control sessions. Same as we had observed in Experiment 1, people reported three main types of strategies in Experiment 2: counting strategies (*n* = 13), keeping focus on a certain part of the environment for distance estimation (*n* = 7), and a unique strategy (*n* = 10). For the filtered data, there were still subjects using counting (*n* = 12), focusing on a part of the environment (*n* = 4), and a unique strategy (*n* = 7). Again, we controlled for the influence of strategies by adding strategy as a factor in the above ANOVA analyses for reported rate and reaction time.

### Experiment 3

#### Participants

Similar to Experiment 2, we calculated a sample size of 24 for within-group comparison in Experiment 3 determined based on power analysis using G*Power software (http://www.gpower.hhu.de/) (***Erdfelder et al., 1996***) based on the weakest effect size from Experiment 1 (Extended Data Fig. 7c). We recruited 28 participants for Experiment 3 (12 males, 16 females; 26 not Hispanic or Latino, 1 Hispanic or Latino, 1 not reported; 19 Asian, 9 White), which is more than adequate for the main objectives of this study. All participants were UCI students who participated in return for monetary incentive ($12/hour). Ages of the participants ranged from 18 to 30 (mean 24.71). All participants signed an informed consent form in agreement with the UCI Institutional Review Board requirements in accordance with the Declaration of Helsinki.

The Stimuli and Procedure were the same as Experiment 1.

#### Task

The Task was the same as Experiment 1, except for the following:

1. Each block started with different initial adaptation time periods (18s, 36, 54s, or 72s), correspondingly followed by different top-up time periods (3s, 6s, 9s, or 12s). The control adaptation phases and top-up periods had the same corresponding time lengths. For convenience, we refer to all adaptation trials in terms of their initial adaptation time.
2. Corresponding to the change in the time length for the adaptation phases, we also changed the range of the number of turns of the facing direction that each top-up and adaptation period was sampled from: 0-1 turn for 3s top-ups, 0-2 turns for 6s top-ups, 0-2 turns for 9s top-ups, 0-2 turns for 12s top-ups, 1-3 turns for 18s adaptation periods, 3-5 turns for 36s adaptation periods, 4-6 turns for 54s adaptation periods, 6-8 turns for 72s adaptation periods. Ranges were calculated based on the following equations:

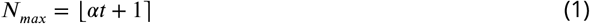

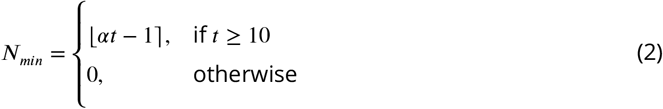

*N* denotes number of turns. *α* is a coefficient which was set to 0.1.*t* denotes the number of time step in the trial (either the adaptation or the test phase). └ ┐ denotes round to the nearest integer.

3. To keep the experiment within two sessions, we used three percentages of virtual movement (30%, 50%, and 70%) and reduced the number of trials in each percentage to 9.

#### Design

A 2 (experimental condition: adaptation, control; within subjects) × 4 (adaptation time blocks: 3s top-up with 18s initial adaptation, 6s top-up with 36s initial adaptation, 9s top-up with 54s initial adaptation, 12s top-up with 72s initial adaptation; within subjects) × 3 (actual percentage of movement toward the sun: 30%, 50%, 70% rate; within subjects) within-subject design was used. Each test condition was repeated for 9 trials, for a total of 108 trials (4 blocks x 3 percentages/block × 9 trials/percentage, not including the initial trial of each block). Each of the 4 blocks corresponded to one adaptation time condition. The order of the 4 blocks was counterbalanced across subjects. The trials with the 3 different percentages of actual movement toward the sun were presented in random order within each block. There was a short break between blocks. A new initial adaptation period occurred after each break to initiate adaptation of a different magnitude.

The experiment was conducted over two sessions for each participant, with one session the experimental task and the other session as the control. The order of completing these two sessions was counterbalanced.

#### Data Analysis

The analysis was largely the same as Experiment 1. We first removed outlier trials that were 3 standard deviations above or below of the mean of each subject’s reaction time. Approximately 1.93% of trials were removed: 1.79% trials were removed from the experimental session and 2.08% trials were removed from the control session. From the remaining trials for each participant, we calculated the reported percentage of movement toward the sun as well as mean reaction time for each percentage level at each adaptation condition (the initial adaptation trial in each block were not included).

First, we conducted a 2 (experimental condition: adaptation, control; within subjects) × 3 (actual sun percentage: 30%, 50%, 70% rate of actual movement toward the sun; within subjects) repeatedmeasures ANOVA for each adaptation time periods (18s, 36s, 54s, or 72s) separately. Because the primary comparison was the difference between the adaptation and the control conditions within each actual percentage of movement, we also conducted Tukey HSD paired t-tests between adaptation trials and control trials within each actual percentage of movement.

Next, we conducted a 4 (adaptation time periods: 18s, 36s, 54s, or 72s; within subjects) × 3 (actual sun percentage: 30%, 50%, 70% rate of actual movement toward the sun; within subjects) repeated-measures ANOVA for each experimental condition (control, adaptation) separately. Because the primary comparison was the difference between different adaptation time periods within each actual percentage of movement, we also conducted Tukey HSD paired t-tests between different adaptation time periods within each actual percentage of movement. We then filtered and analyzed the data using the Weibull function, similar to Experiment 1. We filtered 4 subjects’ data from all conditions based on subjects whose results were excluded by more than one adaptation time period condition.

Same as we have observed in Experiment 1 and 2, people reported three main types of strategies in Experiment 3: counting strategies (*n* = 13), keeping focus on a certain part of the environment for distance estimation (*n* = 13), and a unique strategy (*n* = 2). For the filtered data, there were still subjects using counting (*n* = 10), focusing on a part of the environment (*n* = 12), and a unique strategy (*n* = 2). Again, we controlled for the influence of strategies by adding strategy as a factor in the above ANOVA analyses for reported rate and reaction time.

## Data Availability

All data generated or analysed during this study, as well as codes, will be available on Open Science Framework.

## Acknowledgments

We thank R. Davis, B. Tranquada-Torres, N. Hatamian, S. Kannan, A. Barel, N. Lavoy, O. Cooper, N. Krohn for helping with data collection. The research was supported by National Science Foundation grant (no. BCS-1829398). The funder had no role in study design, program, data collection and analysis, decision to publish or preparation of the manuscript.

## Supplemental Information

### Experimental Videos

**Video 1**. Experiment 1 Videos

**Video 1—source data 1**. adaptation phase

**Video 1—source data 2**. control phase

**Video 1—source data 3**. test phase 20% condition

**Video 1—source data 4**. test phase 50% condition

**Video 2**. Experiment 2 Video

**Video 2—source data 1**. adaptation and test phases

### Experiment 1: Reaction Time

### Experiment 1: Separate Results for Sun and Moon Groups

**Appendix 0—figure6.**
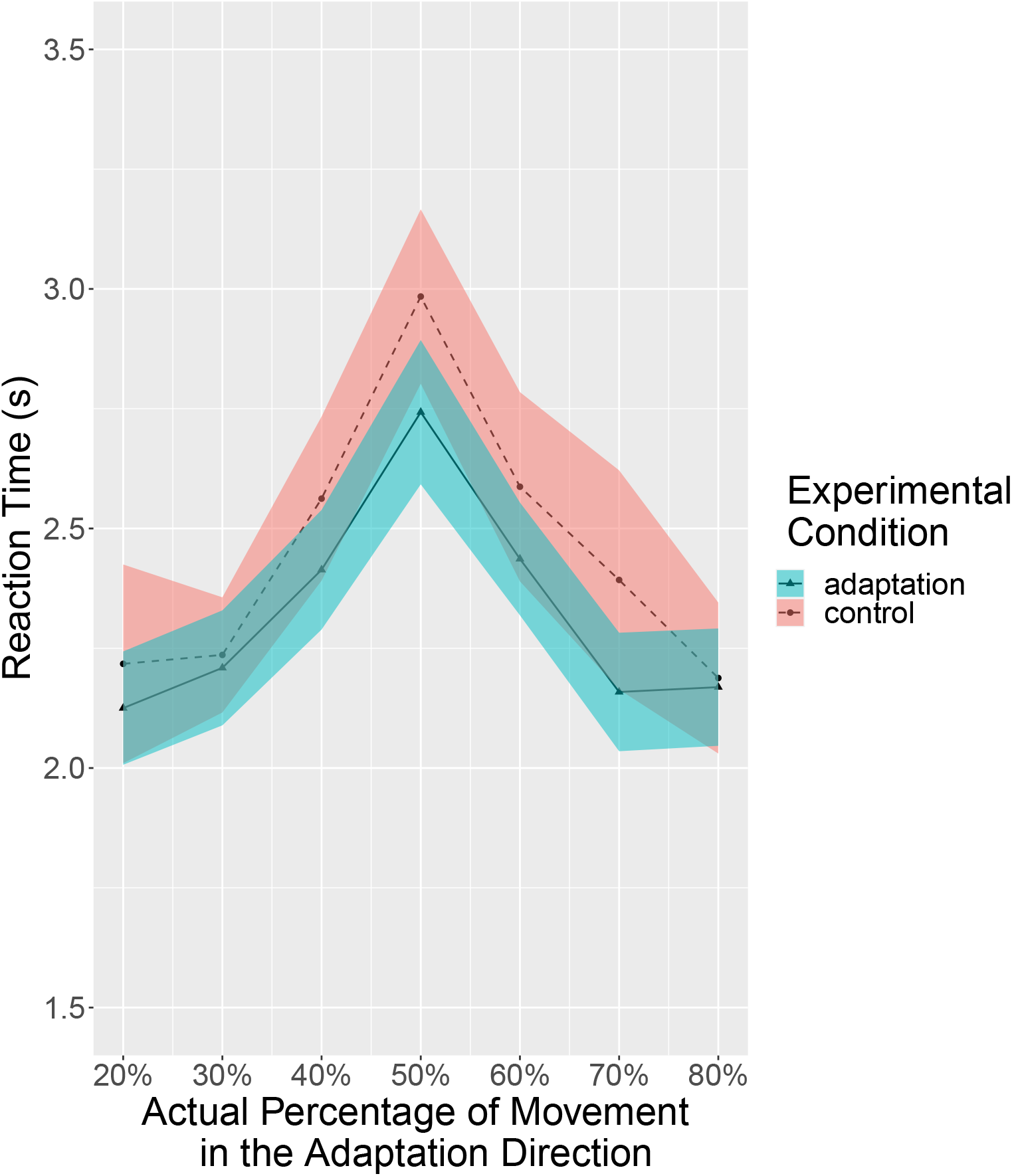
Experiment 1: Reaction times of all subjects(n=60). The reaction time increased as the actual percentage approached 50%. Solid lines indicate the grand average value, and the shaded area indicate 1 standard error of means.

**Appendix 0—figure7.**
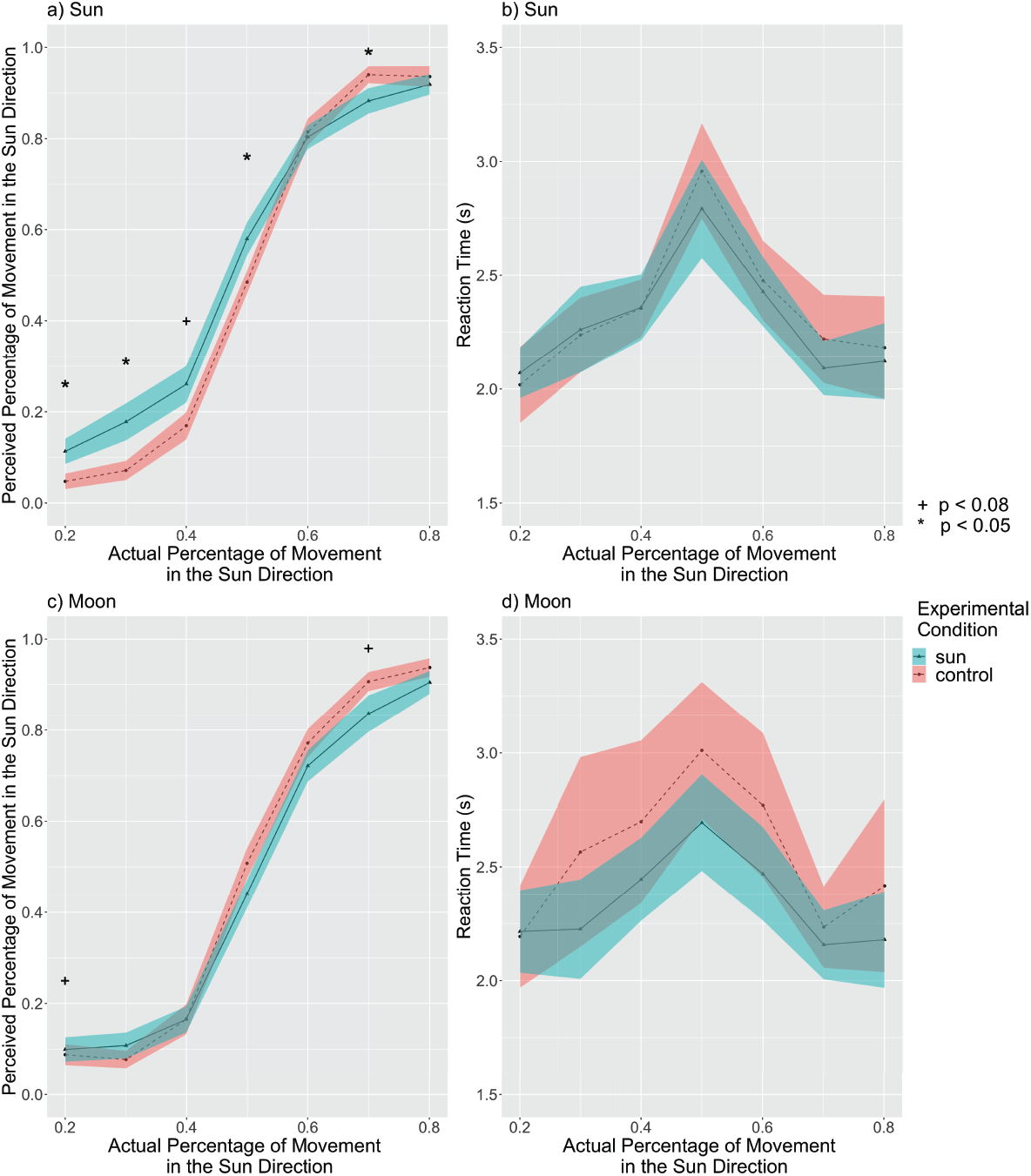
The perceived percentage of movement in the sun direction compared with the actual percentage for all subjects. a) The reported rate of the sun group. The sun adaptation condition had an overall higher reported percentage than the control condition. This result suggests an aftereffect in the same direction of travel (*p* = 0.012). The adaptation condition showed higher reported percentages than corresponding control conditions at 20%, 30%, 40%, and 50%, supporting this aftereffect. This result also suggests an aftereffect in the same direction of travel. There was also a significant interaction between condition and the actual percentage. b) Reaction times of the sun group. The reaction time increased as the actual percentage approached 50%. c) The reported rate of the moon group. The moon adaptation had a marginally lower reported percentage overall than the control condition (*p* = 0.051). This result suggests an aftereffect in the same direction of travel. The adaptation condition showed marginally lower reported percentages than corresponding control conditions at the 50% and 70%, supporting this aftereffect. d) The reaction time of the moon group. The reaction time increased as the actual percentage got closer to 50%. Solid lines indicate the grand average value, and the shaded area indicate 1 standard error of means. ^+^ p *<* 0.08; * p *<* 0.05, Tukey correction.

### Experiment 1: Serial Position Effects for Sun and Moon Adaptation Groups

In order to determine if any portion of the 10-second test phase contributed to the reported rate, we tested for a serial position effect. A serial position effect occurs when people tend to remember the first items (the *primacy effect*) or the last items (the *recency effect*) best in a list. In this case, if there is a serial position effect for movements within the 10-second test trial, people’s performance could be influenced by the movement towards the sun in the first few seconds (primacy effect) or the last few seconds (recency effect) of movement. For example, if by chance the last 3 seconds of movement in the 10-second test phase was in the sun direction, perhaps people would be more inclined to select “sun”, even if the overall movement was more toward the moon. If people’s judgements were based on only a portion of the test trial, then the actual percentage of the 10-second trial was not fully used for judgements. Therefore, we first recalculated the “actual percentage” assuming people only made judgements based on the first 1 - 9 seconds or the last 1-9 seconds separately and then conducted the same analysis protocol used in the full 10-second test trials to see if the adaptation effect would be found in certain partial movement steps.

For convenience, in both groups, “percentage” refers to the percentage of movement in the sun direction.

#### Primacy Effects

We first calculated the actual movement percentage of each of the 9 possible first time steps (i.e., 1 s, 1 s – 2 s, 1 s – 3 s, 1 s – 4 s, 1 s – 5 s, 1 s – 6 s, 1 s – 7 s, 1 s – 8 s, and 1 s – 9 s). Next, we conducted the 2-way experimental condition × actual percentage within-subjects repeated measures ANOVA and then Tukey HSD paired t-tests for adaptation trials and control trials within each actual percentage in each of the 9 first time steps. Note that the possible actual percentages vary in different time steps. For perceived percentage, results in both the sun group (see Figure 8a) and the moon group (see Figure 8b) revealed “opposite aftereffects” that adaptation increased the reported rate in the same direction, which is similar to the previous full 10-second results. Also similar to previous results, reaction time in both the sun group (see Figure 9a) and the moon group (see Figure 9b) did not have any differences between the adaptation condition and the control condition. Therefore, no primacy effect was found to affect the results in either the sun group or the moon group.

**Appendix 0—figure8.**
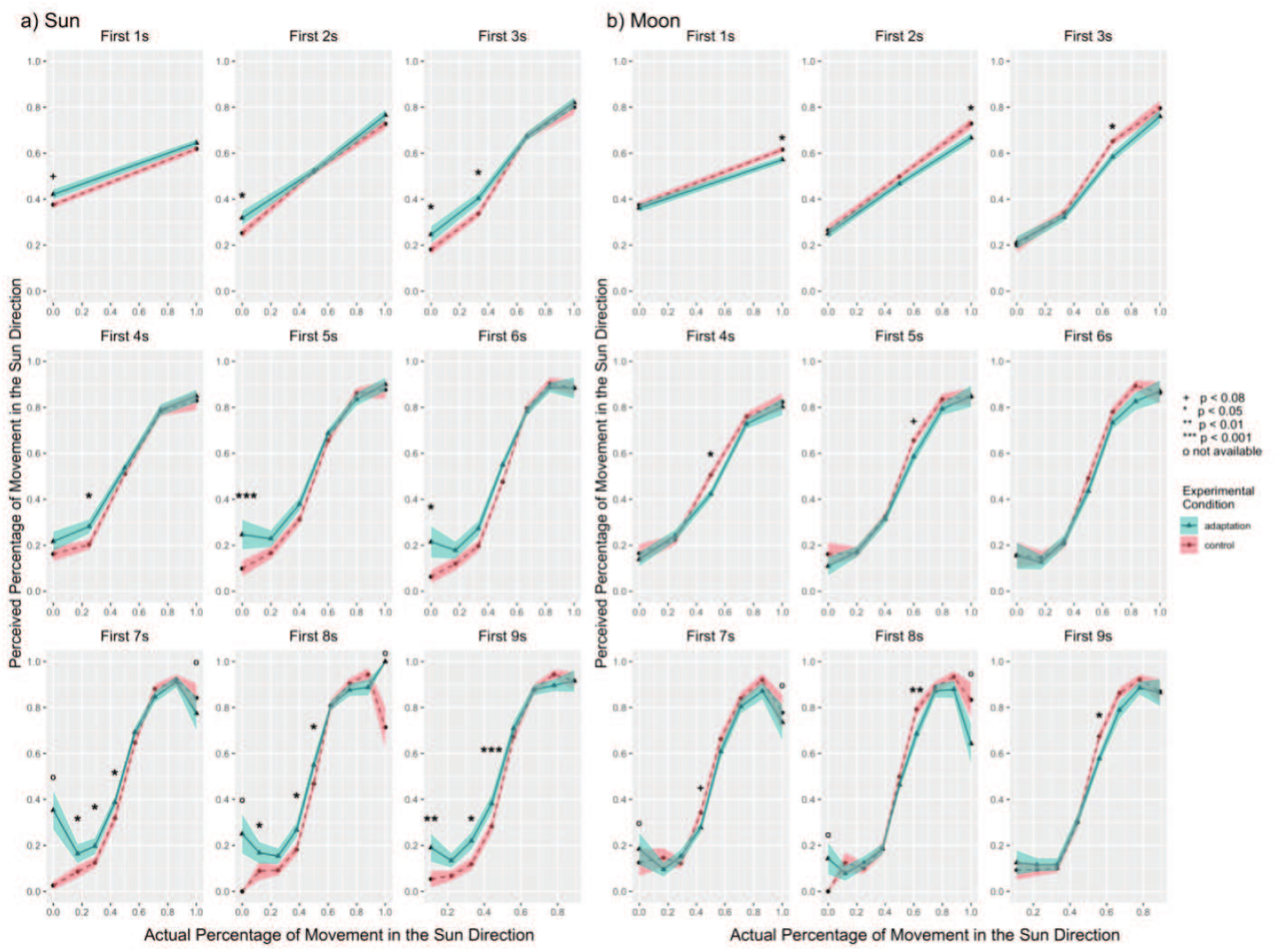
Experiment 1: We looked for primacy effects by examining the reported percentage of the first 1 – 9 seconds of travel. a) The sun group. Adaptation toward the sun direction increased the perceived percentage of movement in the sun direction, similar to the full 10-second trial. b) The moon group. Adaptation toward the moon direction decreased the perceived percentage of movement in the sun direction, i.e., increased perceived percentage of movement in the moon direction, similar to the full 10-second trial. Solid lines indicate the grand average value, and the shaded area indicate 1 standard error of means. ^+^ p *<* 0.08; * p *<* 0.05; ** p *<* 0.01; *** p *<* 0.001; o not enough data for analysis under the corresponding condition, Tukey correction.

**Appendix 0—figure9.**
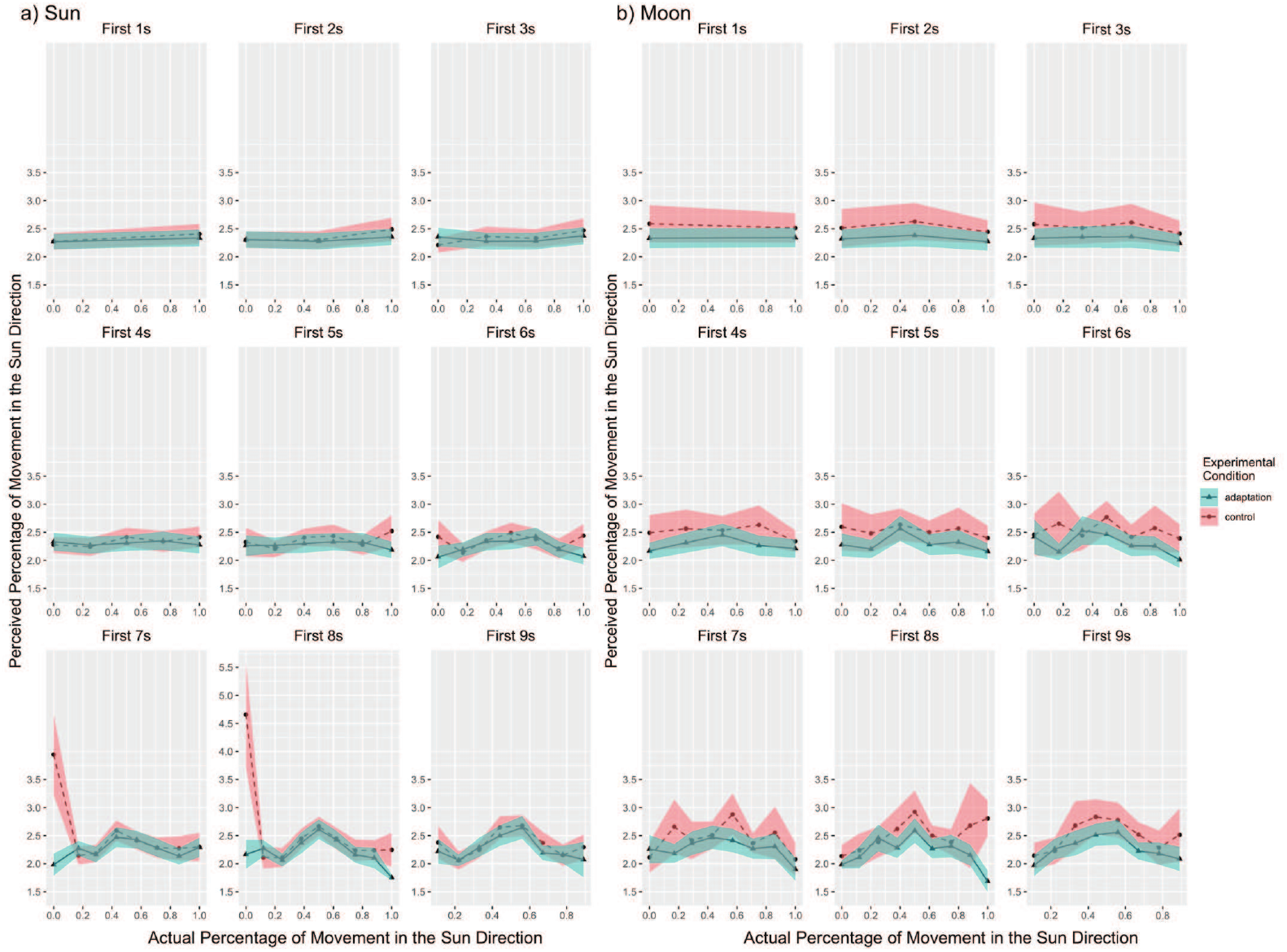
Experiment 1: We looked for primacy effects by examining the reaction time of the first 1 – 9 seconds of travel. a) The sun group. We did not observe any differences between the adaptation condition and the control condition. b) The moon group. We did not observe any differences between the adaptation condition and the control condition. Solid lines indicate the grand average value, and the shaded area indicate 1 standard error of means.

#### Recency Effects

We first calculated the actual movement percentage of each of the 9 possible last time steps (i.e., 10 s, 9 s – 10 s, 8 s – 10 s, 7 s – 10 s, 6 s – 10s, 5 s – 10 s, 4 s – 10 s, 3 s – 10 s, and 2 s – 10 s). Next, we conducted the 2-way experimental condition × actual percentage within-subjects repeated measures ANOVA and then Tukey HSD paired t-tests for adaptation trials and control trials within each actual percentage in each of the 9 last time steps. Note that the possible actual percentages vary in different time steps. The results of the perceived percentage in both the sun group (see Figure 10a) and the moon group (see Figure 10b) showed similar “opposite aftereffects” as found in the full 10-second trial analyses. Also similar to previous analyses, the results of reaction time in both the sun group and the moon group did not show any differences between the adaptation condition and the control condition. Therefore, no recency effect was found to affect the results in either the sun group (see Figure 11a) or the moon group (see Figure 11b). The lack of primacy or recency effects suggests that people used the entire test phase to make their judgments.

**Appendix 0—figure10.**
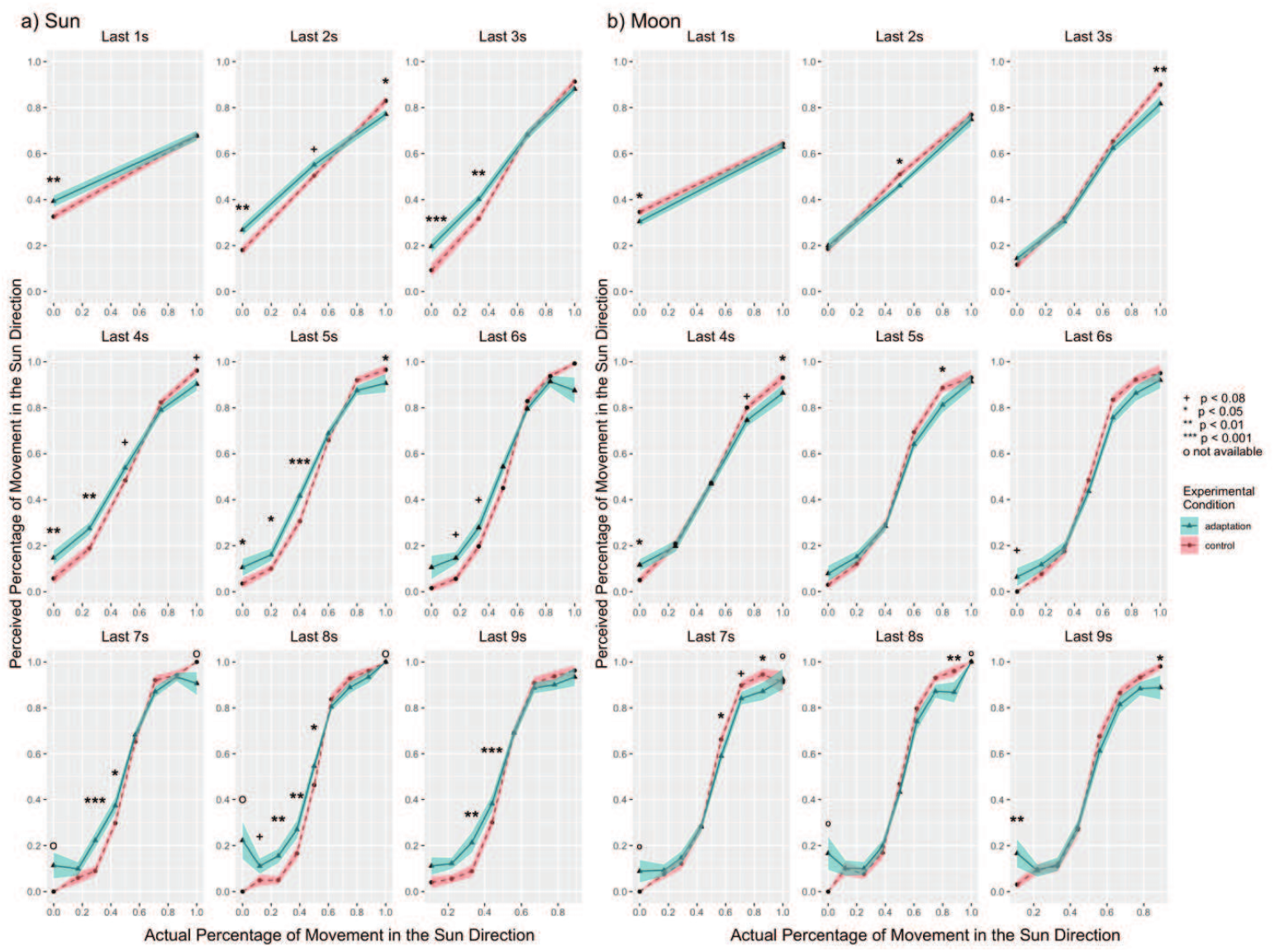
Experiment 1: We looked for recency effects by examining the reported percentage of the last 1 – 9 seconds of travel. a) The sun group. Adaptation toward the sun direction increased the perceived percentage of movement in the sun direction, similar to the full 10-second trial. b) The moon group. Adaptation toward the moon direction decreased the perceived percentage of movement in the sun direction, i.e., increased perceived percentage of movement in the moon direction, similar to the full 10-second trial. Solid lines indicate the grand average value, and the shaded area indicate 1 standard error of means. ^+^ p *<* 0.08; * p *<* 0.05; ** p *<* 0.01; *** p *<* 0.001; o not enough data for analysis under the corresponding condition, Tukey correction.

**Appendix 0—figure11.**
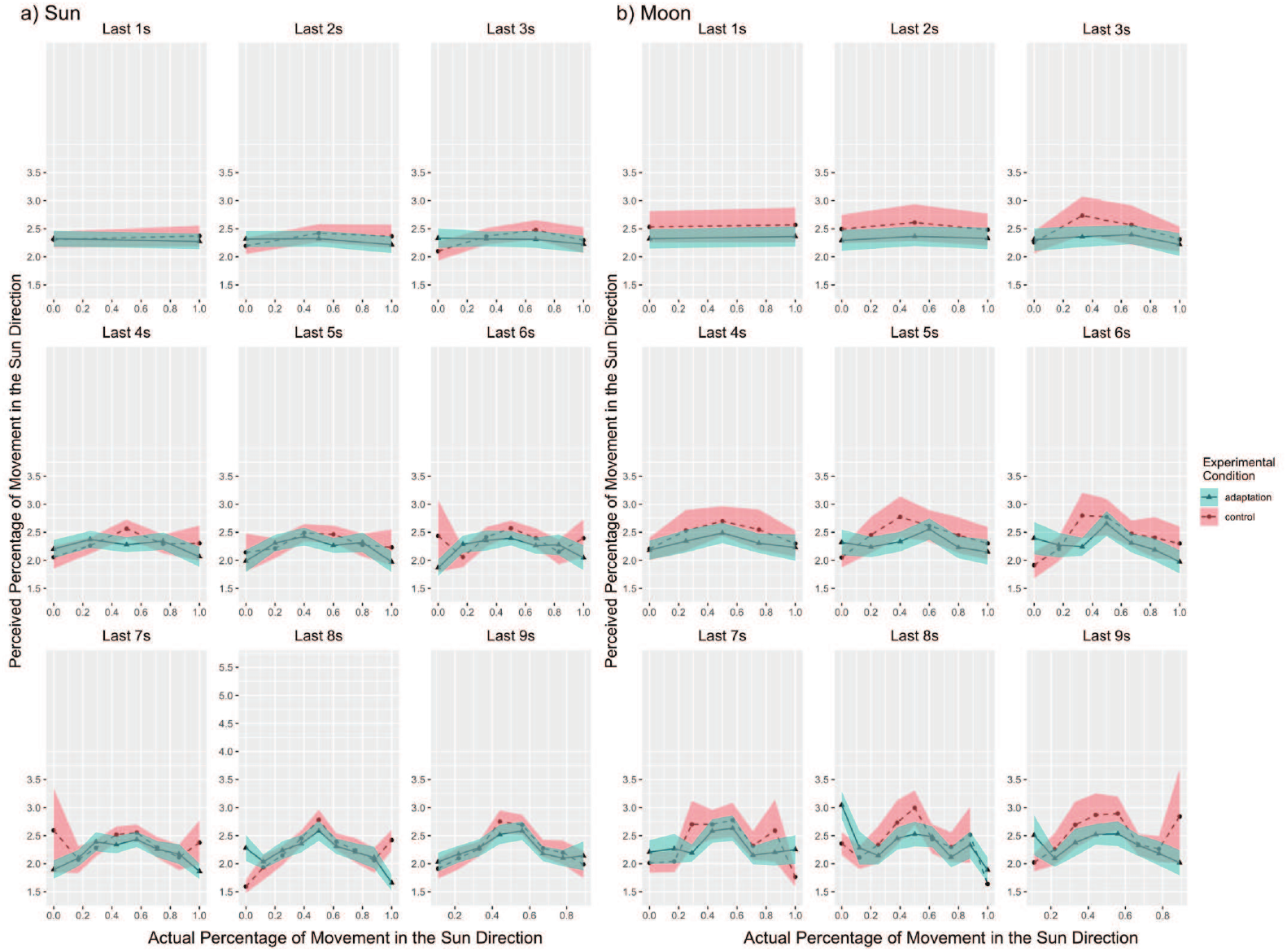
Experiment 1: We looked for recency effects by examining the reaction time of the first 1 – 9 seconds of travel. a) The sun group. We did not observe any differences between the adaptation condition and the control condition. b) The moon group. We did not observe any differences between the adaptation condition and the control condition. Solid lines indicate the grand average value, and the shaded area indicate 1 standard error of means.

### Experiment 1: Initial Trial Results for Sun and Moon Adaptation Groups

**Appendix 0—figure12.**
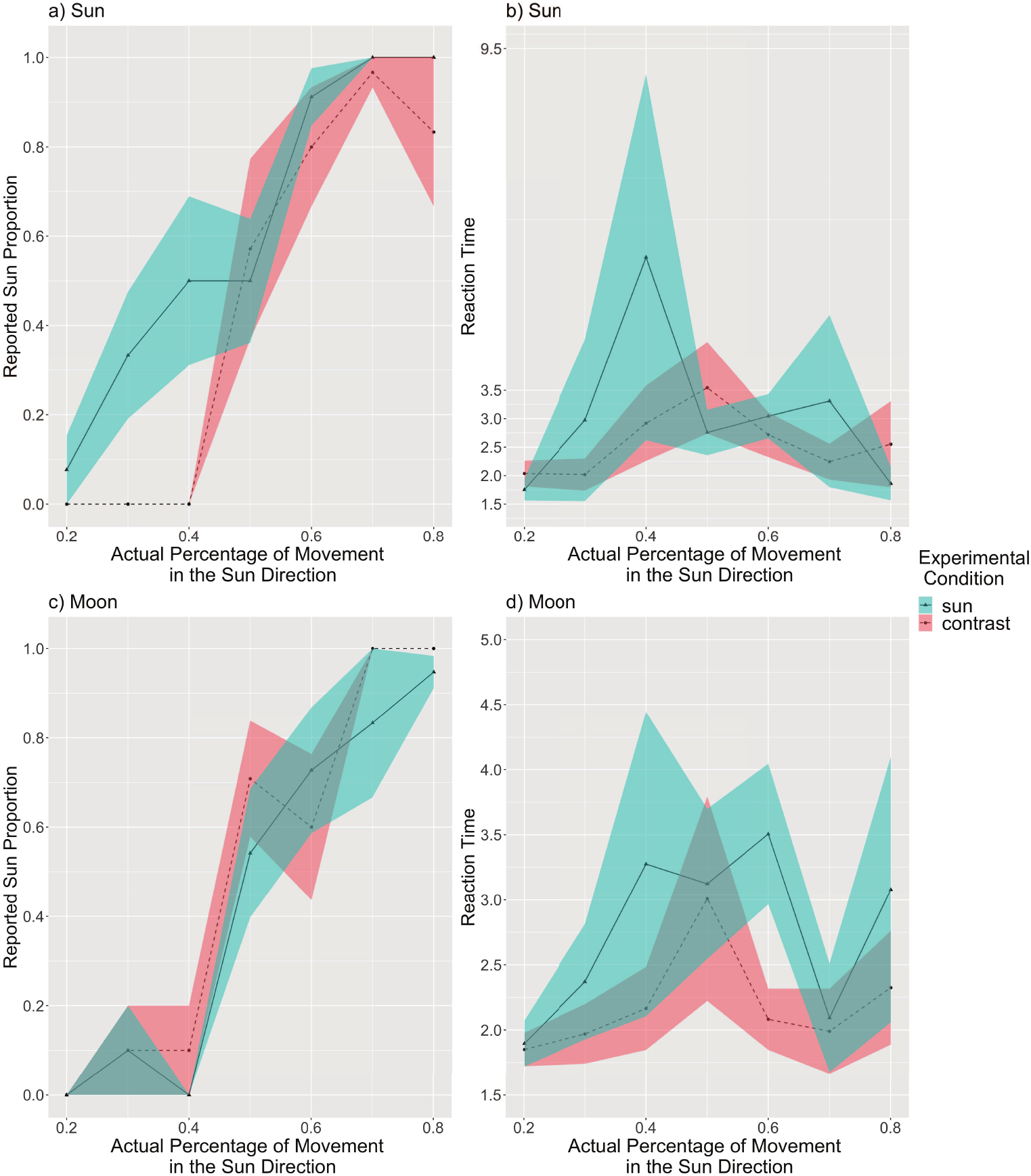
The perceived percentage of movement in the sun direction compared with the actual percentage for all initial trials. a) The reported rate of the sun group. The sun adaptation condition showed a tendency of overall higher reported percentage than the control condition. b) Reaction times of the sun group. No difference were observed in reaction time between the two conditions. c) The reported rate of the moon group. The moon adaptation showed a tendency of lower reported percentage overall than the control condition. d) The reaction time of the moon group. Reaction times of the sun group. No difference were observed in reaction time between the two conditions. Solid lines indicate the grand average value, and the shaded area indicate 1 standard error of means. No inferential statistical analyses was conducted due to not enough initial trials.

### Experiment 1: Filtered Data

**Appendix 0—figure13.**
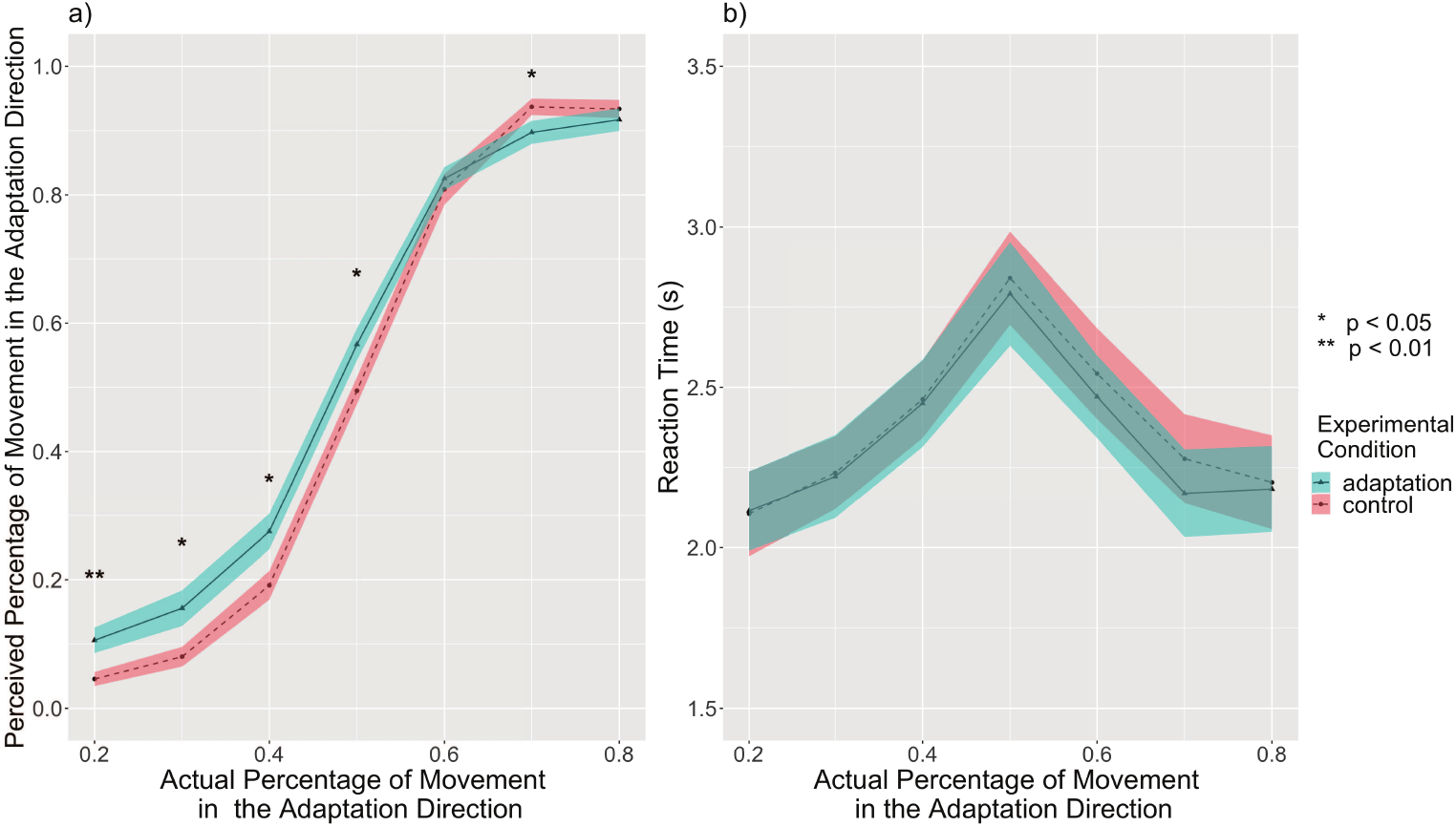
Experiment 1: The perceived percentage of movement in the adaptation direction compared with the actual percentage for filtered subjects (n = 51). a) The reported rate of filtered subjects. The adaptation condition had an overall higher reported percentage than the control condition (*p* = 0.002). The adaptation condition showed higher reported percentages than corresponding control conditions at 20%, 30%, 40%, and 50%, supporting the aftereffect. This result suggests an aftereffect in the same direction of travel. There was also a significant interaction between condition and the actual percentage. b) Reaction times of filtered subjects. The reaction time increased as the actual percentage approached 50%. Solid lines indicate the grand average value, and the shaded area indicate 1 standard error of means. * p *<* 0.05; ** p *<* 0.01, Tukey correction.

### Experiment 2: Reaction Time

#### Raw Data

**Appendix 0—figure14.**
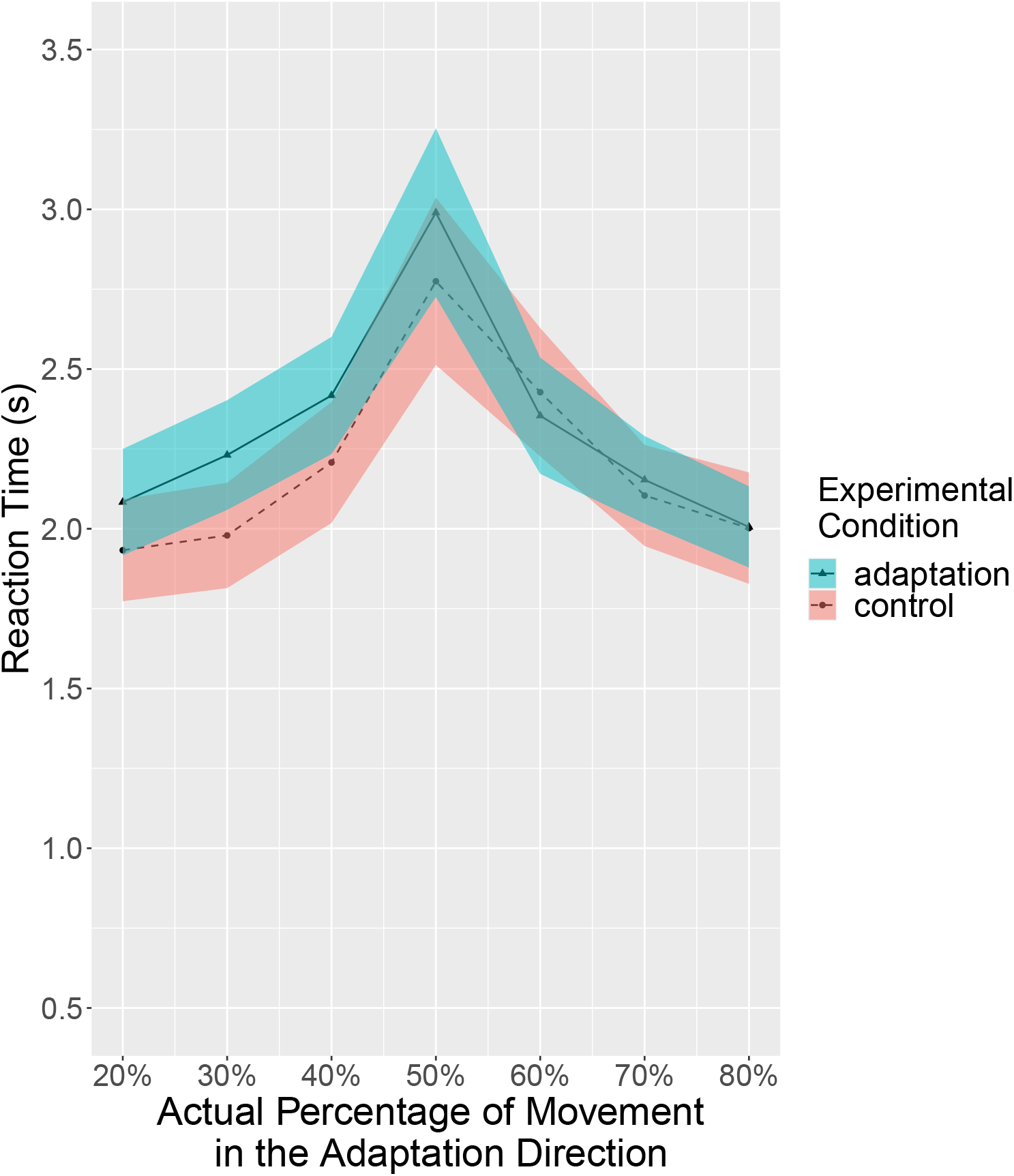
Experiment 1: Reaction times of all subjects (n=30). The reaction time increased as the actual percentage approached 50%. Solid lines indicate the grand average value, and the shaded area indicate 1 standard error of means.

### Experiment 2: Filtered Data

**Appendix 0—figure15.**
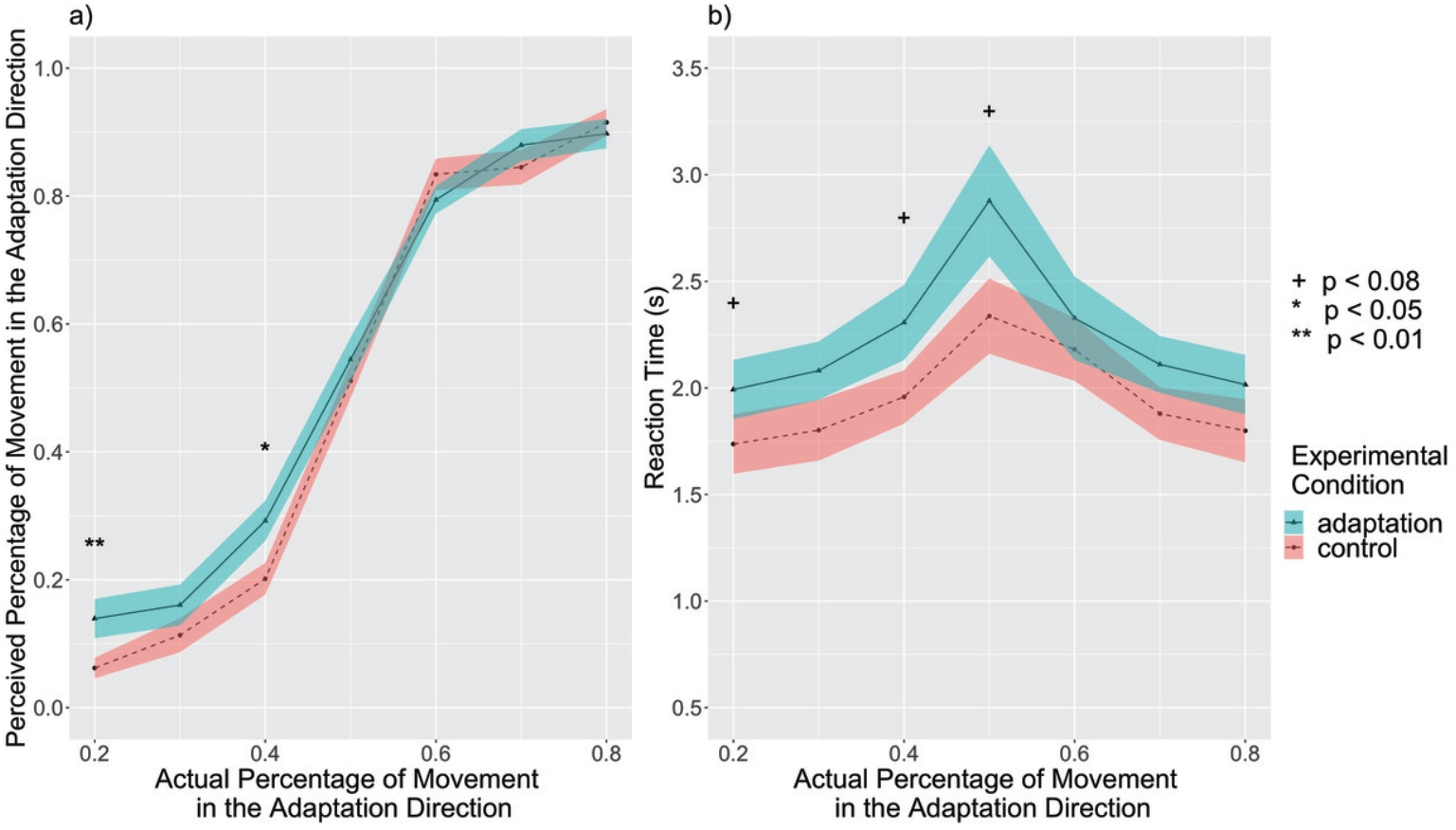
Experiment 2: The perceived percentage of movement in the adaptation direction compared with the actual percentage for filtered subjects (n = 23). a) The reported rate of filtered subjects. The adaptation condition had an overall marginally higher reported percentage than the control condition (*p* = 0.078). The adaptation condition showed higher reported percentages than corresponding control conditions at 20% and 40%, supporting the aftereffect. This result suggests an aftereffect in the same direction of travel. There was also a marginally significant interaction between condition and the actual percentage. b) Reaction times of filtered subjects. The reaction time increased as the actual percentage approached 50%. Solid lines indicate the grand average value, and the shaded area indicate 1 standard error of means. ^+^ p *<* 0.08; * p *<* 0.05; ** p *<* 0.01, Tukey correction.

### Experiment 3: Raw Data

#### Reported Rate by Adaptation Time Periods

**Appendix 0—figure16.**
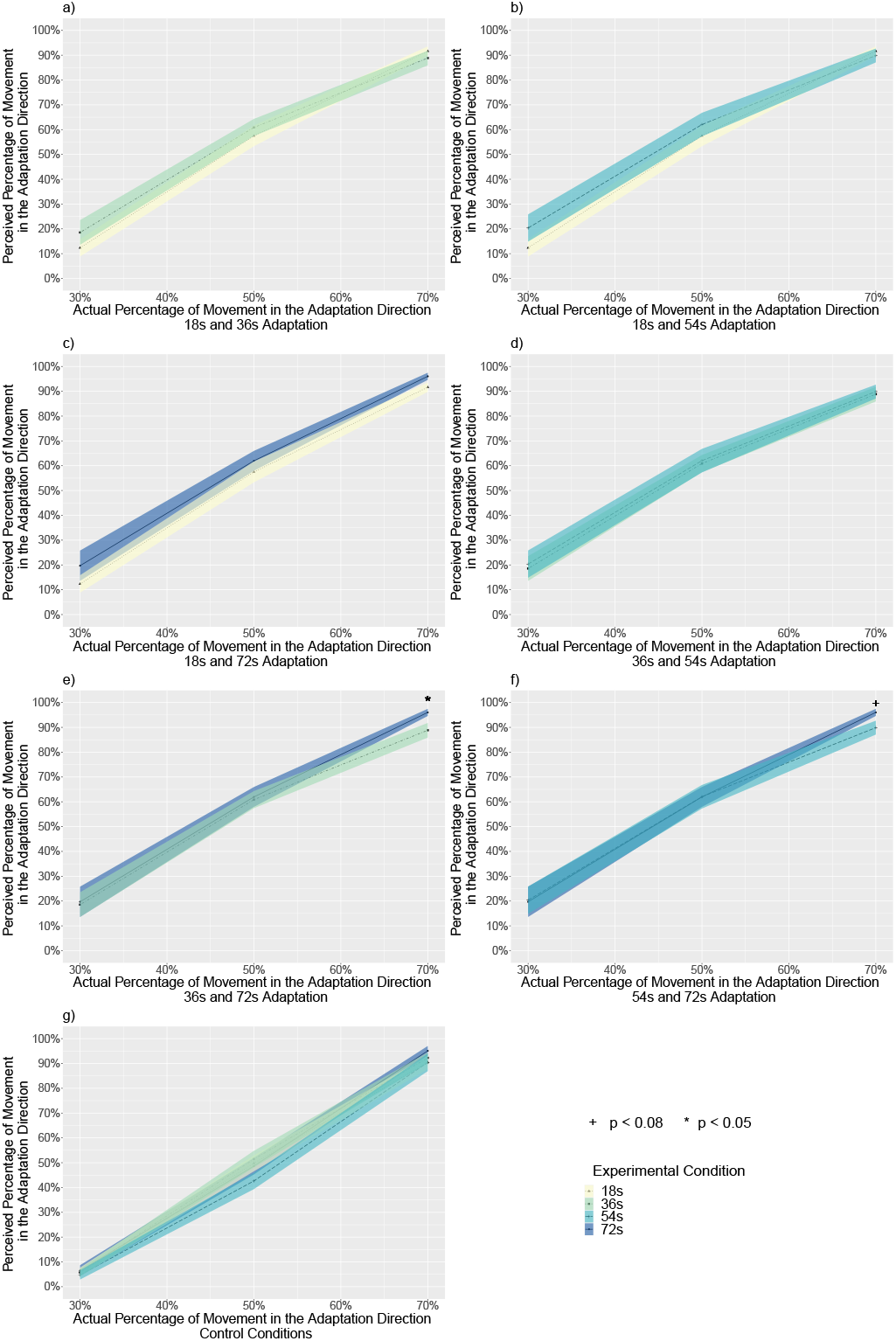
Experiment 3. The perceived percentage of movement in the adaptation direction compared with the actual percentage for all subjects, separated by experimental conditions (n = 28). a) The reported rate for the 18s and 36s adaptation conditions. b) The reported rate for the 18s and 54s adaptation conditions. c) The reported rate for the 18s and 72s adaptation conditions. d) The reported rate for the 36s and 54s adaptation conditions. e) The reported rate for the 36s and 72s adaptation conditions. At 70%, 72s adaptation trials had a significantly higher perceived percentage than 36s adaptation trials (p = 0.014), supporting that the aftereffect increased with adaptation time. The aftereffect is in the same direction of travel. f) The reported rate for the 54s and 72s adaptation conditions. At 70%, 72s adaptation trials had slightly higher perceived percentage than 54s adaptation trials (p = 0.055), supporting that the aftereffect increased with adaptation time. The aftereffect is in the same direction of travel. g) The reported rate for all four control conditions. There were no differences between trials with different adaptation time periods within any actual percentage. Solid lines indicate the grand average value, and the shaded area indicate 1 standard error of means. + p *<* 0.08, * p *<* 0.05. All results are reported with the Tukey correction for multiple comparisons.

#### Reported Rate by Control Conditions

**Appendix 0—figure17.**
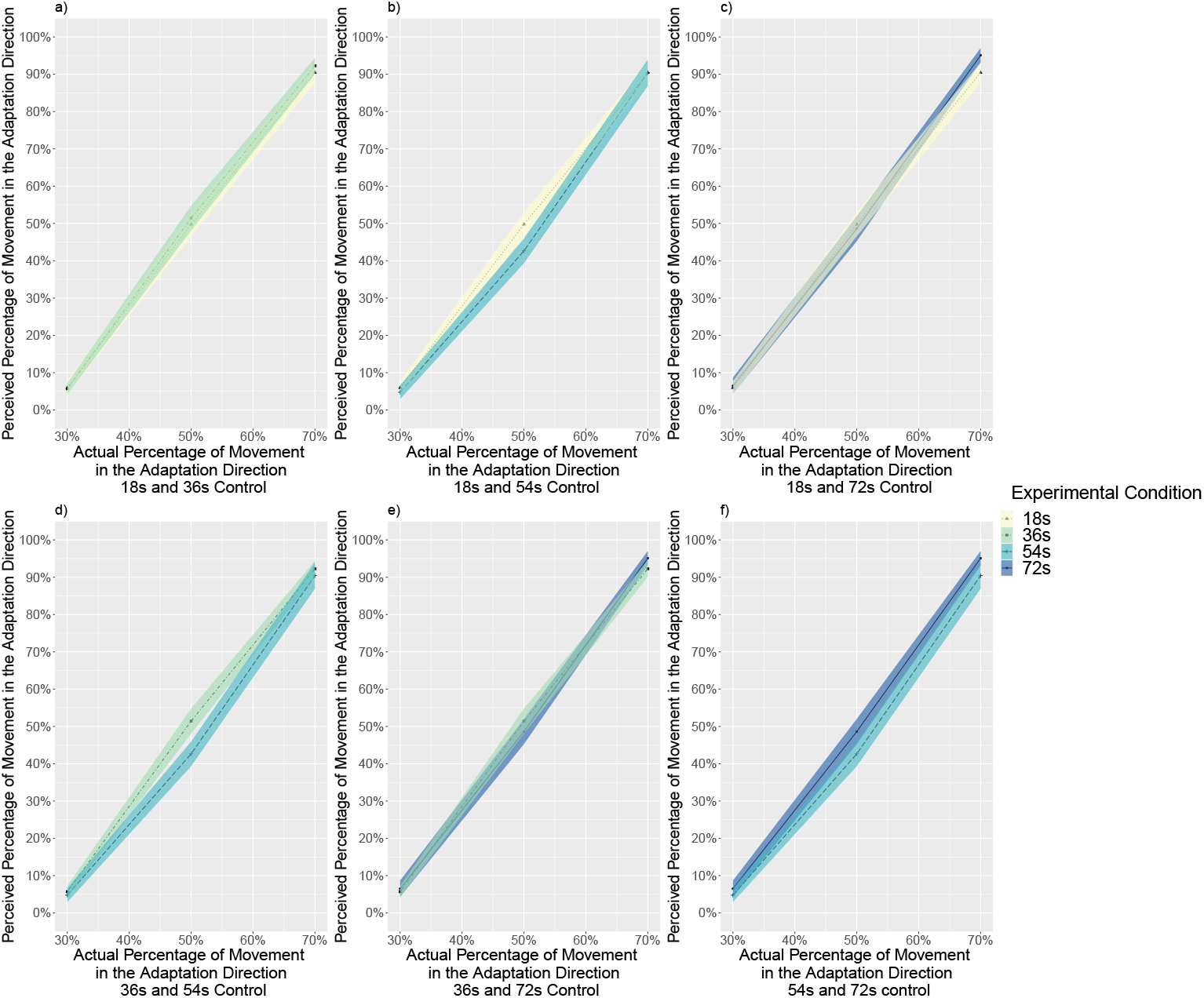
Experiment 3: The perceived percentage of movement in the adaptation direction compared with the actual percentage for all subjects, separated by control conditions (n = 28). a) The reported rate for the 18s and 36s control conditions. b) The reported rate for the 18s and 54s control conditions. c) The reported rate for the 18s and 72s control conditions. d) The reported rate for the 36s and 54s control conditions. e) The reported rate for the 36s and 72s control conditions. f) The reported rate for the 54s and 72s control conditions. Solid lines indicate the grand average value, and the shaded area indicate 1 standard error of means. All results are reported with the Tukey correction for multiple comparisons.

#### Reaction Times by Adaptation Time Periods

**Appendix 0—figure18.**
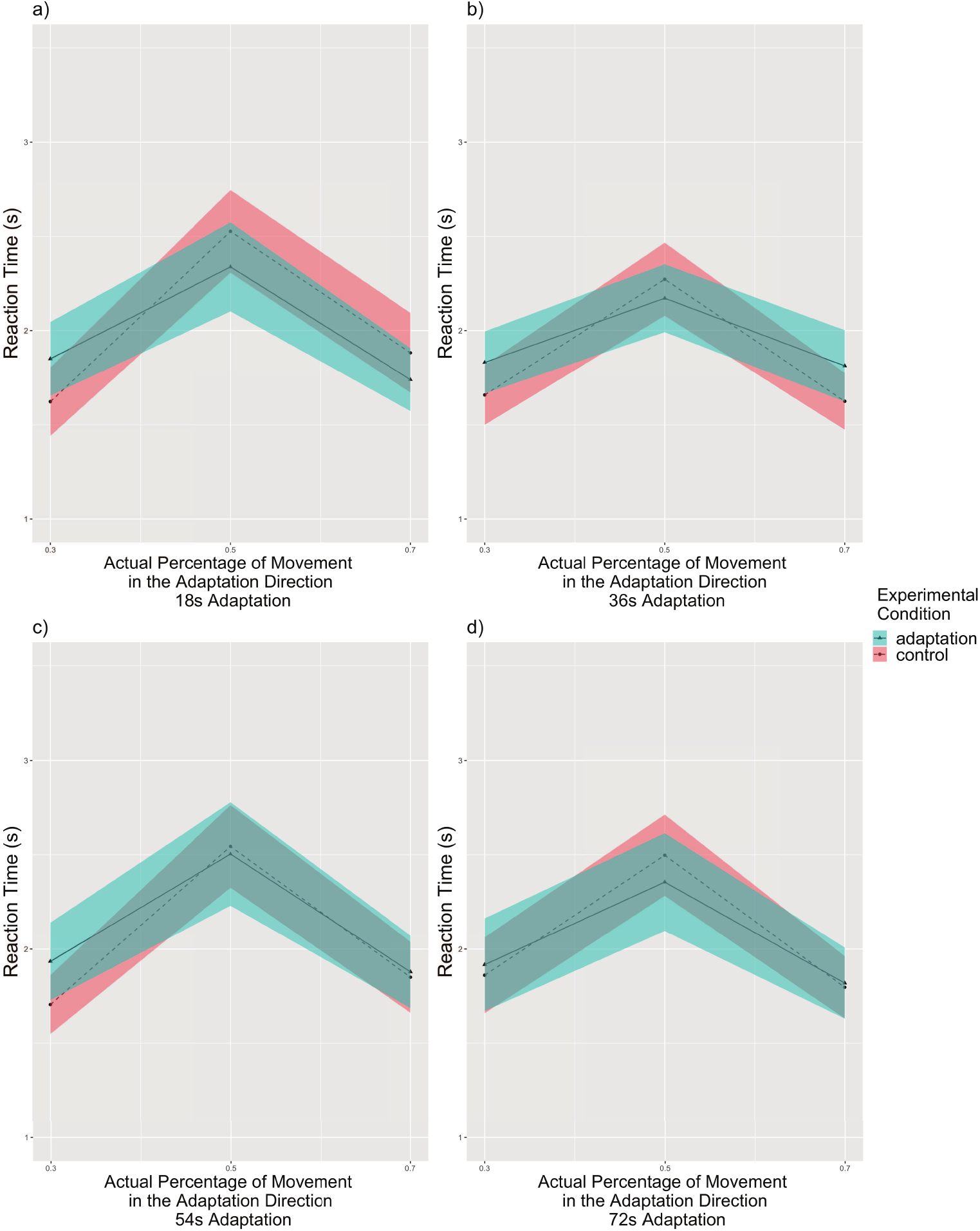
Experiment 3: Reaction times compared with the actual percentage for all subjects, separated by adaptation time periods (n = 28). The reaction time increased as the actual percentage approached 50% in all adaptation time periods. a) Reaction times for 18s adaptation trials. b) Reaction times for 36s adaptation trials. c) Reaction times for 54s adaptation trials. d) Reaction times for 72s adaptation trials. Solid lines indicate the grand average value, and the shaded area indicate 1 standard error of means. Tukey correction.

#### Reaction Times by Experimental Conditions

**Appendix 0—figure19.**
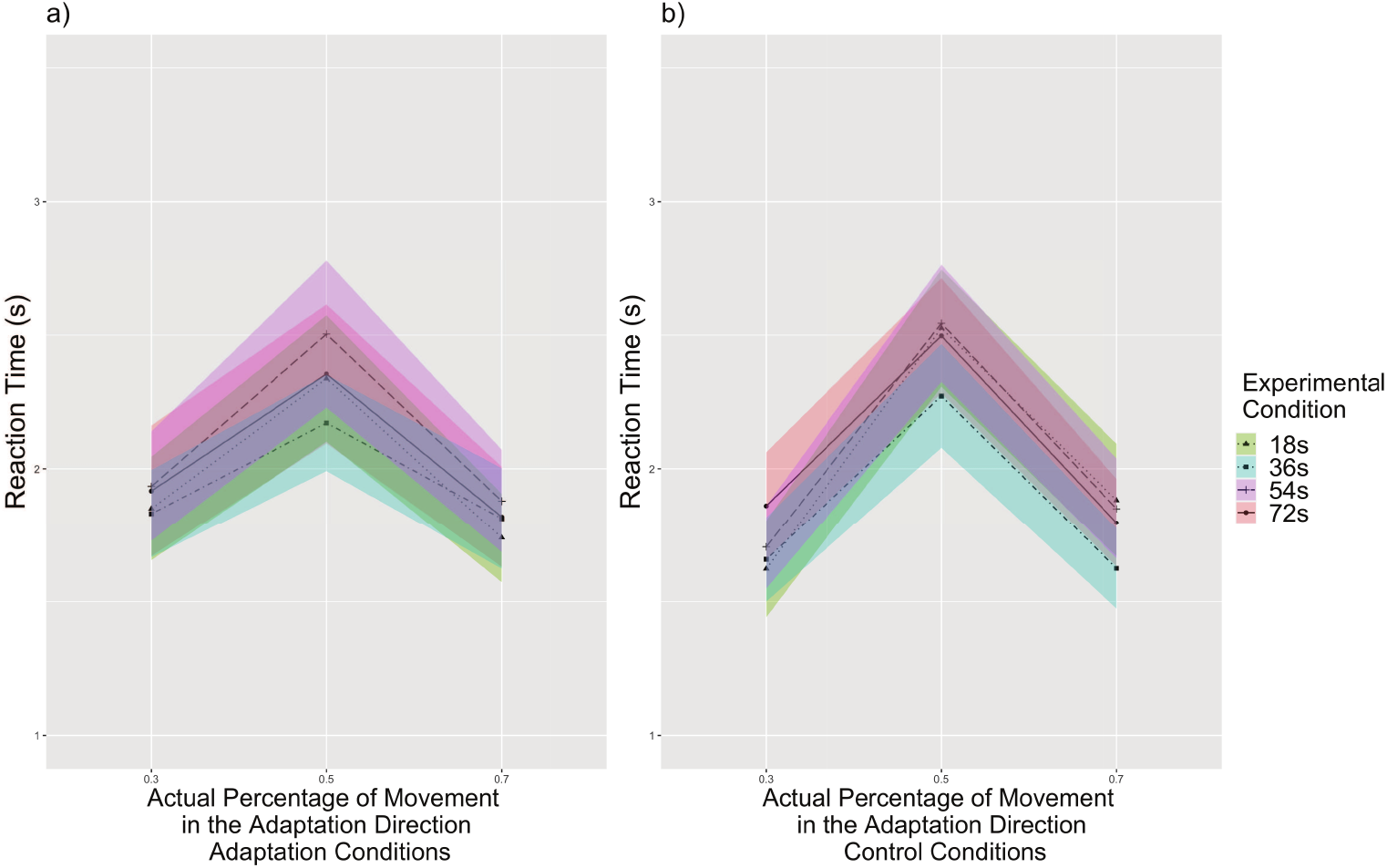
Experiment 3: Reaction times compared with the actual percentage for all subjects, separated by experimental conditions (n = 28). The reaction time increased as the actual percentage approached 50% in both experimental conditions. a) Reaction times for adaptation condition. b) Reaction times for control condition. At 30%, people responded slightly slower to 72s adaptation trials than to 18s adaptation trials (p = 0.054). At 70%, people responded slightly slower to 54s adaptation trials than to 36s adaptation trials (p = 0.074). Solid lines indicate the grand average value, and the shaded area indicate 1 standard error of means. Tukey correction.

### Experiment 3: Filtered Data

#### Reported Rate by Adaptation Time Periods

**Appendix 0—figure20.**
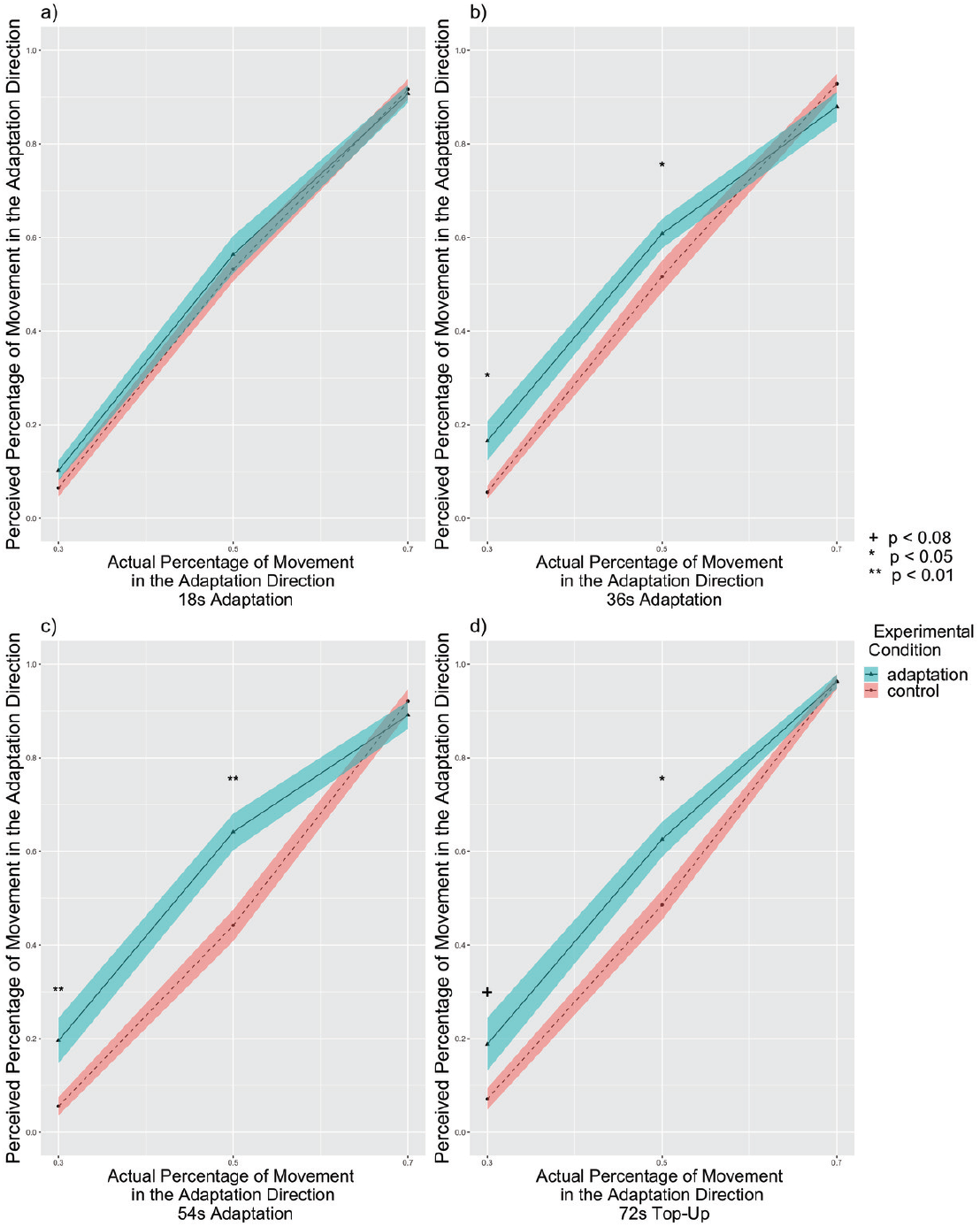
Experiment 3: The perceived percentage of movement in the adaptation direction compared with the actual percentage for filtered subjects, separated by adaptation time periods (n = 24). a) The reported rate for 18s adaptation trials. The adaptation condition showed no difference from the control condition (p = 0.367). b) The reported rate for 36s adaptation trials. The adaptation condition showed slightly higher reported percentages than control conditions (p = 0.061) particularly at 30% (p = 0.013) and 50% (p = 0.048), supporting the aftereffect in the same direction of travel. c) The reported rate for 54s adaptation trials. The adaptation condition showed higher reported percentages than control conditions (p = 0.004) particularly at 30% (p = 0.009) and 50% (p = 0.001), supporting the aftereffect. d) The reported rate for 72s adaptation trials. The adaptation condition showed higher reported percentages than control conditions (p = 0.022) particularly at 30% (p = 0.076) and 50% (p = 0.027), supporting the aftereffect in the same direction of travel. Solid lines indicate the grand average value, and the shaded area indicate 1 standard error of means. + p *<* 0.08, * p *<* 0.05, ** p *<* 0.01, Tukey correction.

#### Reaction Times by Adaptation Time Periods

**Appendix 0—figure21.**
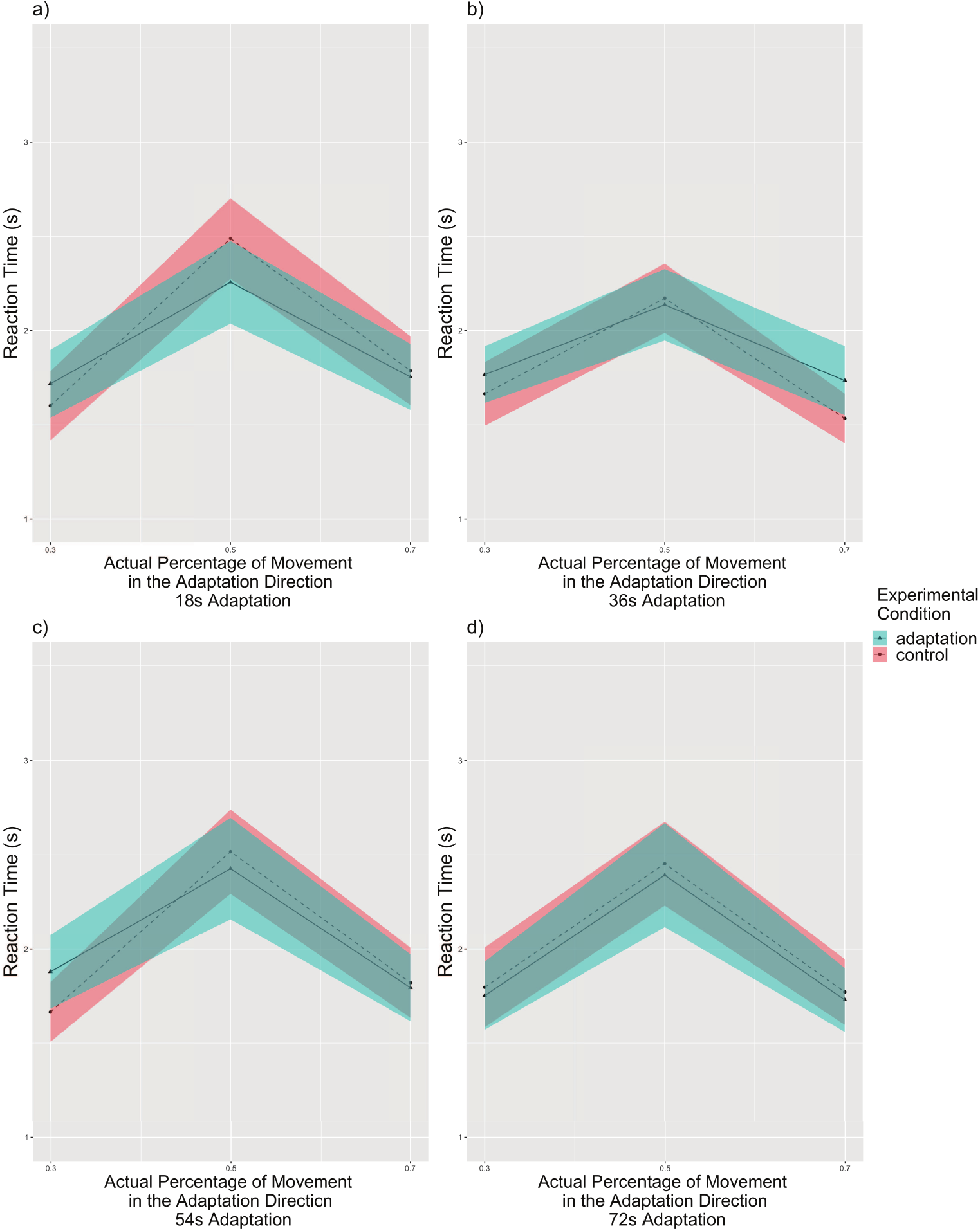
Experiment 3: Reaction times compared with the actual percentage for filtered subjects, separated by adaptation time periods (n = 24). The reaction time increased as the actual percentage approached 50% in all adaptation time periods. a) Reaction times for 18s adaptation trials. b) Reaction times for 36s adaptation trials. c) Reaction times for 54s adaptation trials. d) Reaction times for 72s adaptation trials. Solid lines indicate the grand average value, and the shaded area indicate 1 standard error of means. Tukey correction.

#### Reported Rate by Experimental Conditions

**Appendix 0—figure22.**
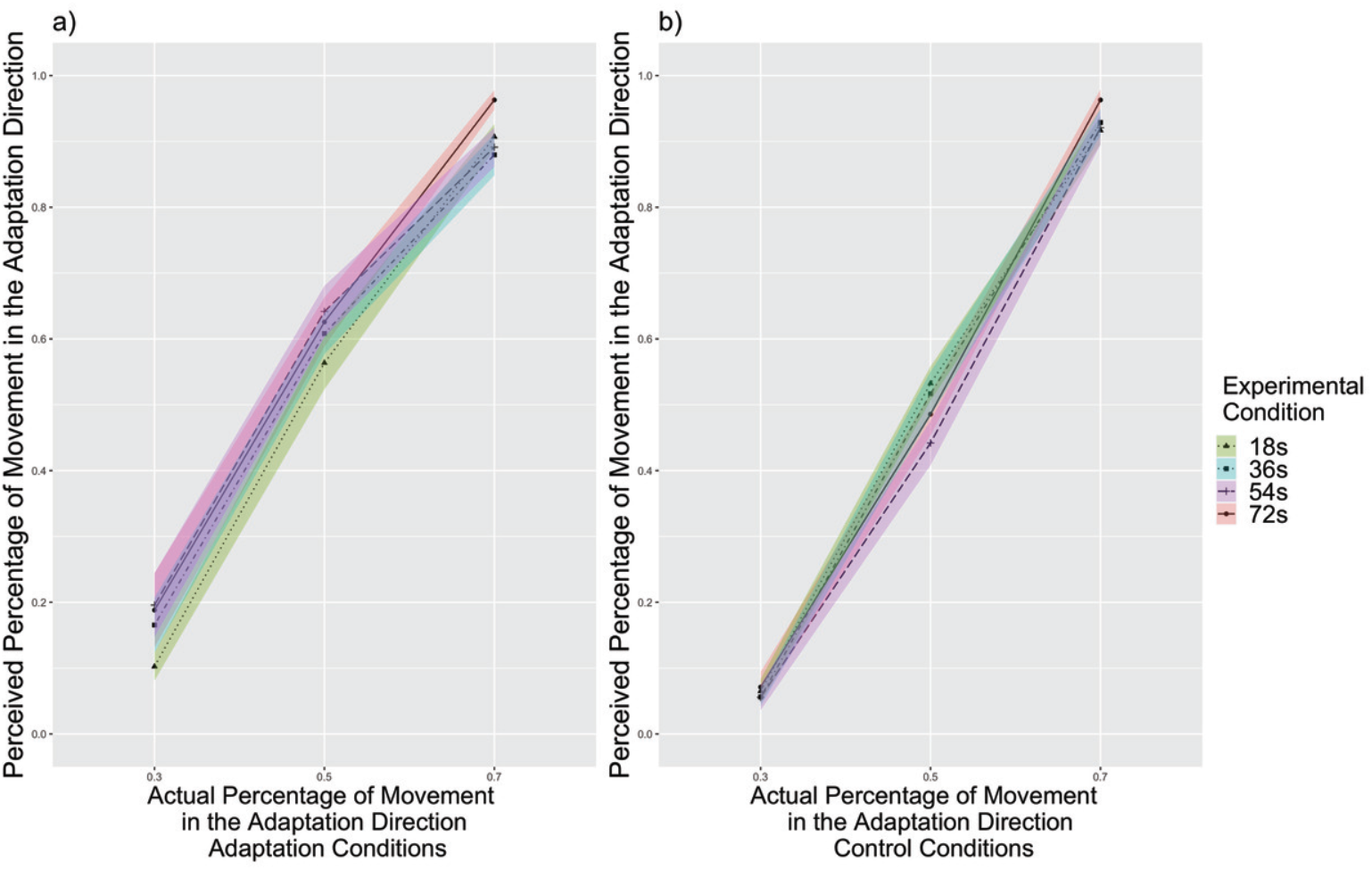
Experiment 3: The perceived percentage of movement in the adaptation direction compared with the actual percentage for filtered subjects, separated by experimental conditions (n = 24). a) The reported rate for adaptation condition. At 70%, 72s adaptation trials had higher perceived percentage than 36s adaptation trials (p = 0.014) and slightly higher than 54s adaptation trials (p = 0.053), supporting that the aftereffect increased with adaptation time. The aftereffect is in the same direction of travel. b) The reported rate for control condition. There were no differences between trials with different adaptation time periods within any actual percentage. Tukey correction.

#### Reaction Times by Experimental Conditions

**Appendix 0—figure23.**
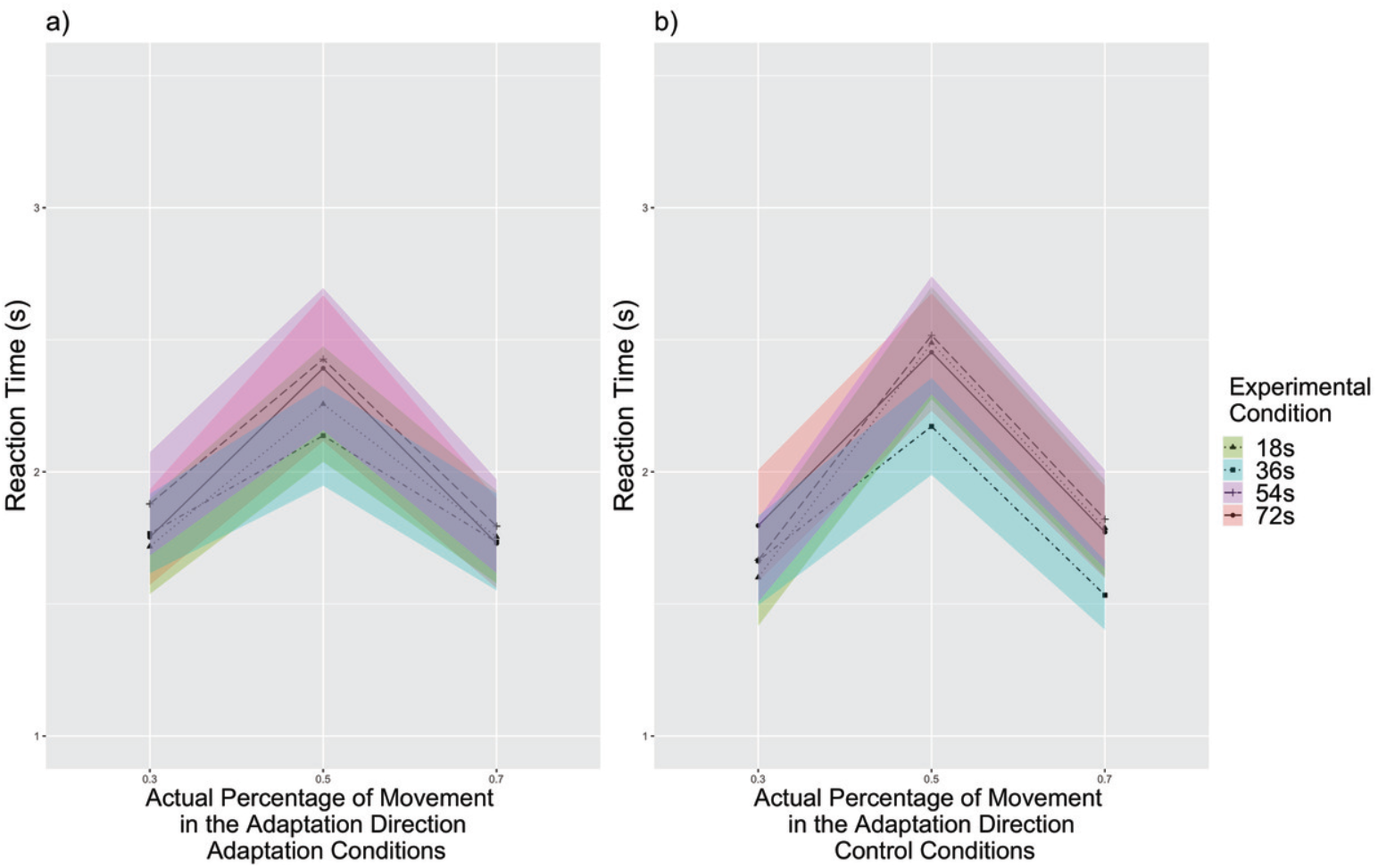
Experiment 3: Reaction times compared with the actual percentage for filtered subjects, separated by experimental conditions (n = 24). The reaction time increased as the actual percentage approached 50% in both experimental conditions. a) Reaction times for adaptation condition. b) Reaction times for control condition. At 30%, people responded slightly slower to 72s adaptation trials than to 18s adaptation trials (p = 0.054). At 70%, people responded slower to 54s adaptation trials than to 36s adaptation trials (p = 0.034). Solid lines indicate the grand average value, and the shaded area indicate 1 standard error of means. Tukey correction.

### Experiment 4: Methods

#### Participants, Stimuli, Task, Design, and Procedure

Similar to Experiment 2 and 3, we calculated a sample size of 24 for within-group comparison in Experiment 4 determined based on power analysis using G*Power software (http://www.gpower.hhu.de/) (***Erdfelder et al., 1996***) based on the weakest effect size from Experiment 1. We recruited 37 participants for Experiment 3, which is more than adequate for the main objectives of this study. All participants were UCI students who participated in return for course credit or monetary incentive ($12/hour). Six participants were discarded for not completing both control and adaptation sessions (*n* = 6). The final pool consisted of 31 participants (15 males, 16 females; 24 not Hispanic or Latino, 5 Hispanic or Latino, 2 not reported; 17 Asian, 6 White, 2 African American, 1 American Indian/Alaskan Native, 4 more than one race, 1 not reported). Ages of the remaining 31 participants ranged from 18 to 31 (mean 21.71). All participants signed an informed consent form in agreement with the UCI Institutional Review Board requirements in accordance with the Declaration of Helsinki.

The Stimuli, Task, Design, and Procedure are the same as Experiment 3. The only change was that in the adaptation session, a few (0 - 3) blocks were inadvertently randomly changed to adaptation to the moon direction. Regardless of whether the block was adapted to the sun or to the moon direction, within each block the initial trial and top-up trials all adapted to the same direction. Participants were not informed of this change in advance. Data from both directions were combined for analysis based on the adaptation direction.

#### Data Analysis

We first removed outlier trials that were 3 standard deviations above or below of the mean of each subject’s reaction time. Approximately 1.88% of trials were removed: 1.70% trials were removed from the experimental session and 2.06% trials were removed from the control session. From the remaining trials for each participant, we calculated the reported percentage of movement toward the adaptation direction as well as mean reaction time for each percentage level at each adaptation condition (the initial adaptation trial in each block were not included). We filtered 4 subjects’ data based on the same criteria from Weibull model fit in Experiment 3. The rest of the data analyses were the same as Experiment.

Same as we have observed in Experiment 1, 2, and 3, people’s report covered three main types of strategies in Experiment 4: counting strategies (*n* = 18), keeping focus on a certain part of the environment for distance estimation (*n* = 4), and a unique strategy (*n* = 9). For the filtered data, there were still subjects using counting (*n* = 15), focusing on a part of the environment (*n* = 4), and a unique strategy (*n* = 8). Again, we controlled for the influence of strategies by adding strategy as a factor in the above ANOVA analyses for reported rate and reaction time.

### Experiment 4: Reaction Times

#### Reaction Times by Adaptation Time Periods

**Appendix 0—figure24.**
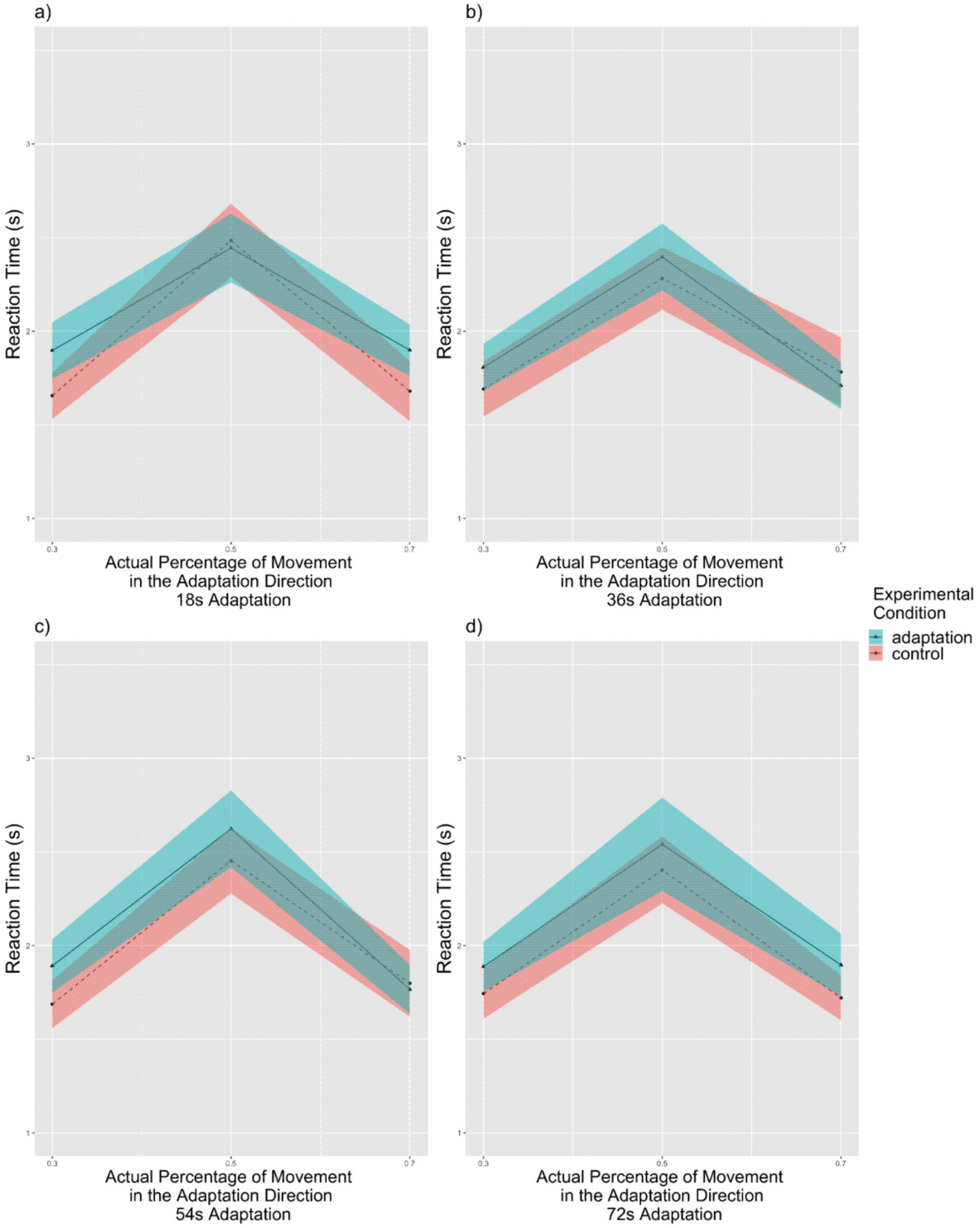
Experiment 4: Reaction times compared with the actual percentage for all subjects, separated by adaptation time periods (n = 31). The reaction time increased as the actual percentage approached 50% in all adaptation time periods. a) Reaction times for 18s adaptation trials. b) Reaction times for 36s adaptation trials. c) Reaction times for 54s adaptation trials. d) Reaction times for 72s adaptation trials. Solid lines indicate the grand average value, and the shaded area indicate 1 standard error of means. Tukey correction.

#### Reaction Times by Experimental Conditions

**Appendix 0—figure25.**
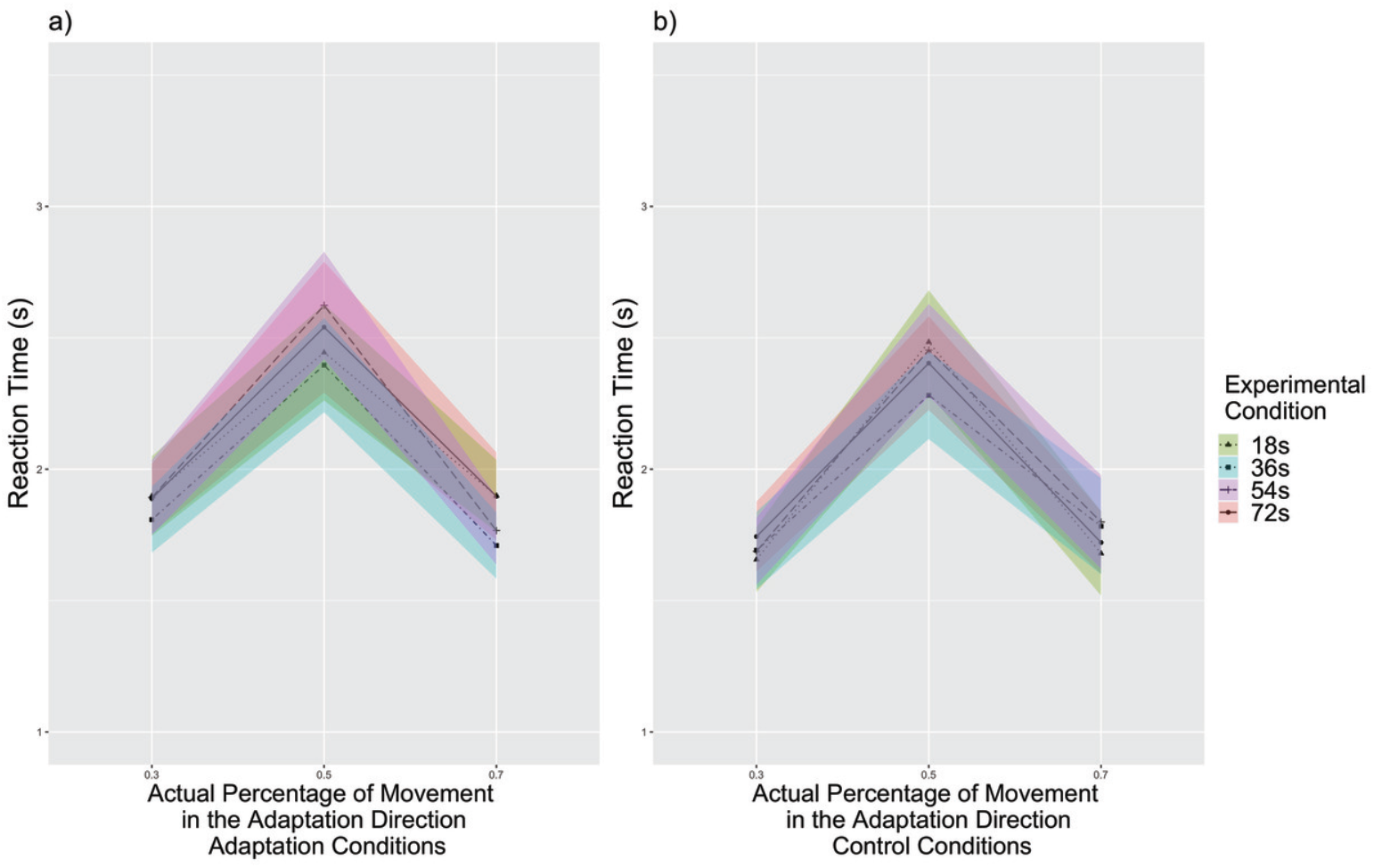
Experiment 4: Reaction times compared with the actual percentage for all subjects, separated by experimental conditions (n = 31). The reaction time increased as the actual percentage approached 50% in both experimental conditions. a) Reaction times for adaptation conditions. b) Reaction times for control conditions. Solid lines indicate the grand average value, and the shaded area indicate 1 standard error of means.

### Experiment 4: Filtered Data

#### Reported Rate by Adaptation Time Periods

**Appendix 0—figure26.**
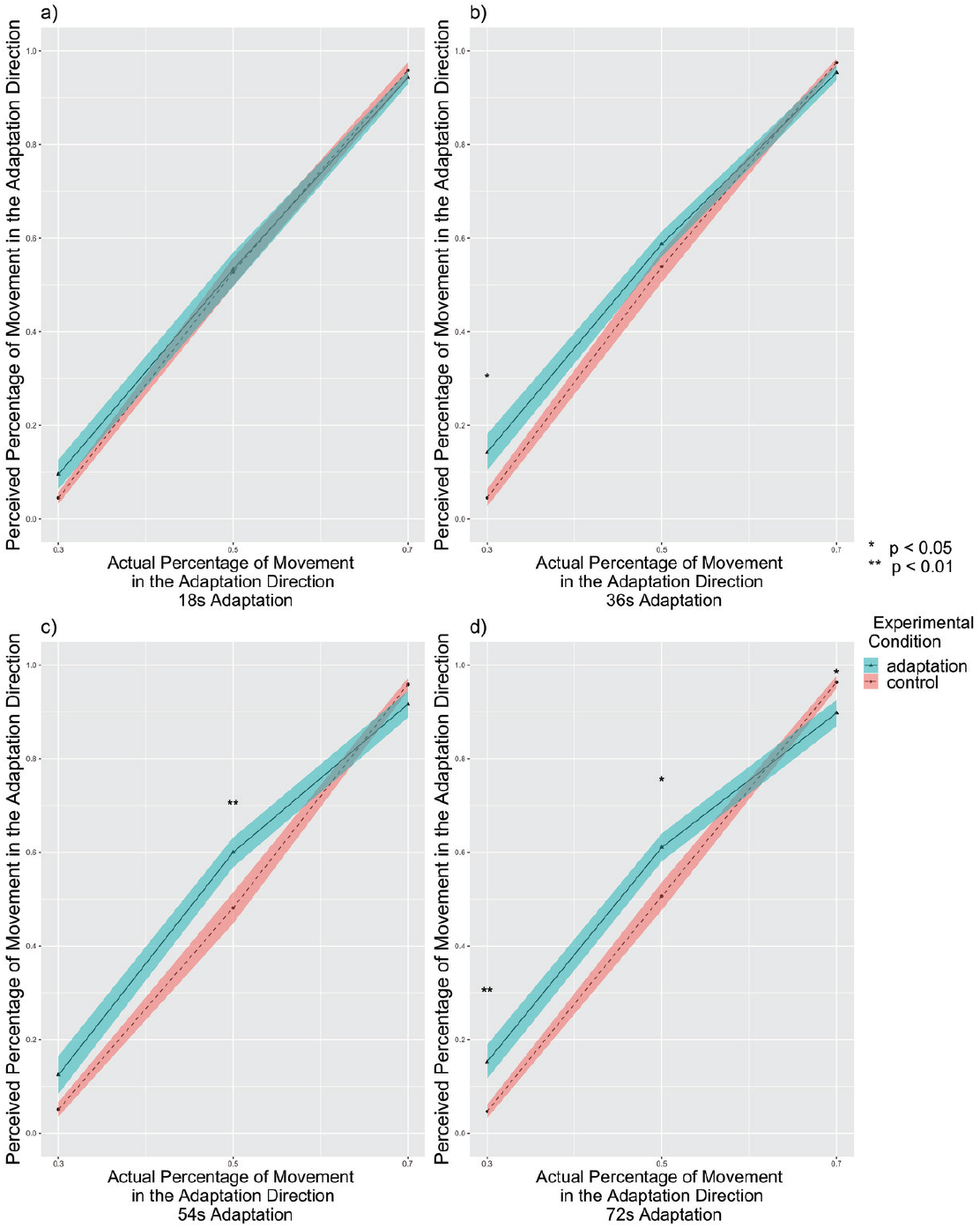
Experiment 4: The perceived percentage of movement in the adaptation direction compared with the actual percentage for filtered subjects, separated by adaptation time periods (n = 27). a) The reported rate for 18s adaptation trials. The adaptation condition showed no difference from the control condition (p = 0.474). b) The reported rate for 36s adaptation trials. The adaptation condition showed higher reported percentages than control conditions (p = 0.029) particularly at 30% (p = 0.012), supporting the aftereffect in the same direction of travel. c) The reported rate for 54s adaptation trials. The adaptation condition showed higher reported percentages than control conditions (p = 0.043) particularly at 50% (p = 0.006), supporting the aftereffect. d) The reported rate for 72s adaptation trials. The adaptation condition showed slightly higher reported percentages than control conditions (p = 0.055) particularly at 30% (p = 0.004) and 50% (p = 0.015), supporting the aftereffect in the same direction of travel. Higher reported percentages was shown in the control condition than in the adaptation condition at 30% (p = 0.043) though. Solid lines indicate the grand average value, and the shaded area indicate 1 standard error of means. * p *<* 0.05, ** p *<* 0.01, Tukey correction.

#### Reaction Times by Adaptation Time Periods

**Appendix 0—figure27.**
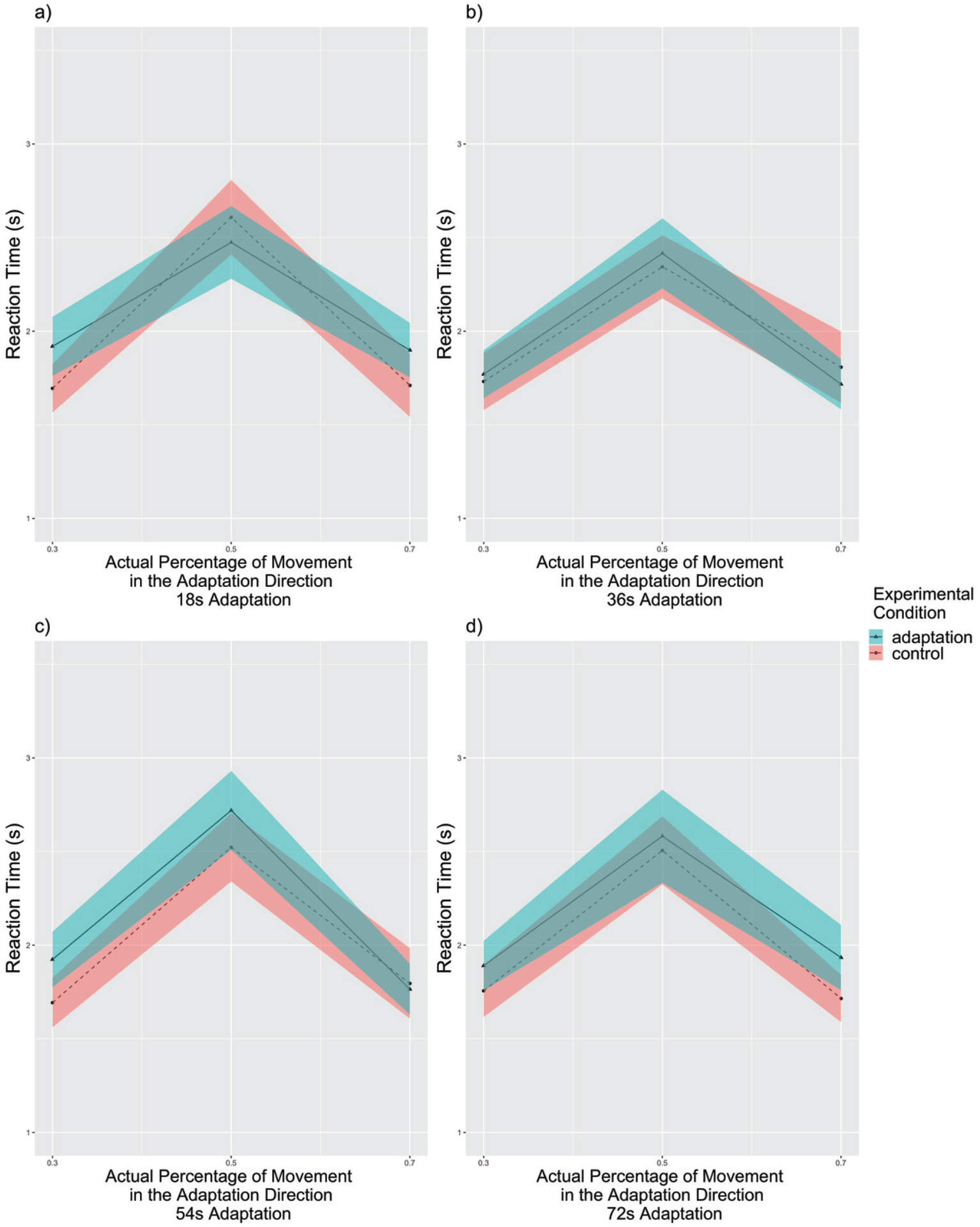
Experiment 4: Reaction times compared with the actual percentage for filtered subjects, separated by adaptation time periods (n = 27). The reaction time increased as the actual percentage approached 50% in all adaptation time periods. a) Reaction times for 18s adaptation trials. b) Reaction times for 36s adaptation trials. c) Reaction times for 54s adaptation trials. d) Reaction times for 72s adaptation trials. Solid lines indicate the grand average value, and the shaded area indicate 1 standard error of means. Tukey correction.

#### Reported Rate by Experimental Conditions

**Appendix 0—figure28.**
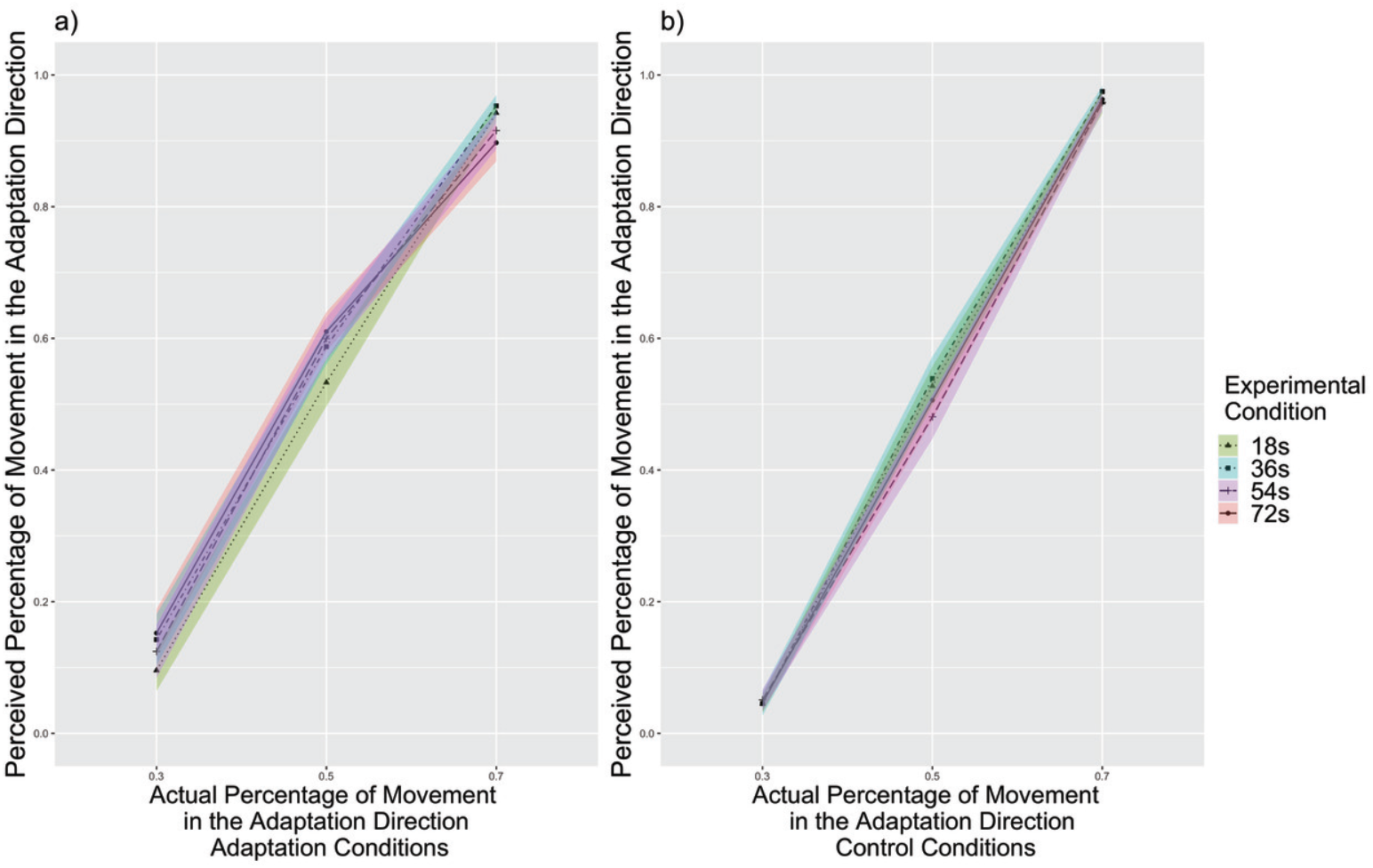
Experiment 4: The perceived percentage of movement in the adaptation direction compared with the actual percentage for filtered subjects, separated by experimental conditions (n = 27). a) The reported rate for adaptation conditions. At 70%, 72s adaptation trials had slightly higher perceived percentage than 36s adaptation trials (p = 0.058), supporting that the aftereffect increased with adaptation time. The aftereffect is in the same direction of travel. b) The reported rate for control conditions. There were no differences between trials with different adaptation time periods within any actual percentage. Tukey correction.

#### Reaction Times by Experimental Conditions

**Appendix 0—figure29.**
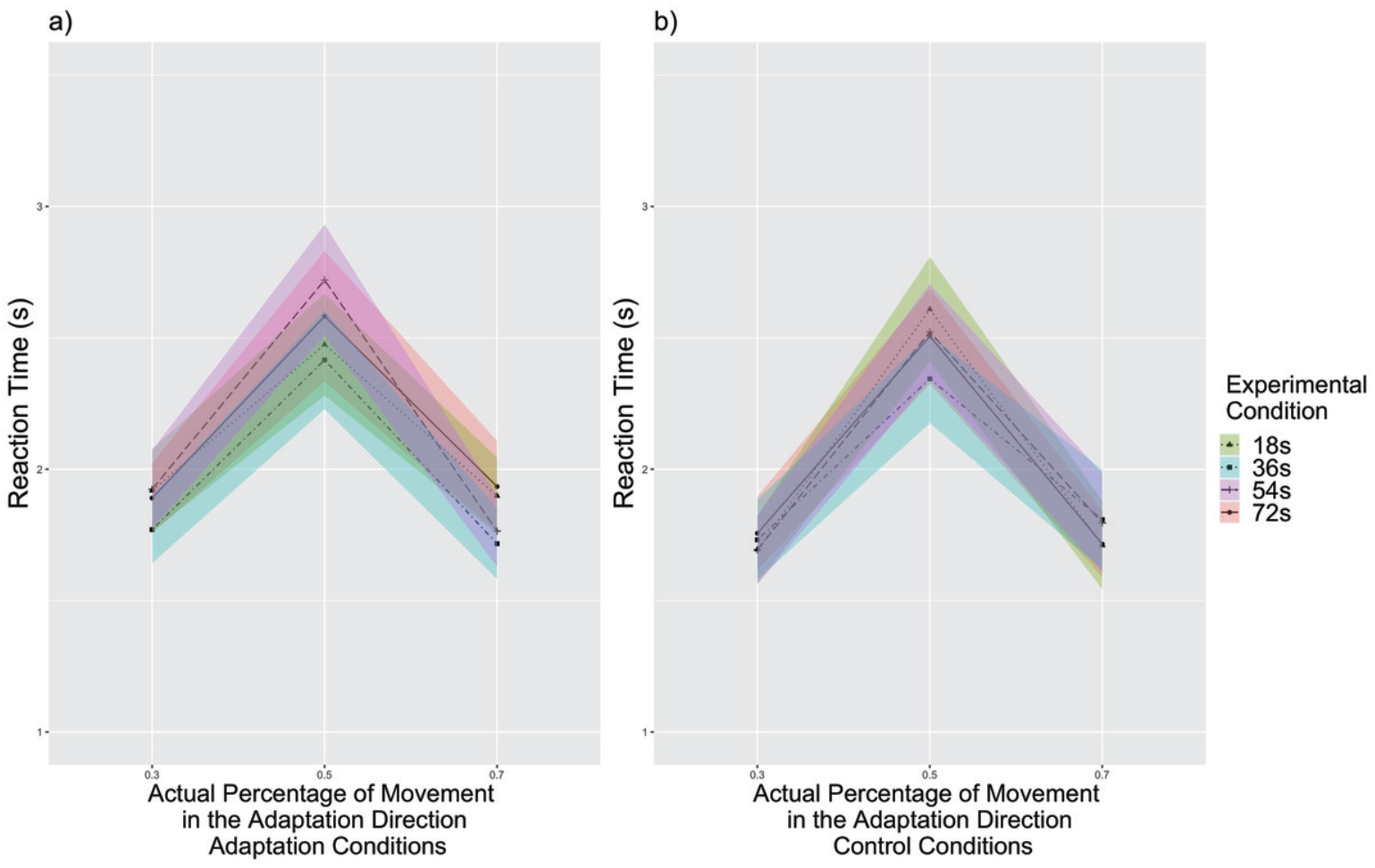
Experiment 4: Reaction times compared with the actual percentage for filtered subjects, separated by experimental conditions (n = 27). The reaction time increased as the actual percentage approached 50% in both experimental conditions. a) Reaction times for adaptation conditions. b) Reaction times for control conditions. Solid lines indicate the grand average value, and the shaded area indicate 1 standard error of means. Tukey correction.

### Experiment 1-4: Dificulty

When comparing difficulty ratings across experiments, there was a main effect of experimental condition that people overall rated adaptation condition to be more difficult than control condition (*F* (1, 145) = 4.57, *p* = 0.034, 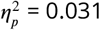, 95%*C.I*. = [0.000, 0.104]), and Experiment 3 was rated more difficult than Experiment 1 (*p* = 0.003) (Supplemental Information Section Experiment 1-4: Difficulty Raw Data Fig. 30, Supplemental Information Section Experiment 1-4: Difficulty Filtered Data Fig. 31).

#### Raw Data

For the task difficulty, we conducted an *experimental condition* (adaptation, control) × *Experiments* (Experiment 1, Experiment 2, Experiment 3, Experiment 4) mixed-subjects repeated measures ANOVA. We found a main effect of the experimental condition (*F* (1, 145) = 4.57, *p* = 0.034, 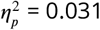, 95%*C.I*. = [0.000, 0.104]) such that subjects found the adaptation condition (*M* = 3.93%±0.12%) to be more difficult than the control condition (*M* = 3.68% ± 0.12%). There was also a main effect of the experiments (*F* (3, 145) = 4.16, *p* = 0.007, 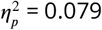, 95%*C.I*. = [0.007, 0.162]). Pairwise t-tests showed people rated Experiment 3 to be more difficult than Experiment 1 (*p* = 0.003) and Experiment 3 to be slightly more difficult than Experiment 4 (*p* = 0.079). There was no interaction between the experimental condition and experiments (*F* (3, 145) = 1.82, *p* = 0.146, 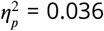, 95%*C.I*. = [0.000, 0.099],). See Figure 30.

Sidak post-hoc analyses revealed that people rated the adaptation session in Experiment 3 to be more difficult than the adaptation session in Experiment 1 (*p* = 0.018). People rated the control session in Experiment 3 to be marginally more difficult than the control session in Experiment 1 (*p* = 0.051). No difference in difficulty ratings were found between adaptation and control conditions within each experiment (all *p >* 0.08).

**Appendix 0—figure30.**
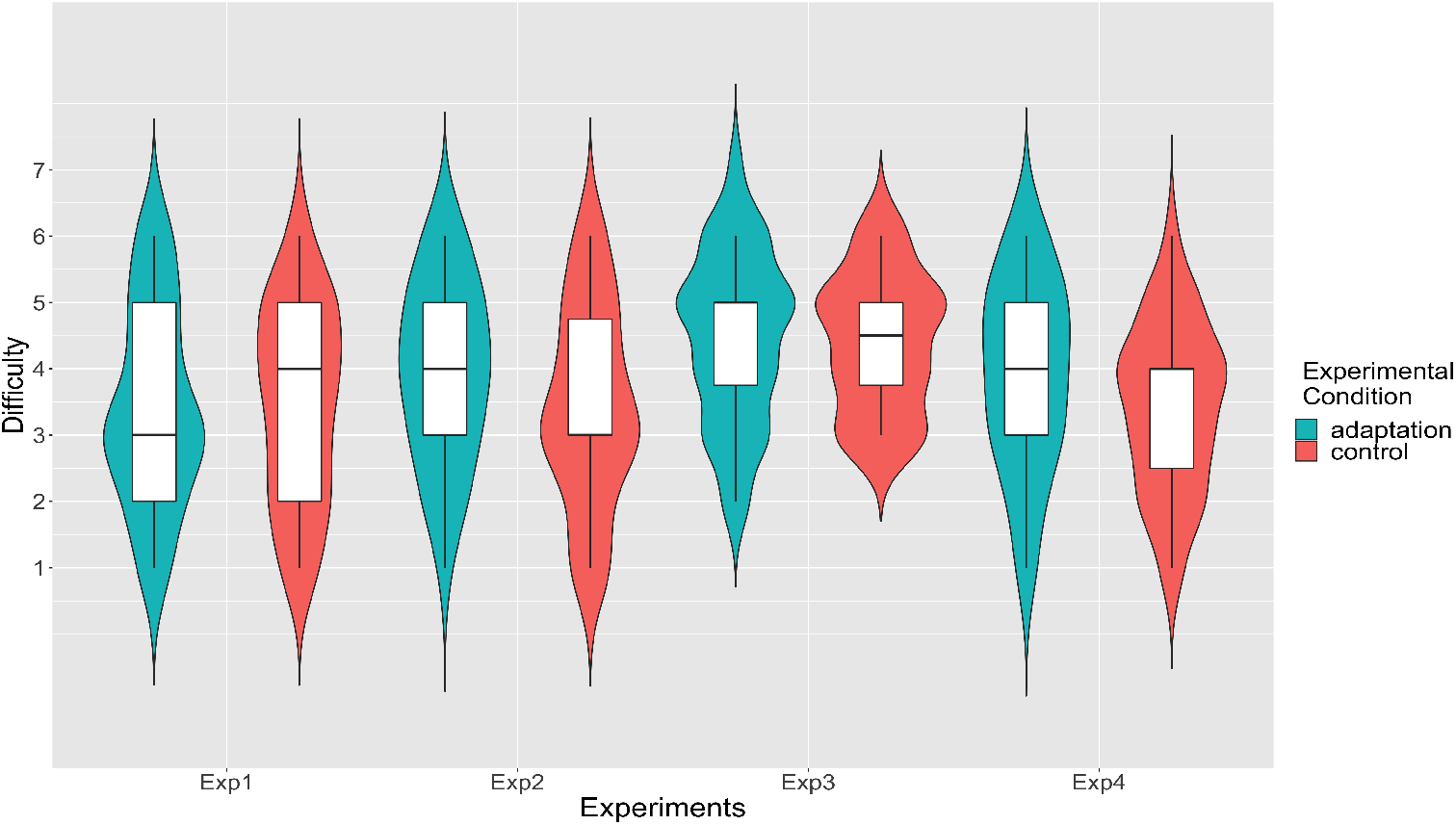
Task difficulty, separated by experiments and adaptation conditions. People reported the adaptation condition to be around the same as the control condition in Experiment 1 (*p* = 0.992), Experiment 2 (*p* = 0.164), Experiment 3 (*p* = 0.998), and Experiment 4 (*p* = 0.176). Note: Experiment 1 (n = 60), Experiment 2 (n = 30), Experiment 3 (n = 28), Experiment 4 (n = 31)

#### Filtered Data

Then we conducted same analyses for task difficulty on filtered data. We found a main effect of the experimental condition (*F* (1, 122) = 4.99, *p* = 0.027, 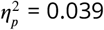, 95%*C.I*. = [0.000, 0.127]) such that subjects found the adaptation condition (*M* = 3.95% ± 0.14%) to be more difficult than the control condition (*M* = 3.67% ± 0.13%). There was also a main effect of the experiments (*F* (3, 122) = 3.31, *p* = 0.023, 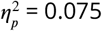, 95%*C.I*. = [0.001, 0.164]). Pairwise t-tests showed people rated Experiment 3 to be more difficult than Experiment 1 (*p* = 0.014). There was a interaction between the experimental condition and experiments (*F* (3, 122) = 2.91, *p* = 0.037, 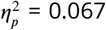, 95%*C.I*. = [0.000, 0.152]). See Figure 31.

Sidak post-hoc analyses revealed that people rated the adaptation session in Experiment 3 to be more difficult than the adaptation session in Experiment 1 (*p* = 0.041). In Experiment 2, people rated the adaptation session to be only marginally more difficult than the control session (*p* = 0.067).

**Appendix 0—figure31.**
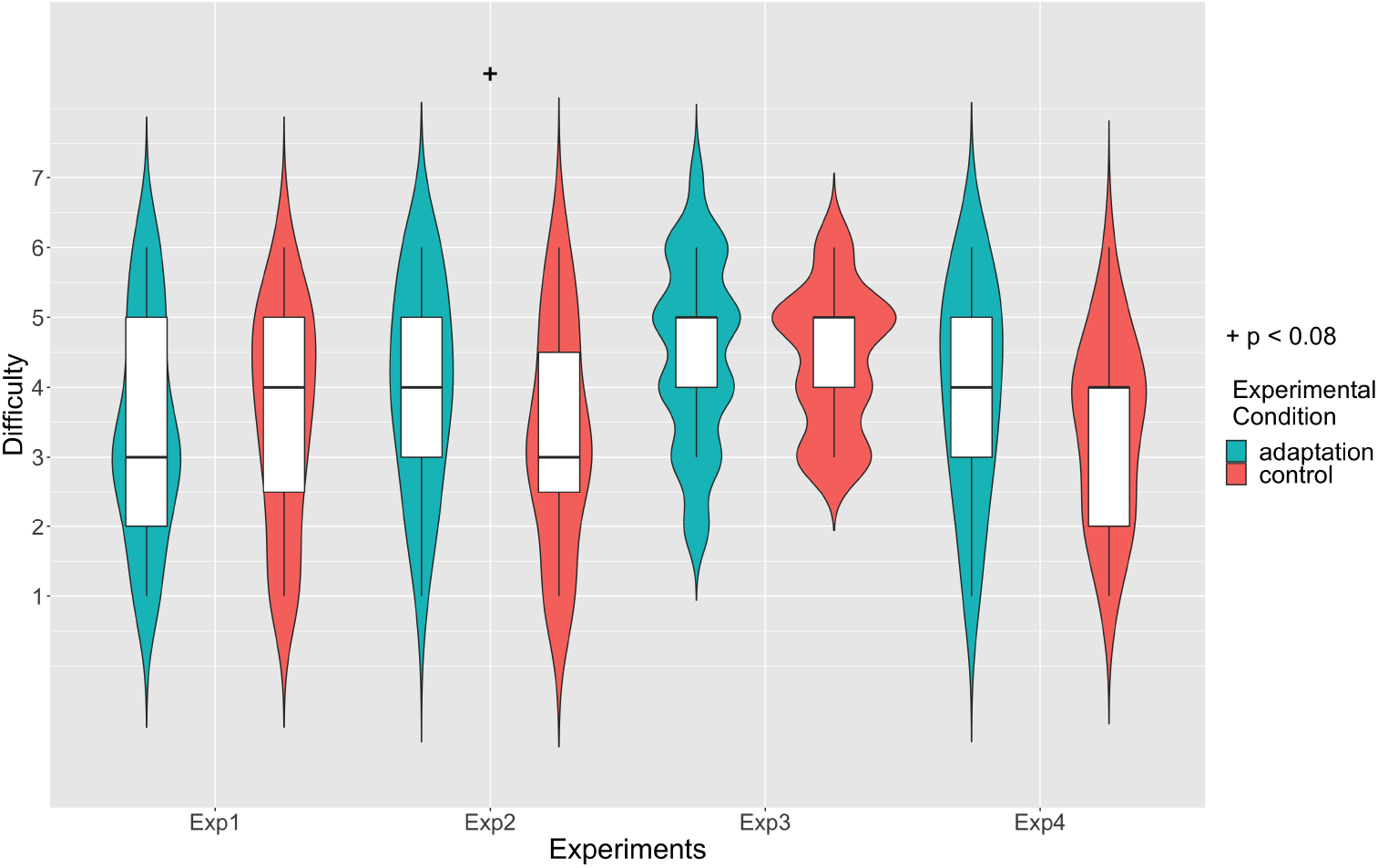
Task difficulty, separated by experiments and adaptation conditions. People reported the adaptation condition to be marginally more difficult than the control condition in Experiment 2 (*p* = 0.067), but not in Experiment 1 (*p* = 0.836), Experiment 3 (*p* = 0.997), or Experiment 4 (*p* = 0.146) (Sidek corrected). Note: Experiment 1 (n = 51), Experiment 2 (n = 23), Experiment 3 (n = 24), Experiment 4 (n = 27)

## Strategies

### Experiment1

#### Reported Rate

##### Raw Data

For the perceived percentage, we conducted an *experimental condition*(adaptation, control) × *actual percentage*(20%, 30%, 40%, 50%, 60%, 70%, 80%) × *strategy*(counting strategies, focusing on part of the environment, unique strategies) mixed-subjects repeated measures ANOVA. There was no main effect of strategy (*F* (2, 53) *<* 0.001, *p* = 0.398, 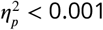, 95%*CI* = [0.000, 0.000], *ns*), condition (*F* (1, 53) = 2.62, *p* = 0.112, 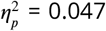, 95%*CI* = [0.000, 0.196], *ns*), or interaction between strategy and condition (*F* (2, 53) = 0.87, *p* = 0.424, 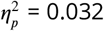, 95%*CI* = [0.000, 0.148], *ns*). As expected, there was also a main effect of the actual percentage (*F* (2.08, 110.39) = 257.10, *p <* 0.001, 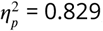, 95%*CI* = [0.799, 0.852]) that showed the perceived percentage increased with the actual percentage. We also found a marginally significant interaction between strategy and the actual percentage (*F* (4.17, 110.39) = 2.34, *p* = 0.057, 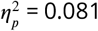, 95%*CI* = [0.007, 0.113]). There was no interaction between condition and the actual percentage (*F* (4.04, 214.16) = 1.47, *p* = 0.213, 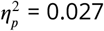, 95%*CI* = [0.000, 0.053], *ns*), or interaction among strategy, condition, and the actual percentage (*F* (8.08, 214.16) = 1.34, *p* = 0.224, 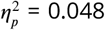, 95%*CI* = [0.000, 0.066],). Mixed-subjects repeated measures ANOVA on the filtered report rate revealed the same pattern of results as the raw data except that there was no interaction between strategy and the actual percentage. For more details, see below.

##### Filtered Data

Then we conducted the same ANOVA analyses on filtered data. There was no main effect of strategy (*F* (2, 44) = 0.02, *p* = 0.977, 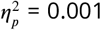, 95%*CI* = [0.000, 0.000], *ns*), condition (*F* (1, 44) = 2.82, *p* = 0.100, 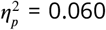, 95%*CI* = [0.000, 0.234], *ns*), or interaction between strategy and condition (*F* (2, 44) = 0.97, *p* = 0.386, 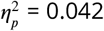, 95%*CI* = [0.000, 0.182], *ns*). As expected, there was also a main effect of the actual percentage (*F* (2.13, 93.83) = 312.39, *p <* 0.001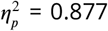, 95%*CI* = [0.852, 0.895]) that showed the perceived percentage increased with the actual percentage. We found no interaction between strategy and the actual percentage (*F* (4.27, 93.83) = 1.38, *p* = 0.244, 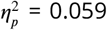, 95%*CI* = [0.000, 0.109]), between condition and the actual percentage (*F* (3.76, 165.34) = 1.03, *p* = 0.393, 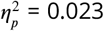, 95%*CI* = [0.000, 0.047], *ns*), or among strategy, condition, and the actual percentage (*F* (7.52, 165.34) = 1.71, *p* = 0.105, 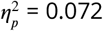, 95%*CI* = [0.000, 0.100], *ns*).

## Reaction Time

There were also no reaction time differences in strategies. For more details, see below.

### Raw Data

For the reaction time, we conducted an *experimental condition*(adaptation, control) × *actual percentage*(20%, 30%, 40%, 50%, 60%, 70%, 80%) × *strategy*(counting strategies, focusing on part of the environment, unique strategies) mixed-subjects repeated measures ANOVA. There was no main effect of strategy (*F* (2, 53) = 1.56, *p* = 0.220, 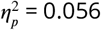, 95%*CI* = [0.000, 0.192], *ns*), condition (*F* (1, 53) = 2.49, *p* = 0.121, 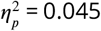, 95%*CI* = [0.000, 0.192], *ns*), or interaction between strategy and condition (*F* (2, 53) = 4.37, *p* = 0.224, 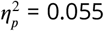, 95%*CI* = [0.000, 0.191], *ns*). As expected, there was a main effect of the actual percentage (*F* (4.23, 223.96) = 7.34, *p <* 0.001, 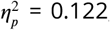, 95%*CI* = [0.051, 0.179]) that showed the perceived percentage increased with the actual percentage. We found no interaction between strategy and the actual percentage (*F* (8.45, 223.96) = 1.32, *p* = 0.229, 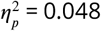, 95%*CI* = [0.000, 0.089], *ns*), between condition and the actual percentage (*F* (3.79, 200.86) = 0.41, *p* = 0.790, 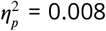, 95%*CI* = [0.000, 0.014],), or interaction among strategy, condition, and the actual percentage (*F* (7.58, 200.86) = 0.66, *p* = 0.715, 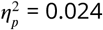, 95%*CI* = [0.000, 0.025], *ns*).

### Filtered Data

Then we conducted the same ANOVA analyses on filtered data. There was no main effect of strategy (*F* (2, 44) = 0.69, *p* = 0.506, 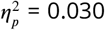, 95%*CI* = [0.000, 0.157], *ns*), condition (*F* (1, 44) = 2.98, *p* = 0.091, 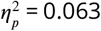, 95%*CI* = [0.000, 0.238], *ns*), or interaction between strategy and condition (*F* (2, 44) = 1.60, *p* = 0.213, 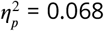, 95%*CI* = [0.000, 0.225], *ns*). As expected, there was a main effect of the actual percentage (*F* (4.19, 184.32) = 7.40, *p <* 0.001, 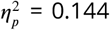, 95%*CI* = [0.061, 0.210]) that showed the perceived percentage increased with the actual percentage. We found no interaction between strategy and the actual percentage (*F* (8.38, 184.32) = 1.37, *p* = 0.208, 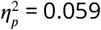, 95%*CI* = [0.000, 0.108],), between condition and the actual percentage (*F* (4.54, 199.65) = 0.31, *p* = 0.891, 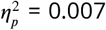, 95%*CI* = [0.000, 0.010], *ns*), or interaction among strategy, condition, and the actual percentage (*F* (9.07, 199.65) = 0.56, *p* = 0.831, 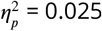, 95%*CI* = [0.000, 0.021], *ns*).

## Experiment 2

### Reported Rate

#### Raw Data

For the perceived percentage, we conducted an *experimental condition*(adaptation, control) × *actual percentage*(20%, 30%, 40%, 50%, 60%, 70%, 80%) × *strategy*(counting strategies, focusing on part of the environment, unique strategies) mixed-subjects repeated measures ANOVA. There was a main effect of strategy (*F* (2, 27) = 8.79, *p* = 0.001, 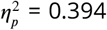, 95%*CI* = [0.097, 0.597]) where people who focused on a certain part of the environment (*M* = 0.578 ± 0.037) reported more toward adaptation direction than people using counting strategies (*M* = 0.496 ± 0.029;*p* = 0.003) or unique strategies (*M* = 0.487 ± 0.029;*p* = 0.002). There was no difference in report rate between people who used counting strategies or unique strategies (*p* = 0.886).

There was also a main effect of condition (*F* (1, 27) = 5.58, *p* = 0.026, 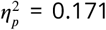, 95%*CI* = [0.000, 0.418]) where people reported more toward the adaptation direction in the adaptation session (*M* = 0.532 ± 0.025) than in the control session (*M* = 0.493 ± 0.026).

There was a marginally significant interaction between strategy and condition (*F* (2, 27) = 3.10, *p* = 0.061, 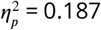, 95%*CI* = [0.000, 0.417]). As expected, there was also a main effect of the actual percentage (*F* (2.12, 57.37) = 214.78, *p <* 0.001, 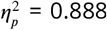, 95%*CI* = [0.859, 0.909]) that showed the perceived percentage increased with the actual percentage. There was no interaction between strategy and the actual percentage (*F* (4.25, 57.37) = 1.33, *p* = 0.269, 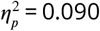, 95%*CI* = [0.000, 0.122], *ns*).

There was a significant interaction between condition and the actual percentage (*F* (3.53, 95.40) = 3.23, *p* = 0.020, 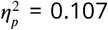, 95%*CI* = [0.012, 0.178]). Tukey HSD analyses revealed that people reported more toward the adaptation direction in the adaptation session than in the condition session at 20% (*p* = 0.003), 40% (*p* = 0.005), 50% (*p* = 0.031), and marginally significant at 30% (*p* = 0.070).

There was a marginally significant interaction among strategy, condition, and the actual percentage (*F* (7.07, 95.40) = 2.10, *p* = 0.050, 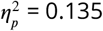, 95%*CI* = [0.003, 0.185]).

The filtered data had similar results; for more details, see below.

#### Filtered Data

Then we conducted the same ANOVA analyses on filtered data. There was a main effect of strategy (*F* (2, 20) = 13.83, *p <* 0.001, 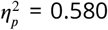, 95%*CI* = [0.241, 0.747]) where people who focused on a certain part of the environment (*M* = 0.582 ± 0.046) reported more toward adaptation direction than people using counting strategies (*M* = 0.497 ± 0.029;*p <* 0.001) or unique strategies (*M* = 0.503 ±0.033;*p <* 0.001). There was no difference in report rate between people who used counting strategies or unique strategies (*p* = 0.910).

There was also a main effect of condition (*F* (1, 27) = 5.58, *p* = 0.026, 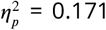, 95%*CI* = [0.000, 0.418]) where people reported more toward the adaptation direction in the adaptation session (*M* = 0.532 ± 0.025) than in the control session (*M* = 0.493 ± 0.026).

There was no interaction between strategy and condition (*F* (2, 20) = 2.03, *p* = 0.158, 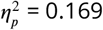, 95%*CI* = [0.000, 0.428], *ns*). As expected, there was also a main effect of the actual percentage (*F* (2.30, 45.93) = 151.55, *p <* 0.001, 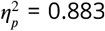, 95%*CI* = [0.847, 0.908]) that showed the perceived percentage increased with the actual percentage. There was no interaction between strategy and the actual percentage (*F* (4.59, 45.93) = 1.24, *p* = 0.307, 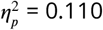, 95%*CI* = [0.000, 0.194], *ns*).

There was a significant interaction between condition and the actual percentage (*F* (3.94, 78.80) = 4.90, *p* = 0.001, 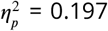, 95%*CI* = [0.056, 0.295]). Tukey HSD analyses revealed that people reported more toward the adaptation direction in the adaptation session than in the condition session at 20% (*p <* 0.001) and 40% (*p* = 0.004), and marginally significant at 50% (*p* = 0.073).

There was a significant interaction among strategy, condition, and the actual percentage (*F* (7.88, 78.80) = 3.21, *p* = 0.003 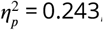, 95%*CI* = [0.060, 0.320]). Tukey HSD analyses revealed that people who used strategy of focusing on part of the environment reported more toward the adaptation direction in the adaptation session than in the condition session at 20% (*p* = 0.012), 40% (*p* = 0.021), and marginally significant at 50% (*p* = 0.072). People who used unique strategies reported more toward the adaptation direction in the adaptation session than in the condition session at 20% (*p* = 0.029). At 40% actual percentage, people who focused on part of the environment reported more toward the adaptation direction in adaptation sessions than people with counting strategies in adaptation sessions (*p* = 0.014) and people with unique strategies in control sessions (*p* = 0.018). At 50% actual percentage, people who focused on part of the environment reported more toward the adaptation direction in adaptation sessions than people with unique strategies in both adaptation sessions (*p* = 0.018) and control sessions (*p* = 0.042), as well as people with counting strategies in both adaptation sessions (*p* = 0.021) and control sessions (*p* = 0.017).

## Reaction Time

There were no reaction time differences in strategies. For more details, see below.

### Raw Data

For the reaction time, we conducted an *experimental condition*(adaptation, control) × *actual percentage*(20%, 30%, 40%, 50%, 60%, 70%, 80%) × *strategy*(counting strategies, focusing on part of the environment, unique strategies) mixed-subjects repeated measures ANOVA. There was no main effect of strategy (*F* (2, 27) = 0.77, *p* = 0.472, 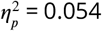, 95%*CI* = [0.000, 0.244], *ns*), condition (*F* (1, 27) = 0.44, *p* = 0.514, 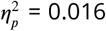, 95%*CI* = [0.000, 0.202], *ns*), or interaction between strategy and condition (*F* (2, 27) = 0.59, *p* = 0.560, 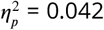, 95%*CI* = [0.000, 0.220], *ns*). As expected, there was a main effect of the actual percentage (*F* (2.61, 70.40) = 14.51, *p <* 0.001, 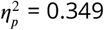, 95%*CI* = [0.221, 0.440]) such that the reaction time increased with actual percentage, peaking at 50%. We found no interaction between strategy and the actual percentage (*F* (5.21, 70.40) = 0.73, *p* = 0.608, 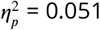, 95%*CI* = [0.000, 0.058], *ns*), between condition and the actual percentage (*F* (2.72, 73.32) = 1.02, *p* = 0.384, 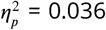, 95%*CI* = [0.000, 0.077], *ns*), or interaction among strategy, condition, and the actual percentage (*F* (5.43, 73.32) = 1.66, *p* = 0.150, 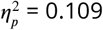, 95%*CI* = [0.000, 0.193], *ns*).

### Filtered Data

Then we conducted the same ANOVA analyses on filtered data. We found a marginally significant main effect of strategy (*F* (2, 20) = 2.68, *p* = 0.093, 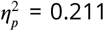, 95%*CI* = [0.000, 0.471]). Tukey HSD analyses did not found difference in reaction time between people focusing on part of the environment (*M* = 1.555 ± 0.096*s*) and people using counting strategies (*M* =2.348 ± 0.072; *p* = 0.089), or people using unique strategies (*M* = 1.986 ± 0.081s, *p* = 0.513). There was also no difference in reaction time between people using counting strategies and people using unique strategies (*p* = 0.444). We found a main effect of experimental condition (*F* (1, 20) = 4.82, *p* = 0.040, 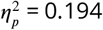, 95%*CI* = [0.000, 0.474]) where people responded slower in the adaptation session (*M* = 2.244 ± 0.077) than in the control session (*M* = 1.956 ± 0.063*a*).

There was no interaction between strategy and condition (*F* (2, 20) = 0.56, *p* = 0.580, 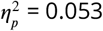, 95%*CI* = [0.000, 0.271], *ns*). As expected, there was a main effect of the actual percentage (*F* (2.43, 48.62) = 7.14, *p* = 0.001, 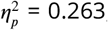, 95%*CI* = [0.112, 0.366]) such that the reaction time increased with actual percentage, peaking at 50%. We found no interaction between strategy and the actual percentage (*F* (4.86, 48.62) = 0.56, *p* = 0.729, 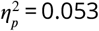, 95%*CI* = [0.000, 0.046], *ns*), between condition and the actual percentage (*F* (3.40, 68.05) = 0.90, *p* = 0.457, 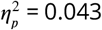, 95%*CI* = [0.000, 0.088],), or interaction among strategy, condition, and the actual percentage (*F* (6.81, 68.05) = 1.16, *p* = 0.337, 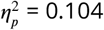, 95%*CI* = [0.000, 0.185], *ns*).

### Experiment 3

#### Reported Rate

##### Raw Data

For the perceived percentage, we conducted an *experimental condition*(adaptation, control) × *actual percentage*(30%, 50%, 70%) × *adaptation time periods* (18s, 36s, 54s, 72s) × *strategy*(counting strategies, focusing on part of the environment, unique strategies) mixed-subjects repeated measures ANOVA. There was a main effect of condition (*F* (1, 25) = 4.69, *p* = 0.040, 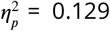, 95%*CI* = [0.000, 0.383]) where people reported more toward the adaptation direction in the adaptation session (*M* = 0.566 ± 0.020) than in the control session (*M* = 0.486 ± 0.021). As expected, there was also a main effect of the actual percentage (*F* (1.37, 34.18) = 298.65, *p <* 0.001, 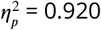, 95%*CI* = [0.849, 0.951]) that showed the perceived percentage increased with the actual percentage. There was a marginally significant interaction between the experimental condition and the actual percentage (*F* (1.65, 41.22) = 2.99, *p* = 0.070, 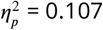, 95%*CI* = [0.000, 0.270]). Because the interaction was only marginally significant, we did not conduct further post-hoc analyses.

There were no main effect of strategy (*F* (2, 25) = 0.06, *p* = 0.946 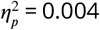, 95%*CI* = [0.000, 0.060],), no main effect of adaptation time periods (*F* (2.68, 66.89) = 0.72, *p* = 0.529, 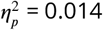, 95%*CI* = [0.000, 0.056], *ns*), no interaction between the strategy and the experimental condition (*F* (2, 25) = 0.33, *p* = 0.725, 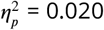, 95%*CI* = [0.000, 0.167],), no interaction between the strategy and the actual percentage (*F* (2.73, 34.18) = 2.50, *p* = 0.081, 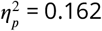, 95%*CI* = [0.000, 0.356], *ns*), no interaction between the strategy and adaptation time periods (*F* (5.35, 66.89) = 0.64, *p* = 0.684, 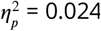, 95%*CI* = [0.000, 0.000], *ns*), no interaction between the actual percentage and adaptation time periods (*F* (4.41, 110.33) = 1.33, *p* = 0.269, 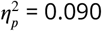, 95%*CI* = [0.000, 0.122], *ns*), no interaction between the experimental condition and adaptation time periods (*F* (4.25, 57.37) = 0.66, *p* = 0.637, 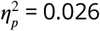, 95%*CI* = [0.000, 0.053], *ns*), no interaction among the strategy, the experimental condition, and the actual percentage (*F* (3.30, 41.22) = 2.10, *p* = 0.050, 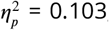, 95%*CI* = [0.000, 0.234], *ns*), no interaction among the strategy, the experimental condition, and adaptation time periods (*F* (4.77, 59.61) = 0.73, *p* = 0.601, 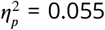, 95%*CI* = [0.000, 0.112], *ns*), no interaction among the strategy, the actual percentage, and adaptation time periods (*F* (8.83, 110.33) = 0.82, *p* = 0.594, 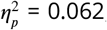, 95%*CI* = [0.000, 0.074], *ns*), no interaction among the experimental condition, the actual percentage, and adaptation time periods (*F* (4.13, 103.23) = 1.45, *p* = 0.223, 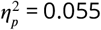, 95%*CI* = [0.000, 0.106], *ns*), or interaction among the strategy, the experimental condition, the actual percentage, and adaptation time periods (*F* (8.26, 103.23) = 1.18, *p* = 0.318, 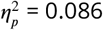, 95%*CI* = [0.000, 0.155], *ns*).

##### Filtered Data

For the perceived percentage, we conducted an *experimental condition*(adaptation, control) × *actual percentage*(30%, 50%, 70%) × *adaptation time periods* (18s, 36s, 54s, 72s) × *strategy*(counting strategies, focusing on part of the environment, unique strategies) mixed-subjects repeated measures ANOVA. There was a main effect of condition (*F* (1, 21) = 4.66, *p* = 0.043, 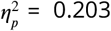, 95%*CI* = [0.000, 0.476]) where people reported more toward the adaptation direction in the adaptation session (*M* = 0.561 ± 0.021) than in the control session (*M* = 0.496 ± 0.022). As expected, there was also a main effect of the actual percentage (*F* (1.32, 27.71) = 281.68, *p <* 0.001, 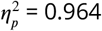, 95%*CI* = [0.927, 0.979]) that showed the perceived percentage increased with the actual percentage.

There was a significant interaction between the experimental condition and the actual percentage (*F* (1.98, 41.56) = 4.57, *p* = 0.016, 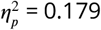, 95%*CI* = [0.007, 0.368]). Tukey HSD post-hoc analyses revealed that people had increased report rate in the adaptation condition than the control condition at 50% actual percentage (*p* = 0.011).

There were no main effect of strategy (*F* (2, 21) = 0.14, *p* = 0.866 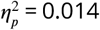, 95%*CI* = [0.000, 0.147], *ns*), no main effect of adaptation time periods (*F* (2.63, 55.16) = 0.82, *p* = 0.474, 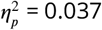, 95%*CI* = [0.000, 0.169], *ns*), no interaction between the strategy and the experimental condition (*F* (2, 21) = 0.01, *p* = 0.993, 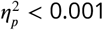, 95%*CI* = [0.000, 0.000], *ns*), no interaction between the strategy and the actual percentage (*F* (2.64, 27.71) = 1.58, *p* = 0.220, 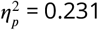, 95%*CI* = [0.000, 0.449], *ns*), no interaction between the strategy and adaptation time periods (*F* (5.25, 55.16) = 0.77, *p* = 0.578 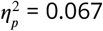, 95%*CI* = [0.000, 0.069],), no interaction between the actual percentage and adaptation time periods (*F* (4.78, 100.36) = 0.71, *p* = 0.614, 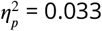, 95%*CI* = [0.000, 0.067], *ns*), no interaction between the experimental condition and adaptation time periods (*F* (2.51, 52.76) = 1.84, *p* = 0.160, 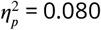, 95%*CI* = [0.000, 0.204], *ns*), no interaction among the strategy, the experimental condition, and the actual percentage (*F* (3.96, 41.56) = 2.00, *p* = 0.113 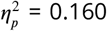, 95%*CI* = [0.000, 0.320], *ns*), no interaction among the strategy, the experimental condition, and adaptation time periods (*F* (5.02, 52.76) = 0.61, *p* = 0.690, 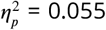, 95%*CI* = [0.000, 0.112], *ns*), no interaction among the strategy, the actual percentage, and adaptation time periods (*F* (9.56, 100.36) = 0.90, *p* = 0.535, 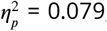, 95%*CI* = [0.000, 0.098], *ns*), no interaction among the experimental condition, the actual percentage, and adaptation time periods (*F* (3.81, 79.98) = 1.80, *p* = 0.140, 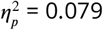, 95%*CI* = [0.000, 0.146], *ns*), or interaction among the strategy, the experimental condition, the actual percentage, and adaptation time periods (*F* (7.62, 79.98) = 1.20, *p* = 0.311, 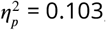, 95%*CI* = [0.000, 0.137], *ns*).

## Reaction Time

There were also no reaction time differences in strategies. For more details, see below.

### Raw Data

For the reaction time, we conducted an *experimental condition*(adaptation, control) × *actual percentage*(30%, 50%, 70%) × *adaptation time periods* (18s, 36s, 54s, 72s) × *strategy*(counting strategies, focusing on part of the environment, unique strategies) mixed-subjects repeated measures ANOVA. We only observed a main effect of the actual percentage (*F* (1.29, 32.36) = 37.87, *p <* 0.001, 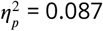, 95%*CI* = [0.000, 0.305]) such that the reaction time increased with actual percentage, peaking at 50%.

There were no effect of condition (*F* (1, 25) = 0.26, *p* = 0.612, 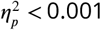, 95%*CI* = [0.000, 0.113], *ns*), no main effect of strategy (*F* (2, 25) = 0.81, *p* = 0.455 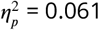, 95%*CI* = [0.000, 0.264], *ns*), no main effect of adaptation time periods (*F* (2.55, 63.65) = 1.52, *p* = 0.223, 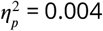, 95%*CI* = [0.000, 0.000], *ns*), no interaction between the experimental condition and the actual percentage (*F* (1.79, 44.70) = 0.98, *p* = 0.374, 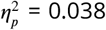, 95%*CI* = [0.000, 0.164], *ns*), no interaction between the strategy and the experimental condition (*F* (2, 25) = 0.11, *p* = 0.896, 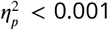, 95%*CI* = [0.000, 0.000], *ns*), no interaction between the strategy and the actual percentage (*F* (2.59, 32.36) = 2.25, *p* = 0.109, 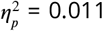, 95%*CI* = [0.000, 0.000], *ns*), no interaction between the strategy and adaptation time periods (*F* (5.09, 63.65) = 0.46, *p* = 0.809, 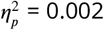, 95%*CI* = [0.000, 0.000], *ns*), no interaction between the actual percentage and adaptation time periods (*F* (4.18, 104.47) = 0.74, *p* = 0.573, 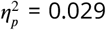, 95%*CI* = [0.000, 0.060], *ns*), no interaction between the experimental condition and adaptation time periods (*F* (2.71, 67.72) = 0.46, *p* = 0.691, 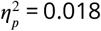, 95%*CI* = [0.000, 0.079], *ns*), no interaction among the strategy, the experimental condition, and the actual percentage (*F* (3.58, 44.70) = 0.75, *p* = 0.551, 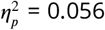, 95%*CI* = [0.000, 0.157], *ns*), no interaction among the strategy, the experimental condition, and adaptation time periods (*F* (5.42, 67.72) = 1.11, *p* = 0.367, 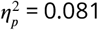, 95%*CI* = [0.000, 0.159], *ns*), no interaction among the strategy, the actual percentage, and adaptation time periods (*F* (8.36, 104.47) = 0.59, *p* = 0.795, 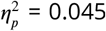, 95%*CI* = [0.000, 0.041], *ns*), no interaction among the experimental condition, the actual percentage, and adaptation time periods (*F* (3.80, 94.98) = 0.73, *p* = 0.568, 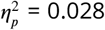, 95%*CI* = [0.000, 0.059], *ns*), or interaction among the strategy, the experimental condition, the actual percentage, and adaptation time periods (*F* (7.60, 94.98) = 0.41, *p* = 0.907, 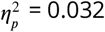, 95%*CI* = [0.000, 0.010], *ns*).

### Filtered Data

Then we conducted the same ANOVA analyses on filtered data. Again, we only observed a main effect of the actual percentage (*F* (1.22, 25.52) = 36.42, *p <* 0.001, 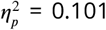, 95%*CI* = [0.000, 0.343]) such that the reaction time increased with actual percentage, peaking at 50%. There were no effect of condition (*F* (1, 21) = 0.15, *p* = 0.705, 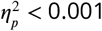, 95%*CI* = [0.000, 0.108], *ns*), no main effect of strategy (*F* (2, 21) = 0.43, *p* = 0.655, 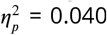, 95%*CI* = [0.000, 0.238], *ns*), no main effect of adaptation time periods (*F* (2.53, 53.17) = 1.80, *p* = 0.166, 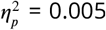, 95%*CI* = [0.000, 0.000], *ns*), no interaction between the experimental condition and the actual percentage (*F* (1.71, 35.88) = 0.58, *p* = 0.539, 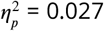, 95%*CI* = [0.000, 0.152], *ns*), no interaction between the strategy and the experimental condition (*F* (2, 21) = 0.24, *p* = 0.791, 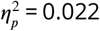, 95%*CI* = [0.000, 0.000], *ns*), no interaction between the strategy and the actual percentage (*F* (2.43, 25.52) = 2.19, *p* = 0.124, 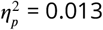, 95%*CI* = [0.000, 0.000], *ns*), no interaction between the strategy and adaptation time periods (*F* (5.06, 53.17) = 0.52, *p* = 0.761, 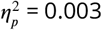, 95%*CI* = [0.000, 0.000], *ns*), no interaction between the actual percentage and adaptation time periods (*F* (4.19, 88.07) = 1.30, *p* = 0.275, 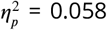, 95%*CI* = [0.000, 0.114], *ns*), no interaction between the experimental condition and adaptation time periods (*F* (2.75, 57.79) = 0.36, *p* = 0.768, 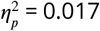, 95%*CI* = [0.000, 0.194], *ns*), no interaction among the strategy, the experimental condition, and the actual percentage (*F* (3.42, 35.88) = 0.58, *p* = 0.653, 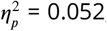, 95%*CI* = [0.000, 0.153], *ns*), no interaction among the strategy, the experimental condition, and adaptation time periods (*F* (5.50, 57.79) = 1.18, *p* = 0.328, 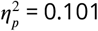, 95%*CI* = [0.000, 0.194], *ns*), no interaction among the strategy, the actual percentage, and adaptation time periods (*F* (8.39, 88.07) = 1.01, *p* = 0.438, 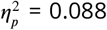, 95%*CI* = [0.000, 0.158], *ns*), no interaction among the experimental condition, the actual percentage, and adaptation time periods (*F* (3.69, 77.59) = 0.74, *p* = 0.558, 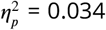, 95%*CI* = [0.000, 0.070], *ns*), or interaction among the strategy, the experimental condition, the actual percentage, and adaptation time periods (*F* (7.39, 77.59) = 0.62, *p* = 0.750, 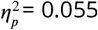, 95%*CI* = [0.000, 0.054], *ns*).

## Experiment 4

### Reported Rate

### Raw Data

For the perceived percentage, we conducted an *experimental condition*(adaptation, control) × *actual percentage*(30%, 50%, 70%) × *adaptation time periods* (18s, 36s, 54s, 72s) × *strategy*(counting strategies, focusing on part of the environment, unique strategies) mixed-subjects repeated measures ANOVA. There was a main effect of condition (*F* (1, 28) = 5.23, *p* = 0.030, 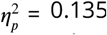, 95%*CI* = [0.000, 0.376]) where people reported more toward the adaptation direction in the adaptation session (*M* = 0.555 ± 0.019) than in the control session (*M* = 0.503 ± 0.020). As expected, there was also a main effect of the actual percentage (*F* (1.46, 40.91) = 570.20, *p <* 0.001, 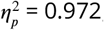, 95%*CI* = [0.948, 0.982]) that showed the perceived percentage increased with the actual percentage. There was a significant interaction between the experimental condition and the actual percentage (*F* (1.57, 43.96) = 3.87, *p* = 0.038, 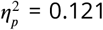, 95%*CI* = [0.000, 0.279]). Post-hoc Tukey analyses revealed that adaptation increased perceived percentage at 30% (*p* = 0.038) and 50% (*p* = 0.016). All other main effects and interactions were not significant (all *p >* 0.08).

### Filtered Data

For the perceived percentage, we conducted an *experimental condition*(adaptation, control) × *actual percentage*(30%, 50%, 70%) × *adaptation time periods* (18s, 36s, 54s, 72s) × *strat-egy*(counting strategies, focusing on part of the environment, unique strategies) mixed-subjects repeated measures ANOVA. There was a main effect of condition (*F* (1, 24) = 4.93, *p* = 0.036, 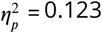, 95%*CI* = [0.000, 0.381]) where people reported more toward the adaptation direction in the adaptation session (*M* = 0.542 ± 0.018) than in the control session (*M* = 0.503 ± 0.006). As expected, there was also a main effect of the actual percentage (*F* (1.49, 35.67) = 579.28, *p <* 0.001, 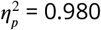, 95%*CI* = [0.962, 0.988]) that showed the perceived percentage increased with the actual percentage. There was a significant interaction between the experimental condition and the actual percentage (*F* (1.67, 40.17) = 4.89, *p* = 0.017, 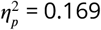 95%*CI* = [0.010, 0.346]). Tukey HSD post-hoc analyses revealed that people had increased report rate in the adaptation condition than the control condition at 50% (*p* = 0.032) and 50% (*p* = 0.019). All other main effects and interactions were not significant (all *p >* 0.08).

## Reaction Time

### Raw Data

For the reaction time, we conducted an *experimental condition*(adaptation, control) × *actual percentage*(30%, 50%, 70%) × *adaptation time periods* (18s, 36s, 54s, 72s) × *strategy*(counting strategies, focusing on part of the environment, unique strategies) mixed-subjects repeated measures ANOVA. We observed a main effect of the actual percentage (*F* (1.14, 31.92) = 34.57, *p <* 0.001, 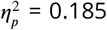, 95%*CI* = [0.000, 0.411]) such that the reaction time increased with actual percentage, peaking at 50%. There was a significant interaction between the strategy and the experimental condition (*F* (2, 28) = 3.79, *p* = 0.035, 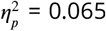, 95%*CI* = [0.000, 0.260]). Post-hoc Tukey paired t-tests did not found difference between any pairs of strategies and conditions. There was also a marginally significant interaction among the experimental condition, the actual percentage, and adaptation time periods (*F* (4.57, 128.02) = 2.11, *p* = 0.074, 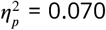, 95%*CI* = [0.000, 0.128]). Because this interaction was only marginally significant, we did not conduct further posthoc analyses. All other main effects and interactions were not significant (all *p >* 0.08).

### Filtered Data

Then we conducted the same ANOVA analyses on filtered data. Again, we observed a main effect of the actual percentage (*F* (1.13, 27.04) = 33.88, *p <* 0.001, 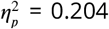, 95%*CI* = [0.000, 0.446]) such that the reaction time increased with actual percentage, peaking at 50%. There was a significant interaction between the strategy and the experimental condition (*F* (2, 24) = 3.95, *p* = 0.033, 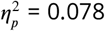, 95%*CI* = [0.000, 0.296]). Post-hoc Tukey paired t-tests found that people with the strategy of focusing on part of the environment responded slower in the adaptation condition than the control condition (*p* = 0.024). There was also a marginally significant interaction among the experimental condition, the actual percentage, and adaptation time periods (*F* (4.59, 110.13) = 2.26, *p* = 0.059, 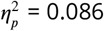, 95%*CI* = [0.000, 0.153]). Because this interaction was only marginally significant, we did not conduct further posthoc analyses. All other main effects and interactions were not significant (all *p >* 0.08).

